# Teasing apart the joint effect of demography and natural selection in the birth of a contact zone

**DOI:** 10.1101/2022.01.11.475794

**Authors:** Lili Li, Pascal Milesi, Mathieu Tiret, Jun Chen, Janek Sendrowski, John Baison, Zhiqiang Chen, Linghua Zhou, Bo Karlsson, Mats Berlin, Johan Westin, Rosario Garcia-Gil, Harry Wu, Martin Lascoux

**Affiliations:** Program in Plant Ecology and Evolution, Department of Ecology and Genetics, EBC and SciLife Lab, Uppsala University; Umeå Plant Science Centre, Department Forest Genetics and Plant Physiology, Swedish University of Agricultural Sciences, SE-90183 Umeå, Sweden; Skogforsk, Ekebo, 2250 SE-268 90 Svalöv, Sweden; Skogforsk, Uppsala Science Park, 751 83 Uppsala, Sweden; Unit for field-based forest research, Swedish University of Agricultural Sciences, SE-922 91 Vindeln, Sweden; College of Life Sciences, Zhejiang University, Hangzhou, Zhejiang 310058, China; CSIRO National Collection Research Australia, Black Mountain Laboratory, Canberra, ACT 2601, Australia

**Keywords:** Local adaptation, contact zone, Last Glacial Maximum, natural selection, demography

## Abstract

Vast population movements induced by recurrent climatic cycles have shaped the genetic structure of plant species. This is especially true in Scandinavia that was repeatedly glaciated. During glacial periods trees were confined to refugia, south and east of the ice sheet, from which they recolonized Scandinavia as the ice melted away. This multi-pronged recolonization led to large contact zones in most species. We leverage large genomic data from 5000 trees to reconstruct the demographic history of Norway spruce (*Picea abies*) and test for the presence of natural selection during the recolonization process and the establishment of the contact zone. Sweden is today made up of two large genetic clusters, a southern one originating from the Baltics and a Northern one originating from Northern Russia. The contact zone delineating these two clusters closely matches the limit between two major climatic regions. This suggests that natural selection contributed to the establishment and the maintenance of the contact zone. To test this hypothesis we first used Approximate Bayesian Computation; an Isolation-with migration model with genome-wide linked selection fits the data better than a purely neutral one. Secondly, we identified loci characterized by both extreme allele frequency differences between geographic regions and association to the variables defining the climatic zones. These loci, many of which are related to phenology, form clusters present on all linkage groups. Altogether, the current genetic structure reflects the joint effect of climatic cycles, recolonization and selection on the establishment of strong local adaptation and contact zones.

**Significance Statement:** Understanding how past climatic events, human actions and evolutionary forces contributed to the present distribution of genetic diversity is crucial to predict their reaction to the current climate crisis. Vast distribution shifts induced by past environmental changes, local ecological processes, natural selection and human transfers contributed to the current distribution of Norway spruce across Northern Europe. Genome-wide polymorphisms from thousands of individuals show that Scandinavia was recolonized after the Last Glacial from both south and north. This two-pronged recolonization established a contact zone between two genetic clusters that matches the limit between two major climate zones. The contact zone is shaped and maintained by natural selection on a large number of loci that form blocks of co-adapted loci spread genome-wide.

## Introduction

All natural populations are structured to varying degrees and more than ever population genetic structure matters (1). It matters for a large range of issues: population structure is intrinsically related to speciation and local adaptation; it conditions the response of species to environmental changes (e.g. climate change or the occurrence of new diseases) and it severely limits our ability to associate genetic polymorphism to phenotypic variation or environmental factors using genome-wide approaches. Population structure, and more generally demography, also hampers efforts to detect genomic signatures of natural selection and recent studies on human height have shown that rather fine-scale structure, if not accounted for, can lead to wrong inferences on past selection (1,2). There is therefore a strong incentive to develop methods to capture fine-scaled population genetic structure and thereby strengthens inferences on the relative parts played by past demography and selection in the evolution of species. So, how far have we gone in this respect? Undoubtedly, the availability of genome-wide polymorphisms has vastly enhanced our ability to describe population genetic structure. Yet capturing fine-scale population genetic structure and making sense of it still remains a major challenge. In part, the difficulty arises from the fact that genetic diversity is distributed both discretely and continuously (3,4). This dual nature of population genetic structure reflects the plurality of processes that shape population structure: vast and complex population movements in response to past climatic changes, gene flow among populations, and sometimes even between species, local adaptation and, in many plant and animal species, human-mediated individual transfers.

All the aforementioned factors have come into play in the history of many plant species. Here we shall focus on one of the most common boreal species, Norway spruce (*Picea abies* (L.) H. Karst)) and mor specifically on Scandinavian populations. Since the seminal study of Lagercrantz and Ryman (5), a large number of studies have outlined the most salient features of the demographic history of Norway spruce (6-16). Basically, as already found by (5), current populations emerged from three main glacial refugia located in the Alps, in the Carpathians and in the Russian plains. This is, of course, a very rough outline and further studies have added more than one twist to it. In particular, recent studies indicated that these main lineages did not evolve independently but instead created many contact zones (8). The nature of these contact zones remains to be elucidated: they could simply be a reflection of past distribution shifts, correspond to ecological zones and be associated to local adaptation, or be the result of both processes. At a larger phylogenetic scale, Siberian spruce (*Picea obovata*) has a major influence in the northern range of *P. abies* with a large introgression zone starting in the Urals and extending quite far westwards (8,9,14). Introgression from *P. obovata* into *P. abies* is lopsided with a much larger contribution at high latitudes (∼65°N and above) than at intermediate ones (∼60°N) (8). The structure of the vast hybrid zone between the two species could, at least in part, be due to differences in ecological requirements between the two parental species (17). Second, in more restricted geographical areas, recent introductions have also contributed significantly to the genetic composition of local populations. The Swedish breeding program was established by selecting trees with superior phenotypes (aka “plus trees”) in natural stands across the whole country. Targeted genome sequencing of the individuals composing the southern part of the breeding program and of individuals sampled across the natural range of *P. abies* revealed that a large proportion of these “plus trees” were recent introductions originating from most parts of the natural range (8). Third, the pattern of differentiation at genotypic, phenotypic, and environmental variables at the site of origin of the trees were highly correlated indicating a strong pattern of local adaptation (18), as often observed in forest trees (19). For example, in both Norway and Siberian spruce, Chen *et al*. (7,20) detected strong latitudinal clines in growth cessation and were able to associate those to a major candidate gene for photoperiodic response, FTL2, and its pattern of expression. Because these populations are recent these results suggest that local adaptation can be established very rapidly. These initial studies were based on a handful of candidate genes but more recent studies relying on a much larger number of markers (e.g.,18,21) suggest that quantitative traits and phenology related traits have a polygenic inheritance with loci involved in local adaptation distributed across the genome. It is, however, still unclear whether adaptive genes are randomly distributed or clustered in some specific regions of the genome. The latter would, for instance, be expected when two populations that are under stabilizing selection for different optima are linked by gene flow as is often the case in forest trees (22,23).

In the present study we sequenced all individuals from the base population of the Swedish *P. abies* breeding program using exome capture (4769 individuals, >500,000 SNPs), generating an unprecedented large and dense sampling along a latitudinal gradient ranging from ∼55°N to ∼67°N. It allowed us to analyze population genetic structure with a very high resolution. More specifically, we were able to test for pattern of isolation-by-distance, identify barriers to gene flow and test whether those reflect physical or environmental barriers or simply historical contingencies. We show that Swedish populations of Norway spruce are divided into two main genetic clusters that closely match the two main climatic regions of the country. Coalescent simulations and Approximate Bayesian Computation allowed the rejection of a purely neutral divergence model between the two main clusters. Furthermore, genome scans indicate that clusters of loci distributed across the 12 linkage groups correspond to areas of high genetic differentiation and are associated to environmental variables. The current distribution of genetic diversity in Norway spruce across Sweden therefore appears to be the result of both demographic processes and local adaptation.

## Results

We first investigated global population structure using the whole dataset which comprises 4607 trees from the base population of the Norway spruce breeding program and 162 trees collected across the natural range of *P. abies* and *P. obovata* (Fig. 1A and Supplementary file 1). Both UMAP (24) and ADMIXTURE (25) retrieved three main domains, Boreal, Carpathians and the Alps, and clusters resulting from admixture between these three domains: Central Europe, Russian-Baltics and Northern Poland (Fig. 1B, C and S1 to S2 in SI Appendix). Sweden is itself divided into two main genetics clusters, one including southern and central Sweden (CSE) and the other one the northern part of the country (NFE) (Fig. 1 A-C). Many trees in southern Sweden also correspond to recent introductions. All this is concordant with (8) and (21). Despite their current geographical closeness, the CSE and NFE clusters are divergent and CSE is more closely related to the Russia-Baltics cluster than to NFE (*F*_*ST*_ = 0.009 and 0.018, respectively, SI Appendix Tab. S1). In addition, the large discrepancy in ancestry components found between two putatively hybrid Russian populations located at the same longitude but at different latitudes support a larger contribution of *P. obovata* to NFE cluster than to CSE (Fig. 1B and C). This general pattern is consistent with a recolonization of the Scandinavian peninsula from refugia with different genetic components and through two different routes, a Northern one and a Southern one. To study more finely the genetic structure of the contact zone and identify the evolutionary forces that shaped it, we focused in the rest of the study on the subset of trees that were native to Sweden and belonged to the CSE (N = 974) and the NFE (N = 784) clusters.

**Figure 1:**
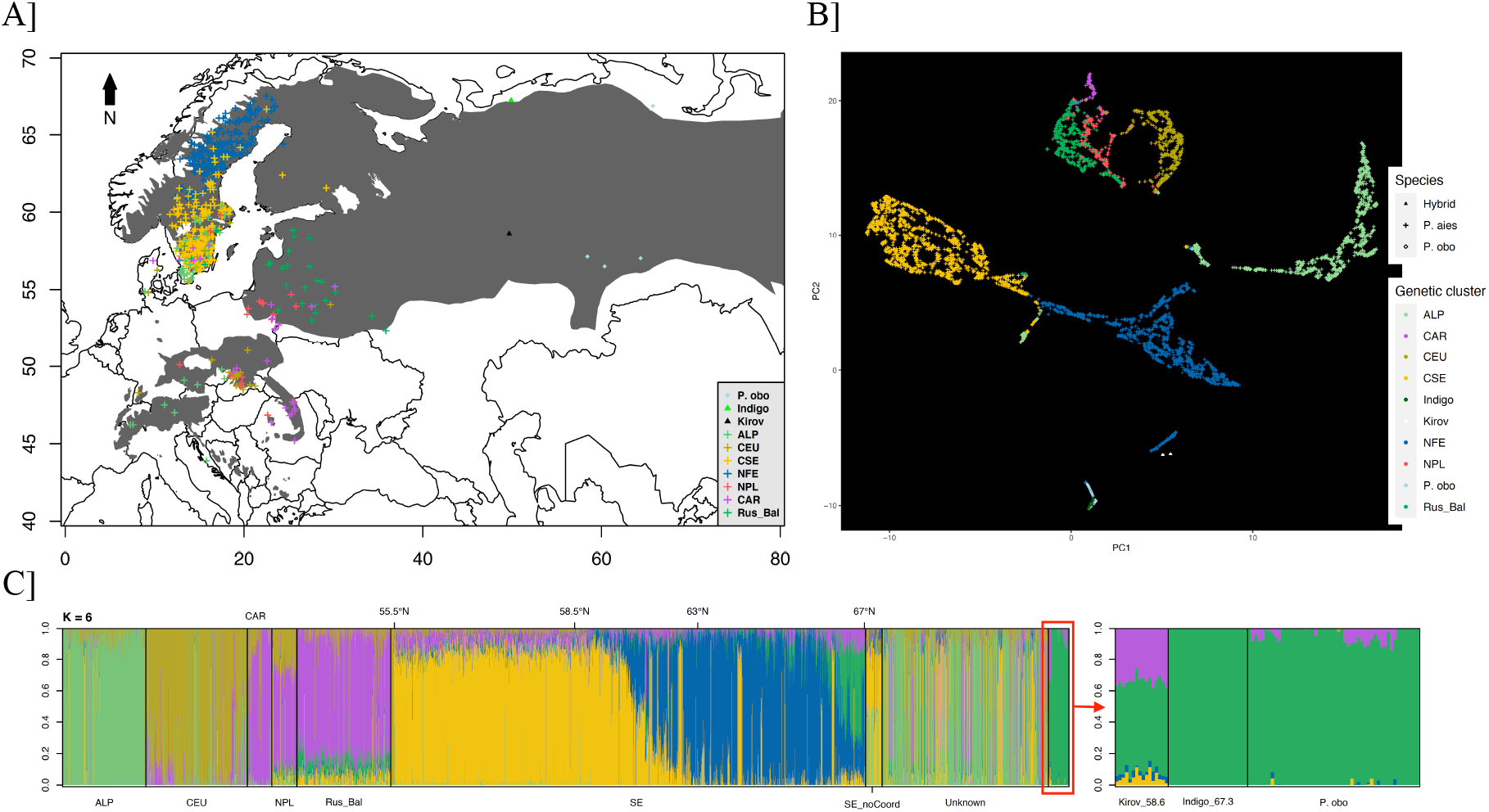
Population genetic structure of *P. abies*. **A)** Sampling location of *P. abies* (plus signs: light green, Alpine (ALP); brown, Central Europe (CEU); purple Carpathian (CAR); red, northern Poland (NPL); dark green Russia-Baltics (Rus_Bal); yellow, southern Fennoscandia (CSE) and dark blue, northern Fennoscandia (NFE), *P. obovata* (diamonds, light blue) and hybrids (Indigo and Kirov) (black triangles). The shaded area corresponds to the distribution range of *P*.*abies* and *P. obovata* **B)** UMAP bi-dimensional plots, colors are the same as for panel A. **C)** Admixture plot for K= 6. Samples from a same geographic origin were grouped. Swedish samples were ordered by latitude. Colors represent different ancestry components.

Both UMAP and ADMIXTURE implicitly aim to detect discrete genetic clusters. However, Norway spruce tends to be continuously distributed and population structure is the result of both isolation-by-distance and discontinuities. To account for this and identify barriers to gene flow, we first used the software *conStruct* (3) that considers different levels of population genetic structure: layers correspond to clusters in ADMIXTURE but isolation by distance is considered within layers. Independently of the number of layers considered, a model including isolation-by-distance within layers predicts the genetic variation pattern better than a non-spatial model (SI Appendix Fig. S3). The lowest cross-validation error (five-fold) was found for three layers (SI Appendix Fig. S3) but, in line with the ADMIXTURE results, two ancestry components explained most of the genetic variation and distinguished southern trees from northern ones (Fig. 1C). The contact zone between these two main clusters occurred between 60°N and 63°N (Fig. 2A). Contributions from the southern cluster into the northern one can be detected at latitudes as high as 66°N while the northern cluster barely contributed to the populations outside of the contact zone. Finally, populations from high latitudes (close to 67°N) also presented a specific ancestry component (Fig. 2A). Based on ADMIXTURE results this ancestry component probably represents more recent introgression from *P. obovata* into the northernmost *P. abies* populations (Fig. 1C). To visualize the variation in effective migration rate across Sweden and detect barriers to gene flow we then fitted the data to a model of isolation-by-distance and estimated effective migration surfaces (EEMS, (26)). The resulting pattern is complex but regions with low effective migration rate (brown areas), correspond to the contact zone already detected by *conStruct* and to mountainous regions in the north. North-South barriers, as the one along the west coast, are likely artifacts dues to the difficulty of EEMS to account for anisotropy (26) (Fig. 2B). To compare gene flow along latitude and longitude we quantified the IBD by regressing a function of pairwise *F*_*ST*_, (*F*_*ST*_ / (1 - *F*_*ST*_)), over the logarithm of distance between populations. According to (27), the inverse of the slope of the regression provides an indirect estimate of dispersal. First, considering all pairs of populations we detected a pattern of IBD, with an estimated dispersal of 209 ± 33 individuals (SI Appendix Fig. S4). But, the IBD was much more pronounced along a latitudinal gradient (228 ± 33), than along a longitudinal gradient (680 ± 187).

**Figure 2:**
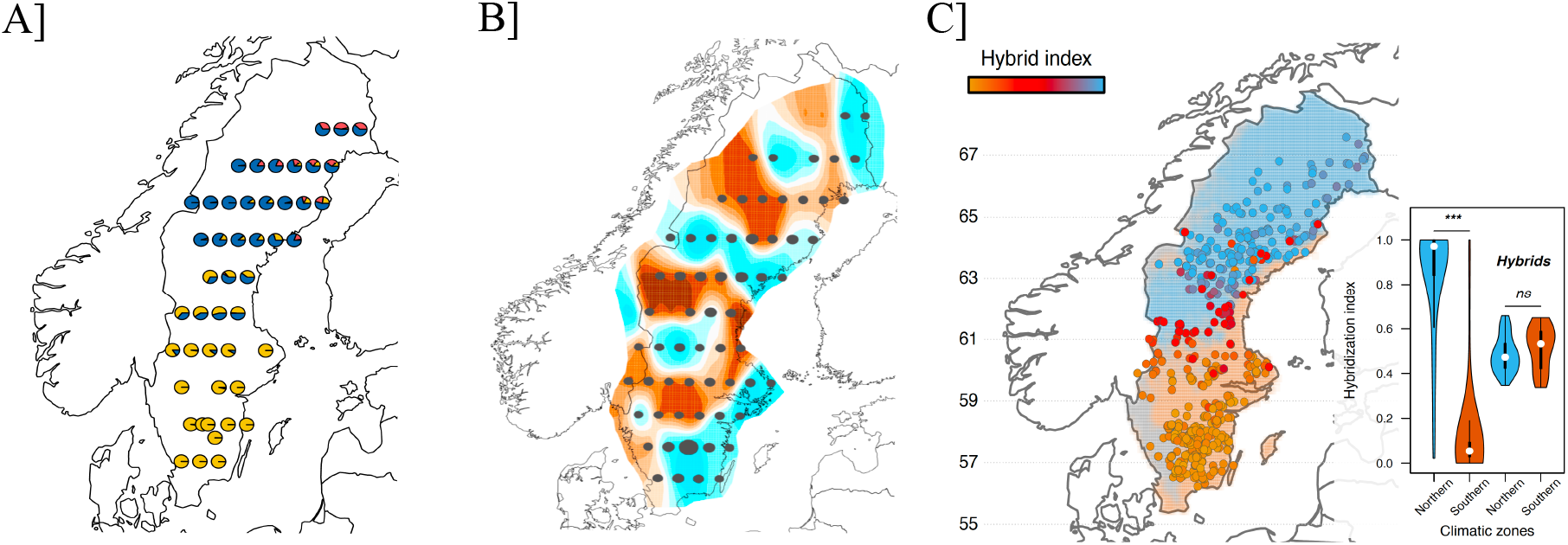
Fine genetic structure of the contact zone and relation to climate zones. **A)** Admixture proportions based on the best spatial model using *conStruct* (K = 3). Colors represent different ancestry components. Close-by samples were grouped into “populations”. **B)** Estimated effective migration surfaces (EEMS). Blue and brown areas respectively indicate regions with a higher or a lower effective migration rate than expected under a model of isolation by distance (IBD). Gray dots represent individual aggregations. **C)** The genetic contact zone overlaps with the transition between the two main climatic zones, the southern one (orange background) and the northern one (blue background). Dots represent tree locations and the color scale corresponds to the hybridization level (from 0, full CSE, orange, to 1, full NFE, blue). Violin plots represent the distribution of hybrid index within each of the two main climatic zones (all samples or only samples with 0.33 > hybrid index < 0.66; ^ns^, *p* > 0.05; ^***^, *p* < 0.001).

To investigate whether ecological barriers to gene flow contributed to the establishment of the contact zone we analyzed environmental variation across Sweden. Three climatic zones were delineated (Fig. 2C, see Online methods § *“Abiotic environment characterization and climatic zones definition”*): the two main ones separate the northern part from the southern part of the country and the differentiation is mainly explained by temperature-related variables (annual mean temperature, minimum or average temperature of the coldest months, seasonality). The third climatic zone corresponds to the mountainous area and the west coast and is characterized by higher precipitations than the two other climatic zones (Fig. 2C and SI Appendix S5).

The genetic contact zone between the Northern (NFE) and the Southern (CSE) clusters almost perfectly overlap the transition between the Northern and Southern climatic zones (Fig. 2C). Based on ancestry components from ADMIXTURE (K = 6) we computed a hybridization index, *h*_*i*_, which varied from 0, (full CSE), to one, (full NFE). While most of the CSE (*h*_*i*_ ≤ 0.33) or NFE (*h*_*i*_ ≥ 0.66) trees were restricted to the Southern or the Northern climatic zone, respectively (Wilcoxon’s rank-sum test, *W* > 7×10^−5^; *p* < 0.001), the hybrids (0.33 < *h*_*i*_ < 0.66) were located on the transition zone and evenly distributed between the two climatic zones (*W* = 789; *p* = 0.18, Fig. 2C). Such a match between the main environmental zones and the genetic structure strongly suggests that natural selection contributed to the creation and maintenance of the contact zone between the two genetics clusters.

To test whether natural selection contributed to the establishment and maintenance of the contact zone we simulated different coalescent isolation with migration scenarios and calculated their posterior probabilities with an Approximate Bayesian Computation (ABC) approach implemented in the program *DILS* (28). Briefly, in the presence of linked selection one expects a larger variance in effective population size, N_e_, among loci than under a strictly isolation with migration model. In order to measure the effect of hybridization on demographic scenario inferences, we created three samples of 20 individuals (10 from NFE and 10 from CSE) varying in their distance to the contact zone (far, intermediate, or close). In all three cases the most likely model was one with linked selection, with posterior probabilities of 71.24%, 93.39%, and 87.94% for far, intermediate and close, respectively (Tab. 1). This suggests that linked selection occurred over the entire range of each climate zone. This result was further confirmed by additional forward simulations (SI Appendix Section 2).

To identify genomic signatures of local adaptation associated with the contact zone, we then (i) scanned our genomic data for loci with extreme allele frequency differences between geographic regions using *Bayenv2* (*X*^*T*^*X* score) (29) and *pcAdapt* (30) and (ii) ran genotype-environment associations (GEA) using *Bayenv2* and *lfmm2* (31) on a subset of 142,765 SNPs with MAF > 0.05. *Bayenv2* is population based while *pcAdapt* and *lfmm2* are individual-based. Genome scans identified 440 and 990 SNPs showing extreme allele frequency differences between geographic regions, using *pcAdapt* or *X*^*T*^*X* statistic, respectively (32 % overlap at the gene level). With GEA, a total of 1616 (*bayenv2*) and 1298 (*flmm2*) SNPs were associated to at least one of the 24 bioclimatic variables (21% overlap at the gene level). The number of significant associations per bioclimatic variable was correlated between the two analyses (Spearman’s rho = 0.53, S = 1070, *p* < 0.01) (SI Appendix Tab. S2). Most of the significant associations were with the climatic variables that contributed the most to the discrimination of the two main climatic zones (Spearman’s rho = 0.76, *S* = 229.8, *p* < 0.001 and rho = 0.65, *S* = 350.15, *p* < 0.01, respectively for *lfmm2* and *bayenv2*).

The genes putatively involved in local adaptation were tested for gene ontology term enrichment. They were first grouped into four main categories depending on whether they were (i) differentiation outliers or (ii) associated to temperature-related, (iii) precipitation-related or (iv) seasonality-related climate variables. Enrichment was significant for gene ontology terms associated to biological processes related to environmental stimulus detection, metabolic pathways, growth and morphogenesis regulation, as well as biotic interactions (SI Appendix, Fig. S6). Since GO term annotation for the *P. abies* genome is incomplete we also adopted an *ad hoc* approach, specifically focusing on functions of interest, namely, response to photoperiod, cold or abiotic stimuli, growth, flowering and circadian clock. In total we identified 134 candidates SNPs located within or in the vicinity of 81 unique genes involved in these functions. We used a heatmap to illustrate how allele frequencies at these SNPs changed across populations. Populations clustered according to latitude (SI Appendix, Fig. S7) and this clustering was mostly driven by genes associated to the circadian clock and therefore to phenology and growth rhythm: *XAP5 time keeper* (Spearman’s *rho* = 0.70), *flowering-time-like loci* (*FTL, rho = 0*.*59*), *early flowering loci 3* (*EFL3, rho* = 0.92), early *flowering loci 3 high* (*EFL3-high, rho* = 0.91), *sensitivity to red light reduced 1* (*SSR1, rho* = 0.78) and *gigantea* (*rho* = 0.76), (Fig 3A).

**Figure 3:**
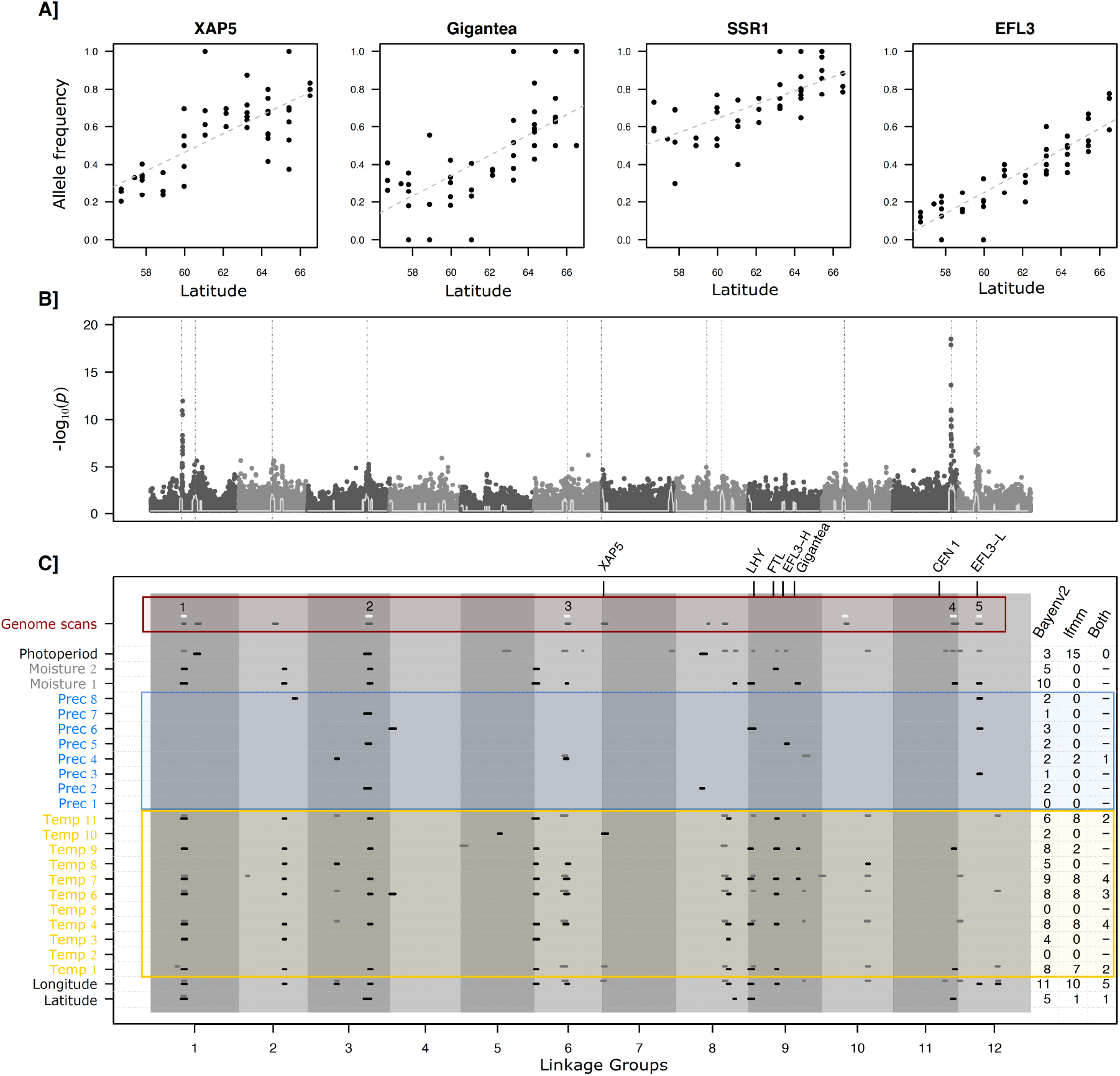
Genome scans for differentiation outliers and genotype-environment association (GEA). **A)** Examples of allele frequency variation as a function of latitude for four candidate genes involved in the control of the circadian clock. **B)** Manhattan plot (-log10 *p*-values) of genome scan for excess of differentiation (*pcAdapt*). Dark and light grey backgrounds delineate linkage groups. Vertical dotted grey lines represent regions enriched for low *p*-values, “peaks” in the profile. A detailed analysis is provided for each of the genome scan and GEA in Supplementary material 3. **C)** For each genome scan (white, *X*^*T*^*X*; grey, *pcAdapt*) and genotype-environment-association (black, *Bayenv2*; dark grey, *lfmm2*) significant peaks are localized on the Norway spruce genetic map. For each geographic and bioclimatic variable, the number of significant peaks is indicated on the right as well as the number of shared peaks. Numbers at the top of the graph identify significant peaks detected by the two genome scans methods and at least one GEA method. When possible, genes involved in the control of circadian clock were placed onto the genetic map.

In spite of the high fragmentation of the Norway spruce reference genome (32), we successfully mapped 89,940 SNPs onto the Norway spruce genetic map (33). Genes putatively involved in local adaptation clustered in a limited number of genomic regions spread across the genome (four genes on average per regions, maximum 14 for *bayenv2* analysis and six on average and maximum 22 for *lfmm2*), with one or several clusters on most linkage groups (Fig. 3B and C, SI Appendix, Section 3). All candidate regions with extreme allele frequency differences between geographic regions were associated to at least two environmental variables, suggesting a direct or indirect causal relationship between high genetic differentiation and environmental factors. Regions enriched for candidate genes were more often associated to temperature-related variables (on average 4.5 ± 3.6 regions across the two GEA analyses, the maximum being nine for *temperature annual range*) than to precipitation-related ones (0.94 ± 1.1, maximum being three for *precipitation of driest quarter*). The climatic variables that contributed the most to the discrimination of the two main climatic zones were also those for which we detected the highest number of genomic regions enriched for candidate genes (Spearman’s *rho* = 0.65; *S = 469*.*32*; *df* = 18; *p* = 0.002 for *bayenv2*). Similar results were obtained with *lfmm2*, the number of candidate genomic regions per variable being highly correlated between the two analyses (*rho* = 0.63, *S = 853, p* < 0.001 and Figure 3C). Genomic regions associated to local adaptation were found across all linkage groups but formed large clusters on individual chromosomes. Taken together, these results and those of the ABC analysis strongly support a significant contribution of natural selection to the establishment and maintenance of the contact zone.

## Discussion

Contact zones are a rich source of information on the interplay between demography and selection in shaping the genetic structure of species (34). Leveraging genomic data from almost 5000 trees sampled across Sweden and the natural range of Norway spruce, we reconstructed the origin of the contact zone separating the south and the north of Scandinavia and showed that natural selection acting on gene clusters dispersed across the whole genome contributed to the differentiation between the two main genetic clusters. Given that Norway spruce has been present in Scandinavia a rather limited number of generations (35), this is an important result with respect to climate change since, unless trees were pre-adapted before invading Scandinavia, it suggests rapid local adaptation.

### A recent contact zone

The general clustering is congruent with what was observed in earlier studies using smaller sample sizes (8) and different markers (14). According to these population genetics studies and the paleo-ecological record (pollen fossil data but also macrofossils) (35-39), current European populations of *P. abies* originate from at least three main ancient refugia located in the Alps, in the Carpathians and in the Russian and Western Siberia Plains. Introgression from Siberian spruce (*P. obovata*) also contributed significantly to the latter, especially at high latitudes (14). What our data show is that these three lineages did not evolve independently but rather entered into contact at many points. For example, as apparent from the Admixture analysis, both Northern Poland and the Russian-Baltic domain, are three ways admixture, with a major contribution from the Carpathians and more limited contributions from the Alps and *P. obovata*.

The recolonization of Northern Europe by *P. abies* started relatively late and spruce migration rates for Fennoscandia varied between 200 and 500 m.year^-1^ (38). Our data supports the existence of two routes of recolonization of Scandinavia, both from east to west, but one entering Scandinavia from the north and moving southward and one entering Scandinavia at a lower latitude and moving both northward and southward (35). The two routes joined between 60°N and 63°N and created an admixture zone that was identified in the present study. Fossil data indicate that trees entered Scandinavia around 13,000-12,000 years ago from the South and 4,000-3,000 years ago from the North (39). The recolonization of Scandinavia by Norway spruce occurred in two phases: a first phase during which small outposts were established and, later on, a second phase when dispersal from those and from a larger front started (39). If their average migration rate was 300 m/year, trees should have reached the current location of the contact zone after around 3300 years and 2000 years, respectively. So, the contact zone would have been created some 2000 years ago, or, assuming a generation time of around 50 years, some 40 generations ago. The pollen fossil record suggests a somewhat lower migration rate and the fronts reaching central Sweden some 3000 years ago, so around 60 generations ago. Of course, these are approximate dates and we do not expect the northwards and southwards migrations to progress at similar speed since it is a well-established fact that Norway spruce can easily be transferred some 3-4 degrees of latitude north without much loss in growth but that a southwards move is generally much less successful (40). We indeed observed an asymmetry, with the southern cluster contributing to the northern one as high as latitudes 66°N while the northern cluster contribution to the southern one was much more limited. In any case, given that gene flow is important in Norway spruce, this implies that one would likely have expected the contact zone to have started to be eroded by gene flow unless it were maintained by selection.

These two recolonization routes are not unique to spruce and are observed in others species, for example, humans, where they are well established (41,42). The resulting admixture zone coincides with a postulated zone of postglacial contact for many plant and animal species (12,43). A similar contact zone is for instance observed in *Populus tremula L*. (44), brown bears (45) or rodents (46). In all these organisms, the contact zone has been initially interpreted as the meeting point between the two main lineages that recolonized Scandinavia after the Last Glacial Maximum (about 25 Kya). In *Populus tremula L*., though, the contact zone corresponds to a sharp change in allele frequency at the FTL gene that is involved in the control of budset (47).

In addition to the main contact zone, in the *conStruct* analysis, populations located north of 62°N contain an ancestral component that was specific to those populations (Fig. 2A, red component). A similar result was obtained by (48) who analyzed the genetic diversity at 15 SSR loci in nine of the breeding populations from northern Sweden. The two northernmost of these nine populations formed a separate cluster in a PCA and both populations presented signs of bottlenecks (48). Those populations are characterized by a higher contribution from *P. obovata*. (11) also observed that trees collected from Northern Fennoscandia and Russia-Urals clustered in a Neighbor-joining tree based on seven SSR loci. Thus, this genetic group reflects how far west *P. obovata* genetic influence was felt, an influence that might have been reinforced locally by bottlenecks during the recolonization process (14). This westward recolonization pattern at high latitudes is not specific to the *P. abies* - *P. obovata* species pair. A similar situation is observed between *Larix sibirica* and *Larix gmelinii* with introgression of mtDNA from the local species in the west, *L. sibirica*, into the invading species from the East, *L. gmelinii* (49-51). This trend does not preclude migration in the opposite direction. For example, *Pinus sylvestris* apparently dispersed primary from western Europe (52).

Finally, pollen analysis and simulations supported a moving front recolonization of Scandinavia rather than population expansion from local refugia (35,38,53). Putative local refugia have been found in mountainous area of central Sweden (54) and might have had a local impact but the fit to an isolation by distance pattern, together with the importance of the contribution of *P. obovata*, would rather argue for a recolonization from populations located outside of the main glaciated areas. Also, these refugial populations are made of small trees that reproduce mainly asexually (54) and it is highly doubtful that they could have contributed massively to surrounding populations. More generally, comparison between *Picea* and *Larix* in Eastern Siberia suggests that *Picea* biology (relatively heavy seeds, low genetic diversity in survival pockets) might explain why *Larix* and not *Picea* was capable of population expansion from small, scattered refugia (55).

### Polygenic architecture of local adaptation along the contact zone

We have so far discussed the data in term of demographic events. However, the major contact zone that was observed in Scandinavia (i) corresponds to a discontinuity in bioclimatic factors, (ii) is better explained by a model incorporating linked selection than by a purely neutral one and (iii) is accompanied by a large number of genomic areas containing clusters of genes characterized by high genetic differentiation and association with climatic variables changing across latitude (e.g. photoperiod, temperature-related climatic variables). While this is not the first study indicating the presence of selection and adaptive cline in forest trees along a latitudinal gradient this is the first one that demonstrates the genome wide impact of local adaptation. The observed pattern is expected under polygenic adaptation for different optima when populations are linked by gene flow (22,23) and could be further reinforced or even caused by structural rearrangements that allow the spread of co-adapted alleles. Unfortunately, the current state of the genome assembly (> 1.5 M scaffolds) does not allow us to investigate further this hypothesis. However, as the largest region includes up to 22 genes carried by different scaffolds, we can expect that some regions enriched for candidate genes are structural variants that can further limit gene-flow between the northern and southern clusters.

A large number of genes were significantly associated to environmental variables and differentiation outliers; 205 unique genes carried at least one significant SNPs associated to environmental variables and 91 were outliers in genome scans. In line with (18) and (21), this suggests a high degree of polygenicity of local adaptation in Norway spruce. Because of the confounded effects of population structure and of the main environmental gradient, these numbers are likely under-estimates (18). The more pronounced pattern of isolation by distance at these loci than at all loci considered jointly strongly suggests that they contribute to local adaptation. This is further supported by the involvement of many of the identified candidate genes in the control of the circadian clock (*XAP5, FTL, EFL-3, EFL-3 high, Gigantea, CEN1, SRR1, LHY*) and therefore in phenology and growth rhythm. Interestingly three important genes for phenology, *FTL, EFL-3 and Gigantea*, are located close by on linkage group 8 (Fig. 3C). This co-localization could have been favored by the strong selection pressure on juvenile trees exerted by frost in late-spring and early-fall (56). In any case, selection on phenology will induce differences in reproductive period that could partly explain the maintenance of the contact zone by limiting the gene-flow between the two clusters.

Considering, the overall low population genetic differentiation together with the relatively short time spent by trees in Scandinavia the establishment of such a strong clinal gradient would seem to imply a rather strong selection pressure, even at individual loci. Assuming that i) local refugia did not contribute significantly to the recolonization of Scandinavia, ii) Norway spruce entered Scandinavia around 10,000-12,000 cal. BP and reached central Sweden around 3,000 cal. BP (11,39) and iii) considering a generation time of about 50 years implies that the observed gradient at adaptive loci over Sweden was established in around 150-200 generations. However, it cannot be ruled out that pre-adapted loci also contributed to local adaptation in newly-colonized areas. As trees from the two main clusters originate from similar latitudes than the ones found today in Scandinavia, a certain level of pre-adaptation seems likely. Additional samples from northwestern Russia and from the Baltics would be necessary to test this hypothesis.

### Practical implications

The genetic structure of the breeding population is important for the management of its genetic resources, for genome-wide association studies and when establishing training sets for genomic selection (57). All our analyses indicate that the individuals used to establish the current breeding population belong to at least seven main genetic clusters. Southern Sweden is particularly complex due to the presence of a large fraction of recent introductions (8) but central Sweden is more homogeneous (CSE) and the northern part of the country consists of two clusters. Our data therefore suggest that at least three training sets may be sufficient to account for most population genetic structure. Importantly, our results also indicate that the current contact zone is maintained by natural selection and will therefore change as the climate does. Three main scenarios for the reaction of Scandinavian population under rapid climate change seem plausible. First, trees from the northern cluster (NFE) are progressively going to be introgressed with genes from the southern cluster as the latter move northwards and the contact zone will progressively disappear. Second, barriers to gene flow are strong enough between the two clusters for the contact zone to persist and shift northwards. Third, assuming that growth traits are a good proxy for fitness, global change will advantage populations with more southern origins, for instance favoring trees with an Alpine or Carpathian genetic background and those will progressively replace existing populations. Given that (18) showed that, at least in the southern and central parts of Sweden, trees with an Alpine or a Carpathian origin outperformed the trees from local provenance for growth traits, this may well occur. This evolution of the contact zone will need to be monitored and incorporated into future genotype-by-climate zone interaction studies for optimizing the delineation of breeding zones, something that, to the best of our knowledge, has not yet been implemented in forest tree breeding.

At any rate, predicting the future evolution of natural populations, for instance for conservation and breeding, is and will remain a complex task, even more so for species such as Norway spruce that are tightly associated to human activity. The detection of adaptive loci that are associated with phenotypic traits and/or environment will not be sufficient to predict future adaptation under climate change scenarios without a deep knowledge of both global and local genetic diversity and how this diversity translates into fitness under various environments. First, introgression from closely related species (or from individuals from outside of the focal range) plays a role in shaping genetic diversity and response to environment. Second, adaptation to a highly dimensional environment requires a high degree of polygenicity. It is therefore intrinsically challenging to extrapolate both genotype-phenotype and genotype-environment relationships under various scenarios involving either demographic or environmental changes. This would require extensive studies, at both local and global geographical scales, repeated over time and with an exhaustive sampling of genetic diversity in the target species but also in species with whom it can hybridize.

## Materials and Methods

### Sample collection

The study was based on 4769 spruce trees, originating mainly from the Swedish Norway spruce (*P. abies*) breeding (Figure 1). Among them, 162 individuals were collected in natural populations across the Norway spruce natural distribution range (8). The remaining trees (4607) were “plus” trees (trees of outstanding phenotype) sampled in Skogforsk (The Forestry Research Institute of Sweden) plantations across Sweden. These trees were genotyped ((8), BioProject PRJNA511374 and (21), BioProject PRJNA731384) using an exome capture target re-sequencing strategy (40,018 diploid 120bp-length probes designed to capture 26,219 *P. abies* genes, (58)).

### Single Nucleotide Polymorphism calling

Raw reads were mapped to the *P. abies* genome reference v1.0 (31) and single nucleotide polymorphisms (SNPs) were identified using HaplotypeCaller v.3.6 (59) and quality filtered. Individuals with more than 50% missing data were also removed (N = 282). The filtered dataset included 4508 individuals and 504,110 SNPs. Those SNPs were annotated based on the most recent genome annotation available for *P. abies* (v1.0, http://congenie.org/).

### Population structure and genotype assignment

For population structure analyses, sites in high linkage disequilibrium (*r*^*2*^ > 0.2) as well as singletons were removed using PLINK v.1.9 (60). Among the remaining SNPs, 155,211 putatively neutral SNPs (i.e. within introns and intergenic regions) were kept for demographic analyses. Population structure was first characterized using a principal component analysis (EIGENSOFT, v.7.2.0 with default parameters, https://github.com/DreichLab/EIG, 61). Trees with unknown geographical origin assigned to a genetic cluster using Random Forest classification as in (8). We also analyzed population structure with ADMIXTURE v1.3 (25) and calculated pairwise fixation indices (Hudson’s estimator of *F*_ST_, 62) between *P. obovata*, admixed *P. abies* x *P. obovata* populations and the *P. abies* genetic clusters defined through the UMAP analysis.

### Spatialized analyses of genetic variation

For the following analyses, only trees that were of confirmed Swedish origin (base on genetic clustering) and with known geographic coordinates were considered (N = 1758). To consider both discrete clusters and continuous distribution of the genetic variation of Norway spruce across Sweden, we first used the *conStruct* software v. 1.03 (3) that combines model-based clustering algorithms with an isolation by distance model. To identify corridors or barriers to gene flow, we used EEMS software (v. 0.0.9000, 26) and we quantified the pattern of isolation by distance by regressing a function of *F*_ST_ (27) over the logarithm of the distance between pairs of populations.

### The contribution of linked selection to the contact zone

In order to test whether linked selection contributed to the establishment and maintenance of the contact zone, we used the program DILS (28). Briefly, DILS implements an Approximate Bayesian Analysis to compare two-population demographic models and identifies the most likely demographic scenario with and without linked selection. Considering that distance to the contact zone might influence demographic inferences (*e*.*g*., hybrids have different history than pure individuals), we created three different datasets as inputs for *DILS* depending on the distance to the contact zone.

### Testing for local adaptation

First, to assess whether the contact zone between the main genetic clusters corresponded to a shift in abiotic conditions across Sweden, we defined climatic zones based on 19 bioclimatic records (Chelsa database v1.2, http://chelsa-climate.org/, 30 arc-second resolution). Different approaches were used to test for the presence of local adaptation at the genomic level and to detect association between genomic polymorphisms and environmental variables. To detect genetic differentiation outliers we used the *Bayenv2* software (29,63) and “*pcadapt*” v4.3.2 R package (30,64). To detect Genotype-environment associations we used “*Bayenv2*” and “*lfmm2”* (31). The same 19 Chelsa bioclimatic variables as those used to define the climatic zone as well as derived combination of those were used for each tree location.

### Candidate genes putative functions and genetic mapping

Gene ontology (GO) enrichment was performed using the ‘topGO’ R package (v2.44.0; (65)). About 60 % of all SNPs were successfully positioned onto the *P. abies* consensus genetic map (32). We developed a new approach to identify regions enriched for outliers (either low *p*-values in *pcAdapt*, and *lfmm2* analyses or high Bayes factor for *Bayenv2*). The method (66) is described in *online methods* and freely accessible at https://github.com/milesilab/peakdetection.

## Acknowledgments

We are grateful to Luis Leal and Linus Söderquist for comments on early versions of the manuscript and to Camille Roux for help with running DILS. This project is supported by the Swedish Foundation for Strategic Research (SSF), grant number RBP14-0040. MT was supported by the EU H2020 project, B4EST. The computation and data handling were provided by the Swedish National Infrastructure for Computing (SNIC) at Uppmax, partially funded by the Swedish Research Council through grant agreement no. 2018-05973.

---

**Table 1:** Pairwise comparison of different models with DILS for individuals at varying distance from the center of the contact zone. Demographic models: Strict Isolation (SI), Ancient Migration (AM), Isolation with Migration (IM), Secondary Contact (SC), Homogeneous and Heterogeneous migration (*Nm*) (M-homo and M-hetero), and Homogeneous and Heterogeneous effective population size (Ne) (Ne-homo and Ne-hetero). The value within parentheses, *p*, is the posterior probability of the best demographic model. Distance to contact zone is defined according to the hybrid index.

| Distance to contact zone | AM vs SI | IM vs SC | M-homo vs M-hetero | N<sub>e</sub>-homo vs N<sub>e</sub>-hetero |
| --- | --- | --- | --- | --- |
| close | AM ($p = 1.00$) | IM ($p = 0.52$) | M-homo ($p = 0.89$) | N-hetero ($p = 0.88$) |
| intermediate | AM ($p = 1.00$) | IM ($p = 0.49$) | M-homo ($p = 0.95$) | N-hetero ($p = 0.93$) |
| far | AM ($p = 1.00$) | IM ($p = 0.55$) | M-homo ($p = 0.89$) | N-hetero ($p = 0.71$) |

## Online methods

### Sample collection

The study was based on 4769 spruce trees, originating mainly from the Swedish Norway spruce (*P. abies*) breeding base population (Figure 1). Among them, 162 individuals were collected in natural populations across the Norway spruce natural distribution range (40 individuals from the hybrid zone between *P. abies* and *P. obovata*, 69 pure *P. abies* and 53 pure *P. obovata*) (8). The remaining trees (4607) were “plus” trees (trees of outstanding phenotype) sampled in Skogforsk (The Forestry Research Institute of Sweden) plantations across Sweden. A total of 873 of those trees lacked geographic information on their origin (Table S1). All trees were genotyped ((8), BioProject PRJNA511374 and (21), BioProject PRJNA731384) using an exome capture target re-sequencing strategy (40,018 diploid probes with 20bp-length were designed to capture 26,219 *P. abies* genes, (58)).

### Single Nucleotide Polymorphism calling

Raw reads were mapped to the *P. abies* genome reference v1.0 (32) using BWA-MEN algorithm with default parameters (67). PCR duplicates were removed using SAMTOOLS v.1.2 (68) and Picard v1.141 (http://broadinstitute.github.io/picard), and INDELs were realigned using GATK v3-5.0 (62). Individual variants were identified using HaplotypeCaller v.3.6 (69) and individual g.vcf were merged using CombineGVCFs v.3.5.0. Variant quality score recalibration (VQSR) was applied following the same procedure as in (70). We additionally filtered SNPs according to four criteria. SNPs were removed if at least one of the following conditions was met: (i) Root Mean Square of the mapping quality (MQ) < 40; (ii) Allele depth (AD) < 2; (iii) genotyping coverage across individuals < 20% and (iv) the site has more than two alleles. Individuals with more than 50% missing data were also removed (N = 282).

The filtered dataset included 4508 individuals and 504,110 SNPs. Those SNPs were annotated based on the most recent genome annotation available for *P. abies* (v1.0, http://congenie.org/): 205,337 (41%) SNPs are within introns, 108,300 (21%) are in intergenic regions and 190,463 are in the exons (38%) of which 63% are synonymous variants (24% of total SNPs) and 37% are nonsynonymous variants (14% of total SNPs).

### Population structure and genotype assignment

For population structure analyses, sites in high linkage disequilibrium (*r*^*2*^ > 0.2) as well as singletons were removed using PLINK v.1.9 (60). Among the remaining SNPs, 155,211 putatively neutral SNPs (i.e. within introns and intergenic regions) were kept for demographic analyses.

#### Principal component analysis

We first conducted a principal component analysis (PCA) using EIGENSOFT with default parameters (v.7.2.0, https://github.com/DreichLab/EIG, (61). We then subset the dataset to insure even number of individuals (N = 80) in the different genetic clusters and re-ran the PCA. The procedure was repeated eight times and a similar clustering was obtained in all runs (SI Appendix, Figure S1).

#### Genotype assignment

The same Random Forest classification procedure as in (8) was then used to infer the geographic origins of the 873 trees whose geographic coordinates were unknown. The assignment was based on genotype similarity on the first five principal components of the PCA (“Random Forest” classification model in R version 3.5.3, ‘*randomForest’* v.4.6-14 package (74,75). The 2572 *P. abies* trees with documented geographical origins and falling in the center of each genetic cluster in the PCA were used as training set. The procedure was repeated 200 times with 8000 iterations to estimate the accuracy of each assignment. If a tree was assigned to the same genetic cluster more than 98% times, it was considered as belonging to that genetic cluster.

#### Model-based clustering

We first estimated population structure using the unsupervised genetic clustering algorithms implemented in ADMIXTURE v1.3 (25) with ten-fold cross validation and 200 bootstraps. The K-value with the lowest cross validation error was retained as the “best” number of theoretical ancestral clusters, but we also reported the results for K varying from 2 to “best-K”+1 as identifying the “true” number of clusters remains an elusive problem (3,76). As for PCA, the whole analysis was repeated on subsets with even number of individuals randomly sampled in each of *P. abies* genetic cluster.

*F*_*ST*_ *estimates*. Pairwise fixation indices (Hudson’s estimator of *F*_ST_, (62)) were then estimated between *P. obovata*, admixed *P. abies* x *P. obovata* populations and the *P. abies* genetic clusters defined through the PCA analysis.

For the following analyses, only trees that were of confirmed Swedish origin (base on genetic clustering) and with known geographic coordinates were considered (N = 1758). We filtered the SNPs dataset to remove loci with > 50% missing genotypes, newly-generated invariant sites and singletons. We retained 113,748 unlinked putatively neutral SNPs (from introns and intergenic regions).

### Spatialized analyses of genetic variation of Norway spruce across Sweden

The Swedish *P. abies* populations are, by and large, continuously distributed and there is a high level of long-distance gene flow (77). To consider both discrete clusters and continuous distribution of the genetic variation of Norway spruce across Sweden, we used the *conStruct* software v. 1.03 (3) that combines model-based clustering algorithms with an isolation by distance model. The 1758 georeferenced samples with genetic Swedish origin were assigned to 47 populations. These “artificial populations” were defined by grouping trees from close geographic origins (N > 5). We ran *conStruct* using K = 1 to K = 7 for one chain and each with 100,000 Markov chain Monte Carlo (MCMC) iterations, and compared spatial and nonspatial models using cross validation across 10 replicates.

### Isolation-by-distance and identification of barriers to gene flow

#### Estimating effective migration surfaces

To identify corridors or barriers to gene flow, we used EEMS software (v. 0.0.9000 (26)). It estimates effective migration surfaces (EEMS) from geographically indexed samples. Sample coordinates and pairwise genetic dissimilarity were used to identify regions with faster or slower change in genetic similarities than predicted under an isolation by distance model. The overall Norway spruce habitat in Sweden was divided into triangular grids corresponding to different deme numbers (30, 50, 80 and 100). Each sample was assigned to the closest point on the grid. To test the stability of the results the program was run five times with 10,000,000 MCMC iterations and 5,000,000 burnins for each deme size. Following authors guideline, we combined estimates over different grids.

#### Isolation by distance estimation through F_ST_

We also quantified the pattern of isolation by distance by regressing a function of *F*_ST_ (27) over the logarithm of the distance between pairs of populations (we used the same populations as those defined for the *conStruct* analysis):

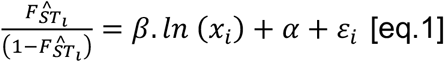

For each pair of sub-populations *i*, 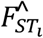 was estimated using vcftools (v0.1.13, (74)), *x*_i_ is the geodesic distance separating a pair of populations (in km, “geosphere” R package v1.5-10, (81)), *β* is the slope of the regression, *α* is the intercept and *ε*_*i*_ is the error term. According to (27), independently of the scale, the slope is inversely proportional to the product of population density *D*, by the second moment of dispersal distance *σ*\*: *β* = 1/(4*πDσ*\*). In Sweden Norway spruce distribution is almost continuous and an even density can be assumed. The “slope” estimate 4*πDσ*\* can thus be interpreted as a neighborhood size, individuals within a neighborhood mating randomly.

### The contribution of linked selection to the contact zone

In order to test whether linked selection contributed to the establishment and maintenance of the contact zone, we used the program DILS (28). Briefly, DILS implements single-population and two-population demographic models and identifies the most likely demographic scenario with and without linked selection. In the case of two-population models that diverged T generations back, four demographic models are possible, namely Strict Isolation (SI), Ancient Migration (AM), Isolation with Migration (IM) and Secondary Contact (SC). The effect of linked selection is estimated through its effect on the distribution of the effective population size, N_e_, or the effective migration rate, Nm, along the genome. *DILS* uses Approximate Bayesian Computations (ABC) and computes summary statistics on the real dataset (nucleotide diversity pi, Tajima’s D, Watterson’s theta, and summary statistics approximating the joint Site Frequency Spectrum), and compare them to a pre-simulated reference table made of 10,000 simulations of 1,000 loci for each demographic scenario. Model comparison is then done with a random forest of 1,000 trees. An error rate per decision tree (e) is estimated, and the posterior probability is computed as 1-e. The population growth was considered constant, mutation rate fixed to 2.763.10^−8^, and priors with a log-Uniform distribution for effective population size (from 100 to 500,000), time of split (from 100 to 1,750,000 generations), and effective migration rate (from 0.4 to 40). Considering that distance to the contact zone might influence demographic inferences (*e*.*g*., hybrids have different history than pure individuals), we created three different datasets of 20 individuals (10 from NFE and 10 from CSE) as inputs for *DILS* depending on the distance to the contact zone. Distance to the contact zone was expressed in terms of hybrid index, the center of the contact zone being arbitrarily defined as the location where the hybrid index is 0.5. The dataset “far” comprised individuals with a hybrid index lower than 0.04 or higher than 0.96; the dataset “intermediate” comprised individuals with a hybrid index between 0.14 and 0.36 or between 0.64 and 0.87; finally, the dataset “close” comprised individuals with a hybrid index between 0.37 and 0.63. Individuals of each dataset were randomly sampled from the respective range of hybrid index. Following software guidelines, a subset of coding regions was used: 1708 coding regions, corresponding to 1% of the exome (from 500 randomly sampled scaffolds).

### Abiotic environment characterization and climatic zones definition

To assess whether the contact zone between the main genetic clusters corresponded to a shift in abiotic conditions across Sweden, we defined climatic zones based on 19 bioclimatic records (Chelsa database v1.2, http://chelsa-climate.org/, 30 arc-second resolution) using an unsupervised clustering approach (i.e. without any *a priori*). Values for each bioclimatic variable were extracted for each geographic coordinate corresponding to land in Sweden and to a grid with nodes every 0.1 degree of latitude and longitude. A principal component analysis was carried out on these data (“*PCA*” function, R package “*FactoMineR*” v1.42, (83)) followed by a hierarchical ascendant clustering approach (“*HCPC*” function, R package “*FactoMineR*”). The optimal number of clusters (*Q*_*opt*_) was defined based on the inertia (*I*) growth between increasing number of clusters (*q*):

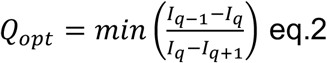

### Testing for local adaptation

Different approaches were used to test for the presence of local adaptation at the genomic level and to detect association between genomic polymorphisms and environmental variables. In line with our analysis of population structure we used two types of approaches. The first family of methods assumes the presence of populations (*n*_*pop*_ = 47) and uses the pattern of allele frequencies within and between these populations to make inference on natural selection or identify genomic areas associated to environmental variation. In contrast, methods from the second group are based on individuals and do not require the *a priori* clustering of individuals into a finite number of populations. The same genomic dataset was used for all subsequent analyses and all SNPs (including also coding sites), with a minimum allele frequency (MAF) higher than 0.05 and less than 50% missing data were kept (*n*_*SNP*_ = 142,766).

#### Detecting excess of differentiation

We used the *Bayenv2* software (41,83) to compute the *X*^*T*^*X* statistic, an analogue to *F*_*ST*_ based on standardized allele frequencies corrected for population structure. To compute the covariance matrix of allele frequencies across the 47 populations, 20 sets of 8,000 non-coding and unlinked loci were randomly selected from the same dataset as the one used in the *conStruct* analysis. The covariance matrix was obtained by averaging over the last matrices generated by the 20 independent runs of 100,000 MCMC iterations. Only SNPs with a *X*^*T*^*X* higher than 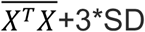 where SD is the standard deviation of the *X*^*T*^*X* distribution were considered significant (> 0,993 quantile).

We also used a PCA-based outlier detection method implemented in the “*pcAdapt*” v4.3.2 R package (30,84). The method assumes that the main part of the SNPs variation along principal component axes reflects demographic processes and population structure. Extreme values correspond to outliers SNPs that are presumed to be in the vicinity of SNPs involved in adaptation. To ensure that the observed pattern is not driven by a small region with extended linkage disequilibrium, PCA loadings were checked. To control for false positives, only SNPs with FDR *q*-value < 0.1 were considered as putatively involved in local adaptation.

#### Genotype-environment associations

In order to further characterize the genetic basis of local adaption in Norway spruce, we conducted genotype-environment associations (GEAs). As for the detection of excess of differentiation, we used two different approaches, one population-based, “*Bayenv2*” and one individual-based, “*lfmm2”* (30). The same 19 Chelsa bioclimatic variables as those used to define the climatic zone were used for each tree location. Three additional climatic variables were computed: (i) annual heat-moisture index (AHM, annual mean temperature / (total annual precipitations/100)), (ii) summer heat-moisture index (SHM, average temperatures of warmest month / (total precipitations of warmest quarter/100)) and (iii) average day length difference between June and January; the latter was used as a proxy for the growth period (SI Appendix Table S2).

**Bayenv2** tests for a correlation between allele frequencies and an environmental variable by using a Bayesian generalized linear mixed model. A variance-covariance matrix of allele frequencies is incorporated as random effect to correct for population structure. For each climatic variable plus latitude and longitude, both Bayes Factor (BF) and Spearman’s *rho* correlation coefficient were computed to measure the intensity of the association between allele frequency variation and environmental variation. For each climatic variable, the following filtering (based on Bayes factor and Spearman’s rho) was applied to retain only the most relevant SNPs: (a) the SNPs were ranked according to their Bayes factor (BF) and a SNP was retained if i) its BF > 100 (very strong strength of evidence according to (85) or, if ii) its BF > 20 (strong strength of evidence) and it was within the 0.1% highest BF.

**Latent factor mixed models** were also used to test for associations between the set of environmental / geographic variables and SNPs variation. The “*lfmm2*” function (“LEA” R package) was used to estimate latent factors based on an exact least-squares approach. Missing genotypes were imputed using the “impute” function following author recommendations. The number of latent factors to be included in the analysis was determined using the *“snmf”* function (MCMC, 10 repetitions, 6% of dataset masked per repetitions). It was defined as being the one minimizing the cross-entropy criterion across all runs. The latent factors were used to correct for population structure in linear regressions between genotypes and environmental variables; *p*-values were recalibrated by using genomic control after correction for confounding effect from population structure (“*lfmm2*.*test*” function). To control for false positives, only SNPs with a *q*-value < 0.1 were considered (“*qvalue*” v2.20.0 R package, method “*fdr*”, Storey et al. 2020).

#### Candidate genes putative functions

Gene ontology (GO) enrichment was performed using the ‘topGO’ R package (v2.44.0; (86). Annotation from ConGenIE (the Conifer Genome Integrative Explorer, http://congenie.org/) was used as reference (i.e. custom input). For various lists of candidate genes, defined through both genome-scans or GEA, enrichment of genes in particular GO terms for biological processes (BP) was assessed using Kolmogorov-Smirnov’s tests “*elimKS”. rrvgo* (87) was then used to summarize gene ontology terms by collapsing redundant terms across hierarchical levels using *Arabidopsis thaliana* as a reference for GO term (v.3.13.0, (88)).

#### Genetic map positioning of loci putatively involved in local adaptation

About 60 % of all SNPs were successfully positioned onto the *P. abies* consensus genetic map (33). We developed a new approach to identify regions enriched for outliers (either low *p*-values in *pcaAdap*, and *lfmm2* analyses or high Bayes factor for *Bayenv2*). The method (66) is described and freely accessible at https://github.com/mtiret/gwas-snp-detection. For each genome scan or GEA, we searched for regions with more outlier SNPs (top 5% of the distribution) than expected under a random distribution. In a first step, we generated a null expectation through randomization (10,000 runs) to determine the maximum number of outliers expected by chance within each 300 bp window. This number was then used as a threshold for peak detection in the actual data. Finally, to be conservative, only peaks containing at least one SNP detected as significant after correction for multiple testing were considered as candidate regions.

