## Supplementary File for "Teasing apart the joint effect of demography and natural selection in the birth of a contact zone"

### Supporting Information

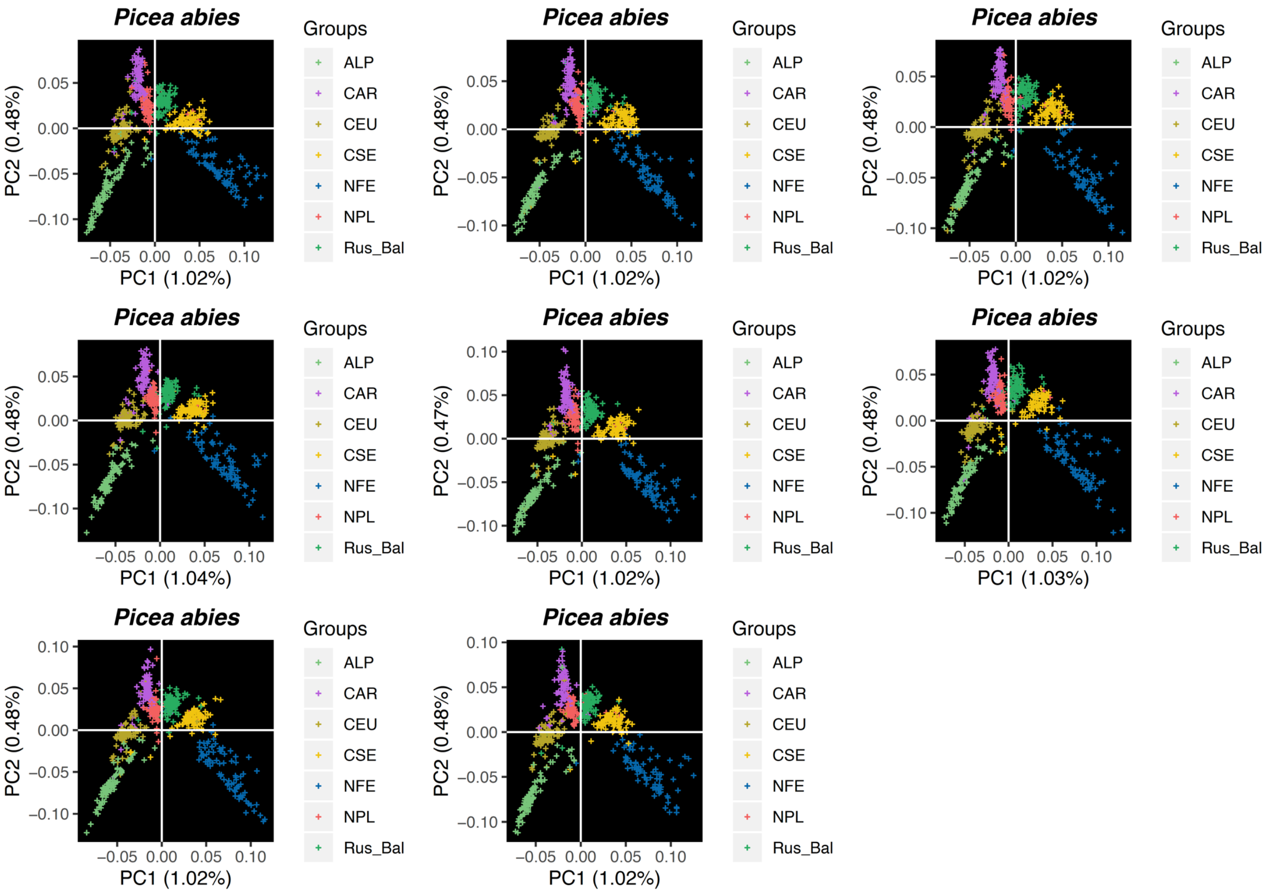


**Figure S1: PCA based on resampling of 80 samples per cluster.**

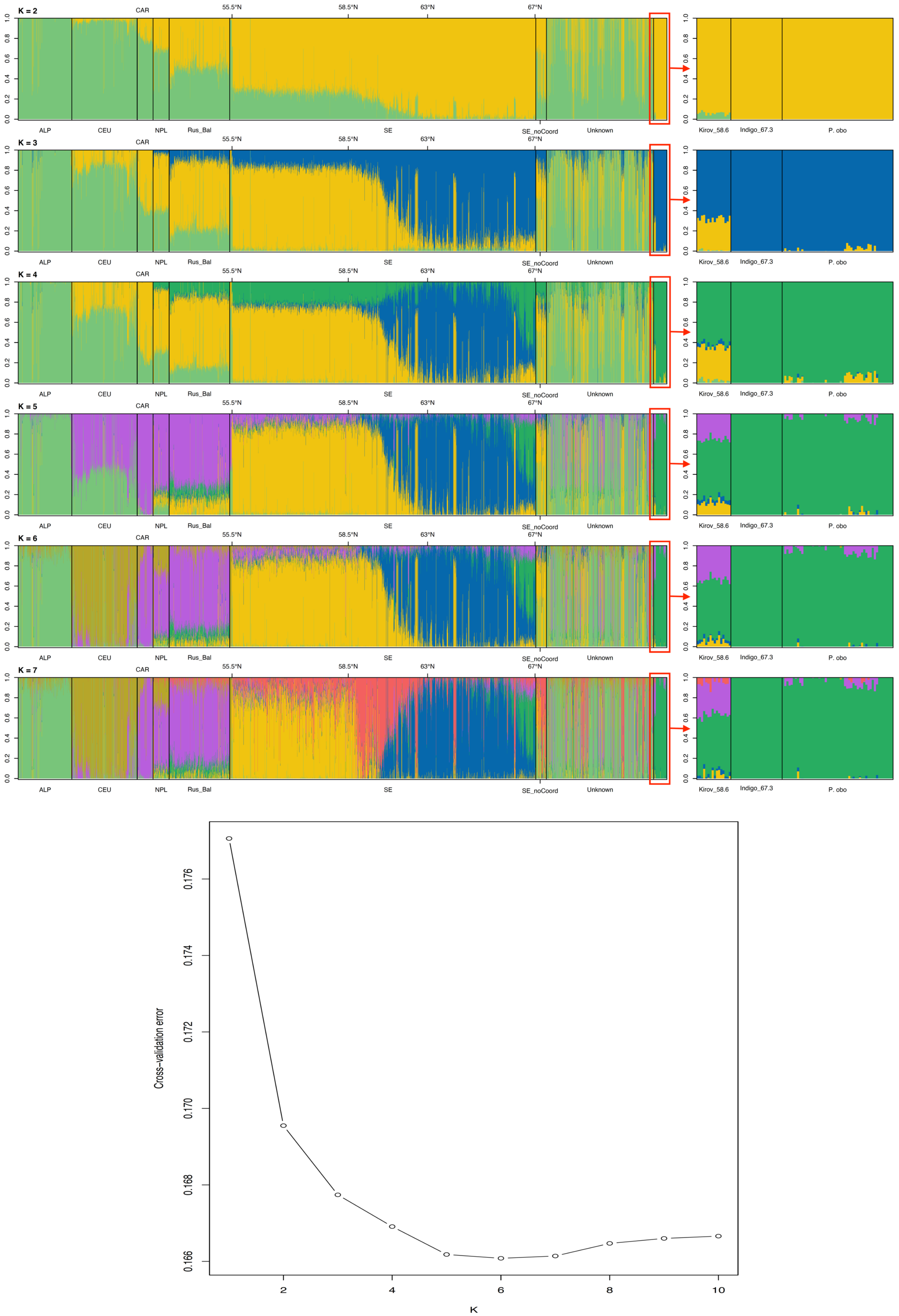


**Figure S2: Admixture plot (K 2 to 7) and cross validation error plot.**

**Table S1** Pairwise $F_{ST}$ index estimates among *P. obovata*, two admixed *P. abies* x *P. obovata* populations (Indigo and Kirov), and the seven main *P. abies* genetic clusters.

| | Kirov | Indigo | NFE | ALP | CAR | CSE | Rus_Bal | NPL | CEU |
|---|---|---|---|---|---|---|---|---|---|
| *P. obo* | 0.064 | 0.042 | 0.079 | 0.168 | 0.144 | 0.095 | 0.112 | 0.136 | 0.16 |
| Kirov | | 0.116 | 0.054 | 0.12 | 0.098 | 0.06 | 0.07 | 0.09 | 0.111 |
| Indigo | | | 0.14 | 0.241 | 0.217 | 0.162 | 0.183 | 0.209 | 0.233 |
| NFE | | | | 0.067 | 0.047 | 0.018 | 0.028 | 0.041 | 0.058 |
| ALP | | | | | 0.023 | 0.039 | 0.027 | 0.018 | 0.01 |
| CAR | | | | | | 0.022 | 0.01 | 0.006 | 0.012 |
| CSE | | | | | | | 0.009 | 0.016 | 0.029 |
| Rus_Bal | | | | | | | | 0.006 | 0.017 |
| NPL | | | | | | | | | 0.008 |

Abbreviations; *P. obo*: *Picea obovata*; *Picea abies*: NFE: Fennoscandian; ALP: Alpine; CAR: Carpathian; CSE: Central and Southern Sweden; Rus_Bal: Russian-Baltic; NPL: Northern Poland; CEU: Central Europe.

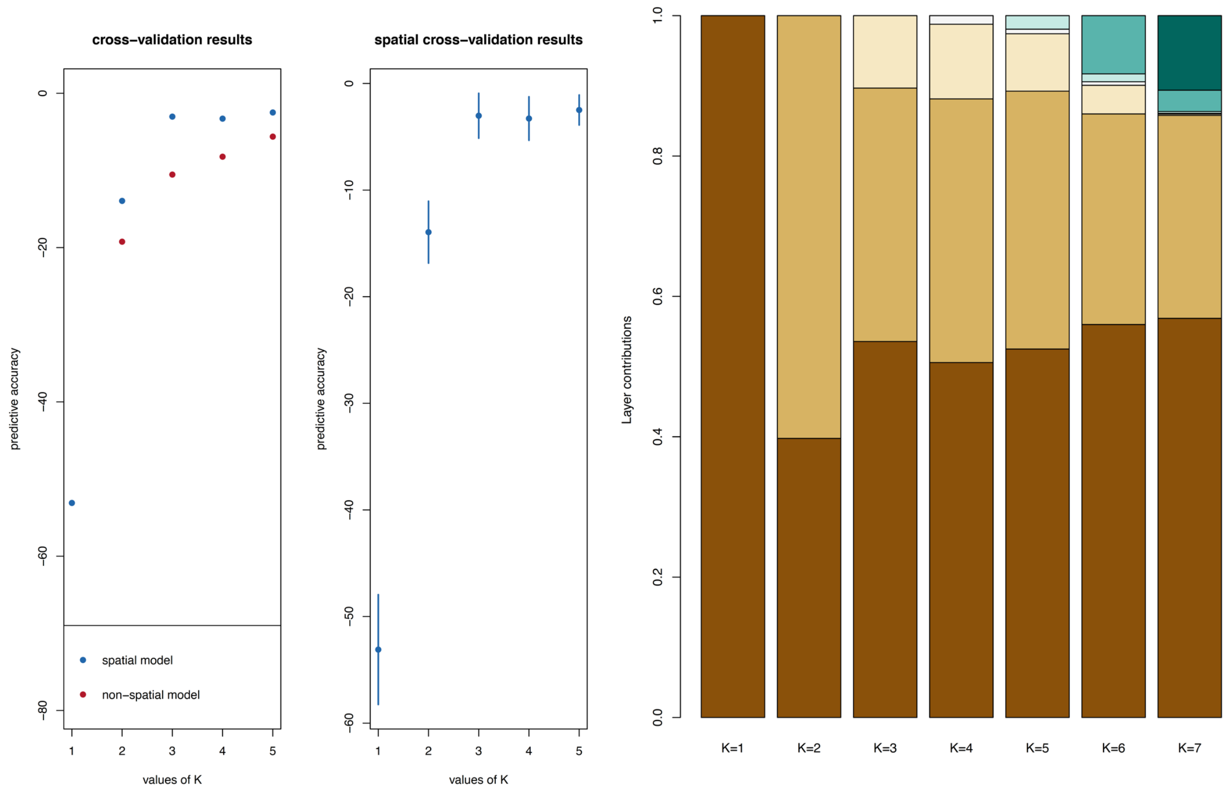


**Figure S3: construct cross-validation and layer contribution plots.**

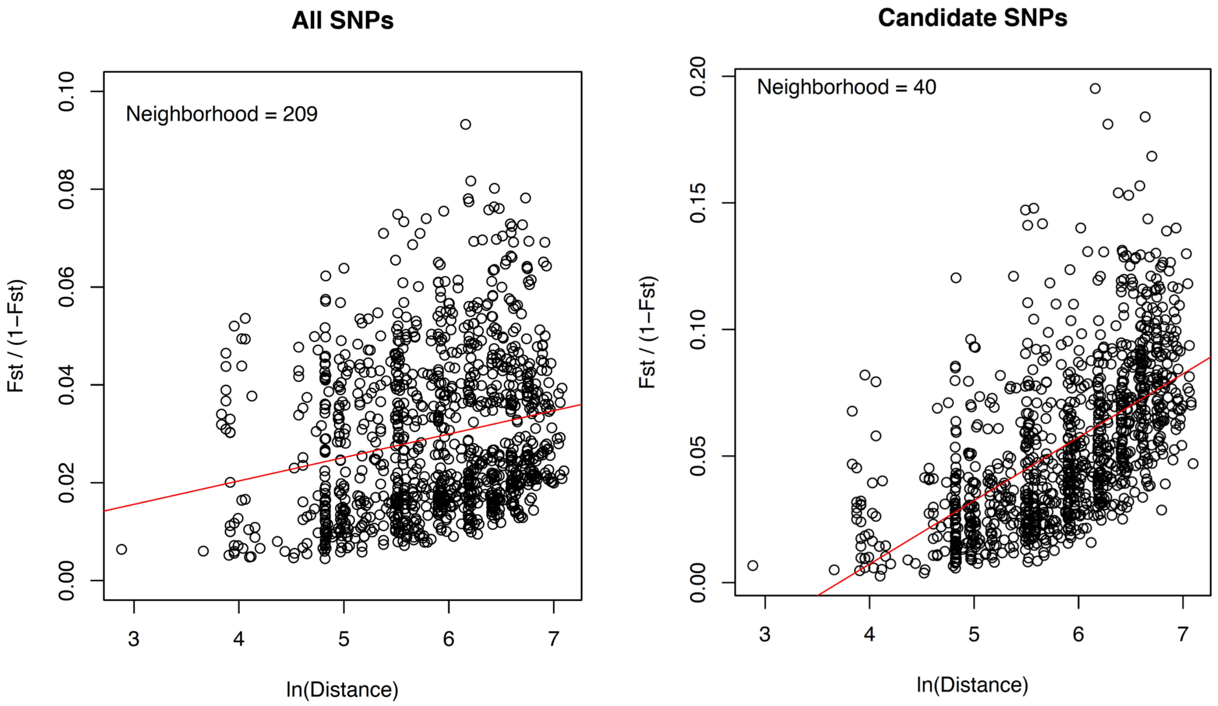


**Figure S4: isolation by distance pattern for all SNPs (left panel) or candidate SNPs (right panel).** Neighborhood size in indicated in the top of the graph, the lower the stronger the IBD.

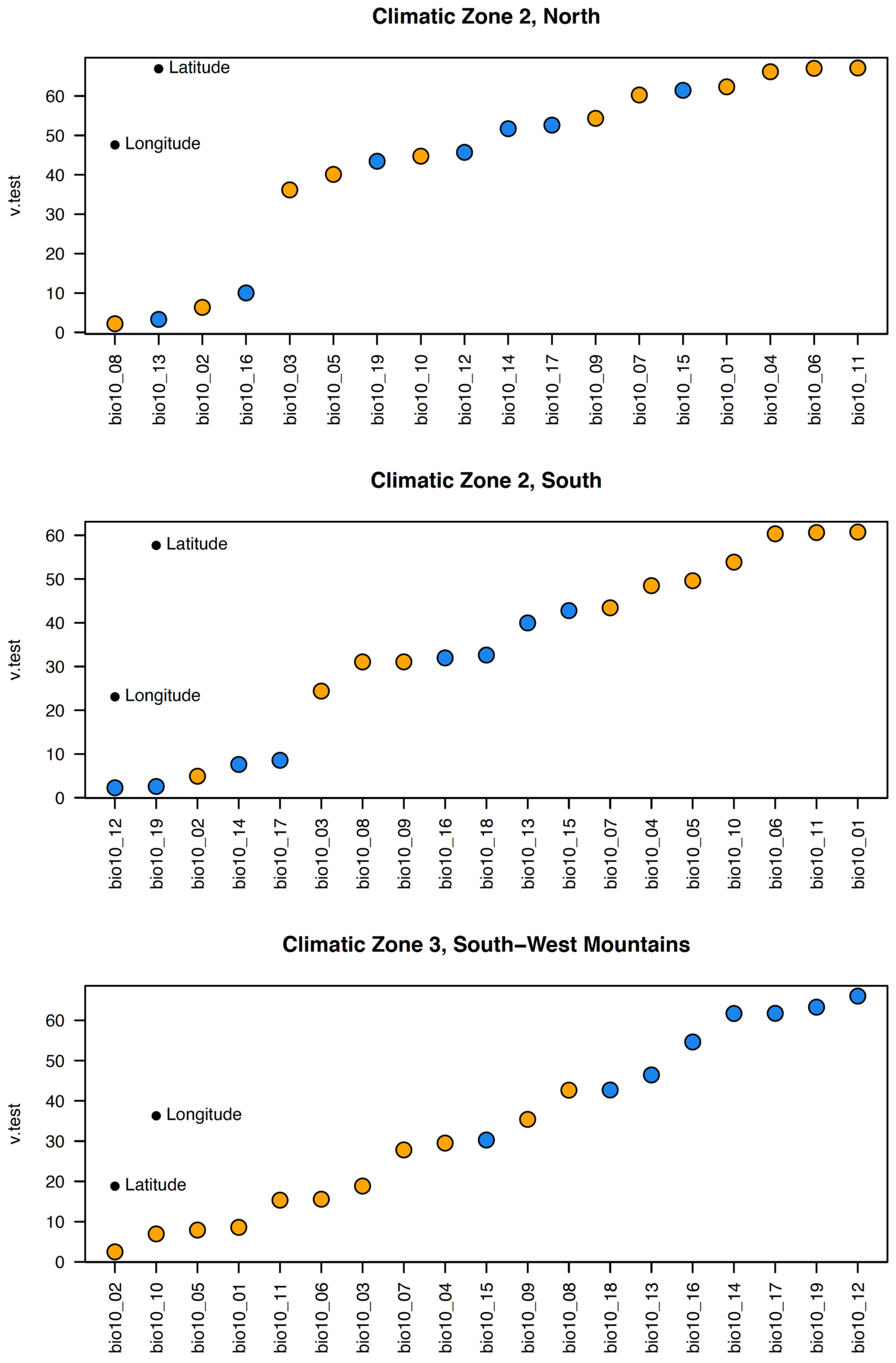


**Figure S5: Relative contribution of the climatic variable to each of the three climate zones.** The contribution of each bioclimatic variables (v.test value, the higher the stronger the contribution) is given for each of the climatic zone. Precipitation related variables are in blue and temperature related variables are in orange. For information, the contribution that latitude and longitude would have had if they were used together with the bioclimatic variables to define the climatic zone is also given.

**Table S2:** Number of candidate loci identified through genotype-environment associations.

| Climate variables | Bayenv2 | | lfmm2 | |
|---|---|---|---|---|
| | SNPs | Genes | SNPs | Genes |
| Longitude | 143 | 124 | 8 | 7 |
| Latitude | 263 | 196 | 795 | 627 |
| Annual Mean Temperature (Temp. 1 / Bio10_01) | 174 | 150 | 568 | 471 |
| Mean Diurnal Range (Temp. 2/ Bio10_02) | 63 | 57 | 1 | 1 |
| Isothermality (Temp. 3 / Bio10_03) | 143 | 112 | 23 | 22 |
| Temperature Seasonality Temp. 4/ Bio10_04) | 143 | 122 | 818 | 632 |
| Max Temperature of Warmest Month (Temp. 5 / Bio10_05) | 86 | 83 | 0 | 0 |
| Min Temperature of Coldest Month (Temp.6 / Bio10_06) | 143 | 116 | 758 | 605 |
| Temperature Annual Range (Temp. 7 / Bio10_07) | 143 | 122 | 680 | 552 |
| Mean Temperature of Wettest Quarter (Temp. 8 / Bio10_08) | 262 | 220 | 13 | 13 |
| Mean Temperature of Driest Quarter (Temp.9 / Bio10_09) | 143 | 122 | 94 | 90 |
| Mean Temperature of Warmest Quarter (Temp.10 / Bio10_10) | 143 | 131 | 7 | 5 |
| Mean Temperature of Coldest Quarter Temp. 11 / Bio10_11) | 143 | 121 | 788 | 630 |
| Annual Precipitation (Prec.1 / Bio10_12) | 72 | 68 | 0 | 0 |
| Precipitation of Wettest Month (PrecWet_Month) (Prec. 2 / Bio10_13) | 143 | 133 | 0 | 0 |
| Precipitation of Driest Month (Prec. 3 / Bio10_14) | 40 | 40 | 3 | 3 |
| Precipitation Seasonality (Prec. 4 / Bio10_15) | 140 | 124 | 228 | 205 |
| Precipitation of Wettest Quarter (Prec. 5 / Bio10_16) | 118 | 104 | 0 | 0 |
| Precipitation of Driest Quarter (Prec. 6 / Bio10_17) | 64 | 60 | 2 | 2 |
| Precipitation of Warmest Quarter (Prec. 7 / Bio10_18) | 143 | 126 | 0 | 0 |
| Precipitation of Coldest Quarter (Prec. 8 / Bio10_19) | 53 | 52 | 10 | 10 |
| Average day length in June – in January (Photperiod) | 257 | 205 | 0 | 0 |
| Annual Heat_Moisture Index (Moist. 1) | 143 | 119 | 0 | 0 |
| Summer Heat_moisture Index (Moist. 2) | 136 | 117 | 781 | 589 |

**Note:**

Bayenv2: the number of SNPs with BF >150 or 20 in 0.1% upper tail.

Lfmm2: Number of SNPs for with *q*-value < 0.1.

Transcript: the number of genes in which candidate loci falls in.

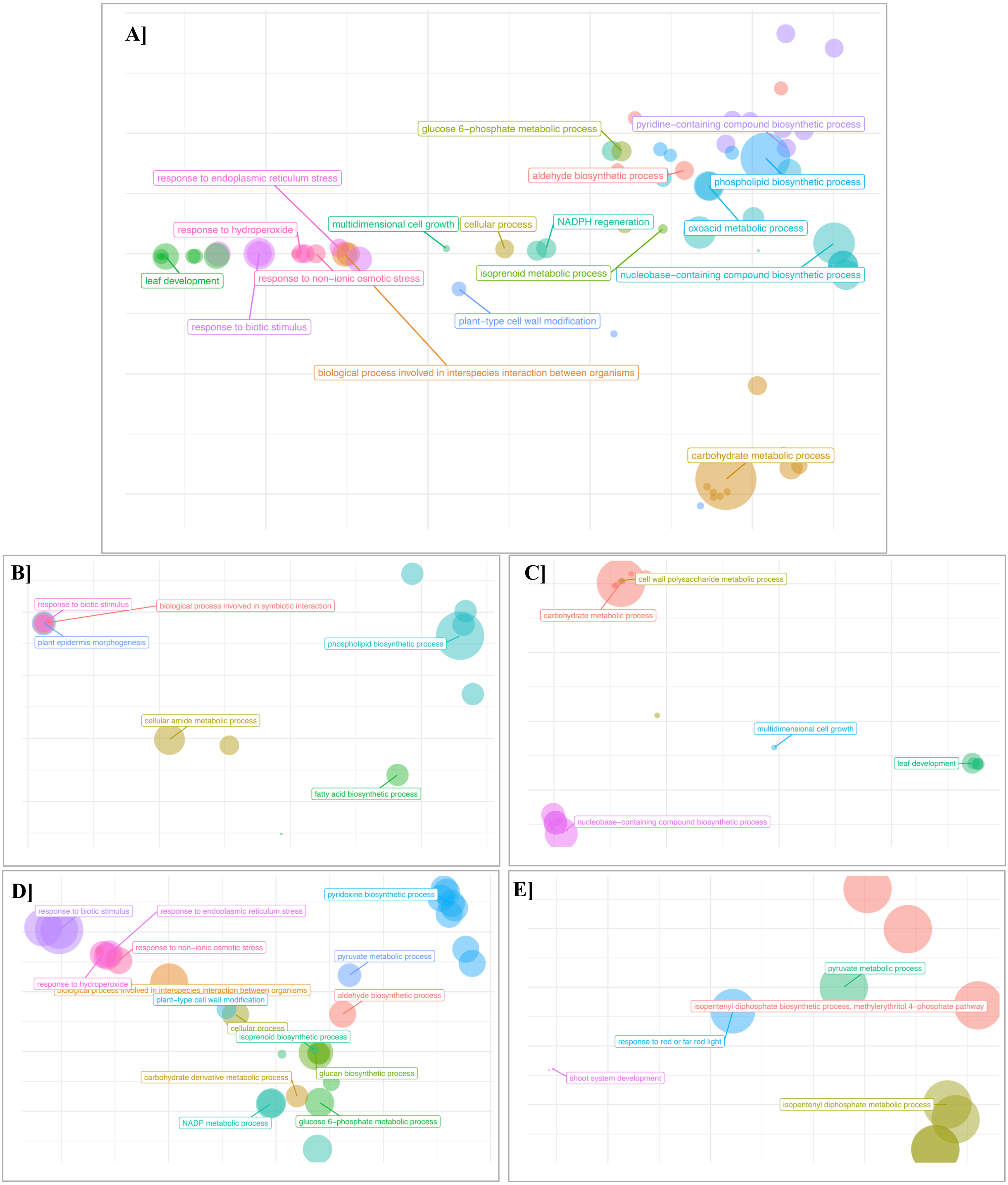


**Figure S6: Scatter plot based on GO term enrichment analysis. A** is for all four categories together, genome scans (**B**), precipitation-related (**C**), temperature-related (**D**) or seasonality related (**E**) candidate SNPs. Colors indicate different "parent-term", dot size is proportional to the number of GO term collapsed under a given parent-term.

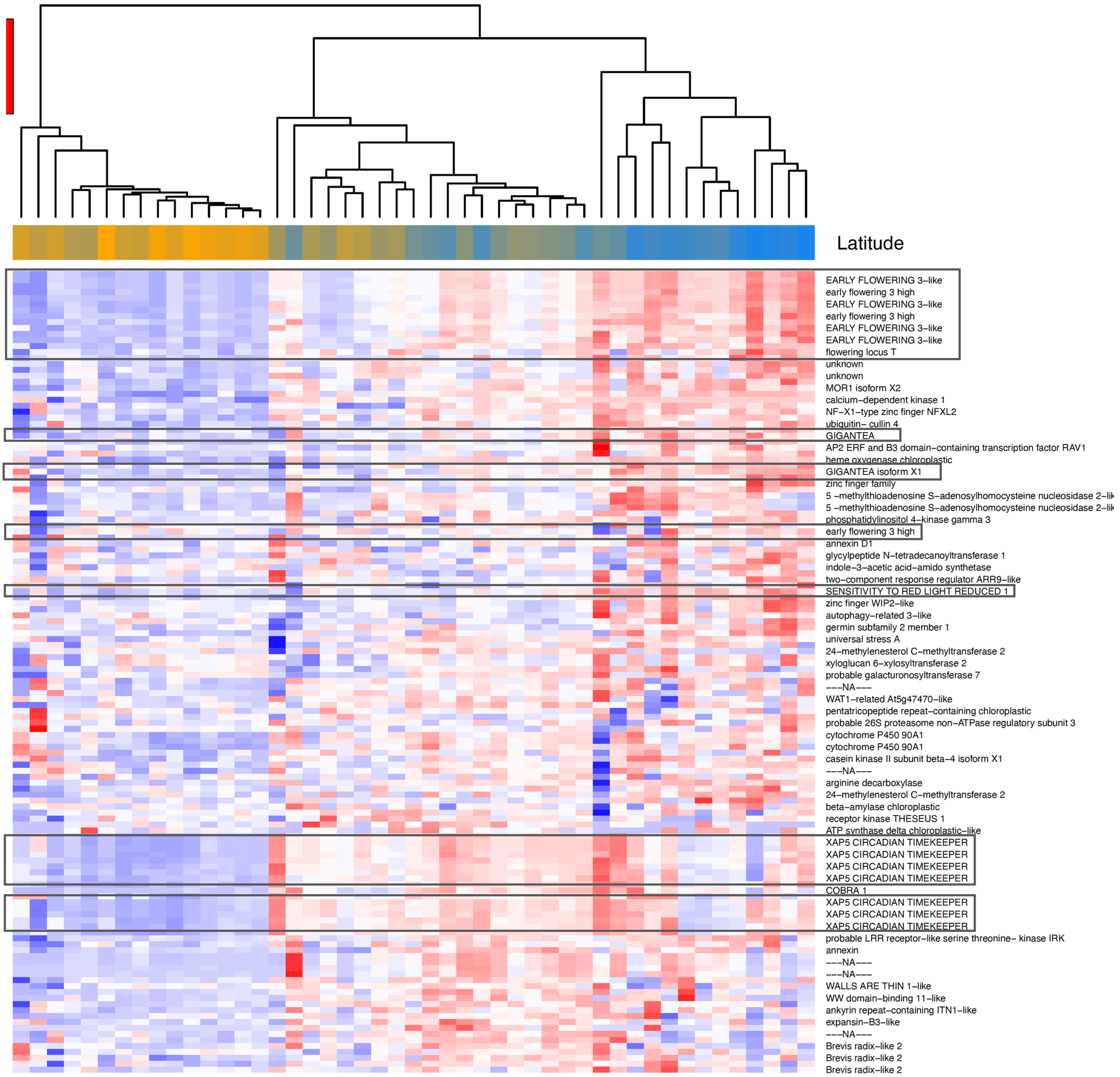


**Figure S7: Heatmap of allele frequency variation between population for candidate SNPs detected with the *ad hoc* procedure.** The dendrogram show populations clustering. For each population the color indicates the latitude from the southernmost (orange) to the northernmost (blue) population. Candidate SNPs located in genes involved in the control of the circadian clock are boxed.

### Section 2: Simulating a neutral demography of NFE vs CSE with forward simulations

In order to test whether the divergence and contact zone between the NFE and CSE clusters could be explained by a purely demographic model we simulated two diverging populations with a neutral demographic history. If it is possible to simulate a population close to the real one in terms of Hudson's $F_{ST}$ and ancestry coefficient, then we cannot reject the hypothesis that the current contact zone was entirely shaped by neutral demographic processes. If no demographic parameters allow us to retrieve the observed pattern of polymorphism and divergence between the two clusters then we will be able to reject the hypothesis that the current contact zone has been solely shaped by demography. We used a grid search approach to identify the set of demographic parameters that best approximate the real Hudson's $F_{ST}$ (0.018) *and* the shape of the cline as described by ordered ancestry coefficients computed by ADMIXTURE. More specifically, we simulated a Wright-Fisher model, forward in time, with two sub-populations, NFE and CSE, with N = 15,000 diploid individuals each. First, the allele frequencies in the ancestral population were drawn from the real Site Frequency Spectrum fitted to a beta distribution of parameters alpha = 0.5853521 and beta = 14.3326236. Second, alleles were sampled without replacement to create the two sub-populations. Third, the two populations were let to diverge during $t_{div}$ generations, with a low migration rate between them, $m_{div}$. Fourth, to mimic a secondary contact, the two populations were let to diverge during $t_{sc}$ generations, with a higher migration rate $m_{sc}$. At the end, to mimic the real sample, 1,734 individuals with 113,748 biallelic SNPs were sampled, half of them from NFE and the other half from CSE. The parameter grid was composed of the divergence time, $t_{div}$, and the migration rate during the secondary contact, $m_{sc}$, ranging from 0.1 to 20 times N (15,000) and from 1e-4 to 5e-2, respectively. $m_{div}$ was fixed to 8.18e-6 as estimated by Fastsimcoal2 previously; likewise, $t_{sc}$ was fixed to 140 generations, as a rough approximation of the time since secondary contact: Norway spruce first reached Sweden 7000 years ago (Giesecke and Bennett, 2004), and we assumed a generation time of around 50 years, so that the estimated duration of the secondary contact was 7000/50 = 140 generations.

For each simulation, we computed Hudson's estimator of $F_{ST}$ and the ancestry coefficients with ADMIXTURE (with K fixed to 2). In order to quantify the distance between the simulations and the real dataset, we computed the distance in $F_{ST}$ (eq.1),

$$d_{F_{ST}} = \left(F_{ST}^{observed} - F_{ST}^{simulated}\right)^2 \quad \text{Eq.1}$$

and the sum of squares of the ancestry coefficients (eq.2),

$$d_{Anc} = \sum_{i=1}^{1734} \left(Anc_{i,observed} - Anc_{i,simulated}\right)^2 \quad \text{Eq.2}$$

where $Anc_i$ is the *ith* ordered ancestry coefficient.

No set of demographic parameters, i.e. total divergence, $t_{div}$, and migration rate during the secondary contact, $m_{sc}$, allowed us to retrieve simultaneously the observed divergence between the two clusters *and* the ancestry coefficients over the two clusters (Fig S1 and S2): $d_{FST}$ is approximately a constant function over $t_{div}$, and a convex function over $m_{sc}$ with a minimizer of $m_{sc}$ = 0.007; on the other hand, $d_{Anc}$ is approximately a convex function over $t_{div}$ and $m_{sc}$ with a minimizer of $t_{div}$ = 5N, and $m_{sc}$ = 0.035. Since the minimizers do not overlap, the hypothesis that the contact zone was established solely through demography seems highly unlikely.

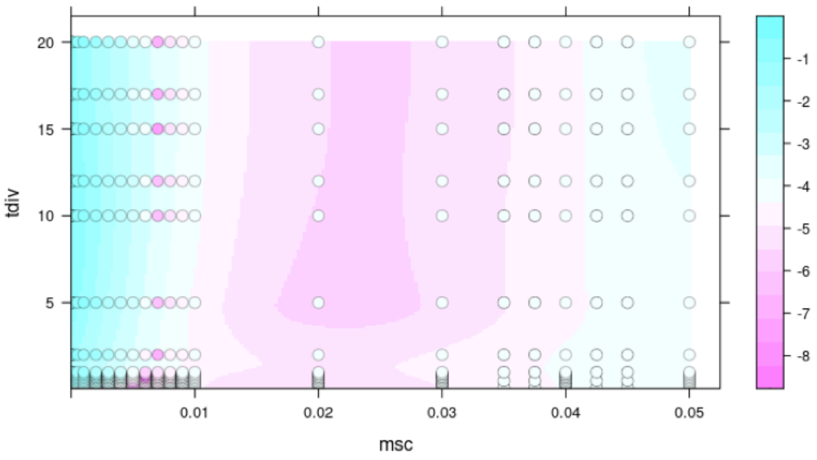


**Figure S1:** Heatmap of $d_{FST}$ (distance between the simulated and the real $F_{ST}$), obtained with a grid search approach over the parameter divergence time $t_{div}$ (expressed in unit of N = 15,000) and the migration rate $m_{sc}$. The minimum values of $d_{FST}$ were obtained along the vertical line $m_{sc}$ = 0.007.

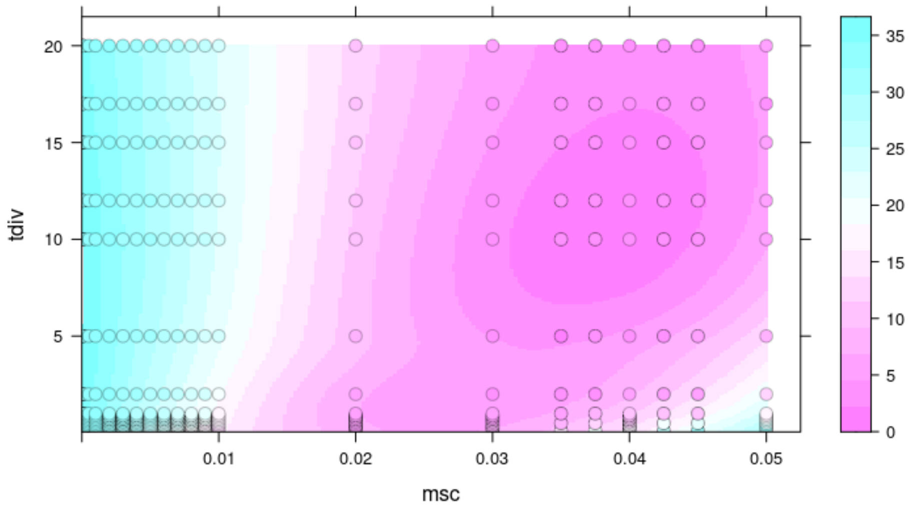


**Figure S2:** Heatmap of $d_{Anc}$ (distance between the simulated and the real ancestry coefficients), obtained with a grid search approach over the parameter divergence time $t_{div}$ (expressed in unit of N = 15,000) and the migration rate $m_{sc}$. The minimum values of $d_{Anc}$ were obtained for $t_{div}$ = 5N, and $m_{sc}$ = 0.035.

### Simulating a neutral demography of NFE vs CSE with fastsimcoal

In order to complete the grid analyses with a likelihood-based approach, we used *fastsimcoal2* (Excoffier et al., 2013) and *abcrf* R package (Marin et al., 2019) to implement an Approximate Bayesian Computation (ABC) pipeline. We simulated two populations (CSE and NFE) of 10,000 loci with gene flow and a bottleneck (in each population, occurring at the same time and of the same strength). We draw 1,000 sets of priors (Table S1), then simulated 10 replicates of each (ending up with 10,000 simulations). Ten diploid individuals were sampled from each population, and we computed classic population genetic summary statistics (*Pi,* Tajima's *D*, $F_{ST}$, and variance of $F_{ST}$ across the genome) with Arlequin (Schneider et al., 2000). To avoid problem from classic rejection algorithm, these summary statistics were used to constitute a reference table and analysed with random forest (Pudlo et al., 2016). Prior values are given in table S1.

Simulating a backward process with selection is difficult, and as such, *fastsimcoal2* is not able to simulate a selection scenario *per se.* However, we can categorize simulations with a high variance of $F_{ST}$ across the genome to be simulating a linked-selection (Fraisse et al., 2020), and assess whether our real data is closer to a "linked-selection" scenario or not. The variance of $F_{ST}$ was then used as a classifier: if the variance is higher than the median value, the simulation is categorized as "selection"; otherwise as "neutral". We then used the reference table (*Pi*, Tajima's *D* and $F_{ST}$) and this categorization to grow a random forest with *abcrf* (1,000 trees, with an out-of-bag prior error rate of 30.0%). Summary statistics were also computed with Arlequin on the real data (thinned to 10,000 SNPs), and was categorized as a "selection" scenario with this random forest, with a posterior probability of 97.1% (1,000 trees).

**Table S1**: Priors of each parameters used for the simulations with fastsimcoal.

| Variable | Distribution | Min | Max |
|---|---|---|---|
| NCSE | log-uniform | 1e3 | 1e5 |
| NNFE | log-uniform | 1e3 | 1e5 |
| TB | log-uniform | 1e2 | 1e3 |
| TC | log-uniform | 1e4 | 6e5 |
| MIG | log-uniform | 1e-8 | 1e-3 |
| BOT | log-uniform | 1e1 | 1e2 |
| GR | log-uniform | 1e-2 | 1 |

NCSE: CSE population size; NNFE: NFE population size; TB: time until bottleneck (for both CSE and NFE); TC: time until coalescence (between CSE and NFE), MIG: symmetrical migration rate until TC; BOT: strength of bottleneck; GR: population size reduction at TC.

### Section 3

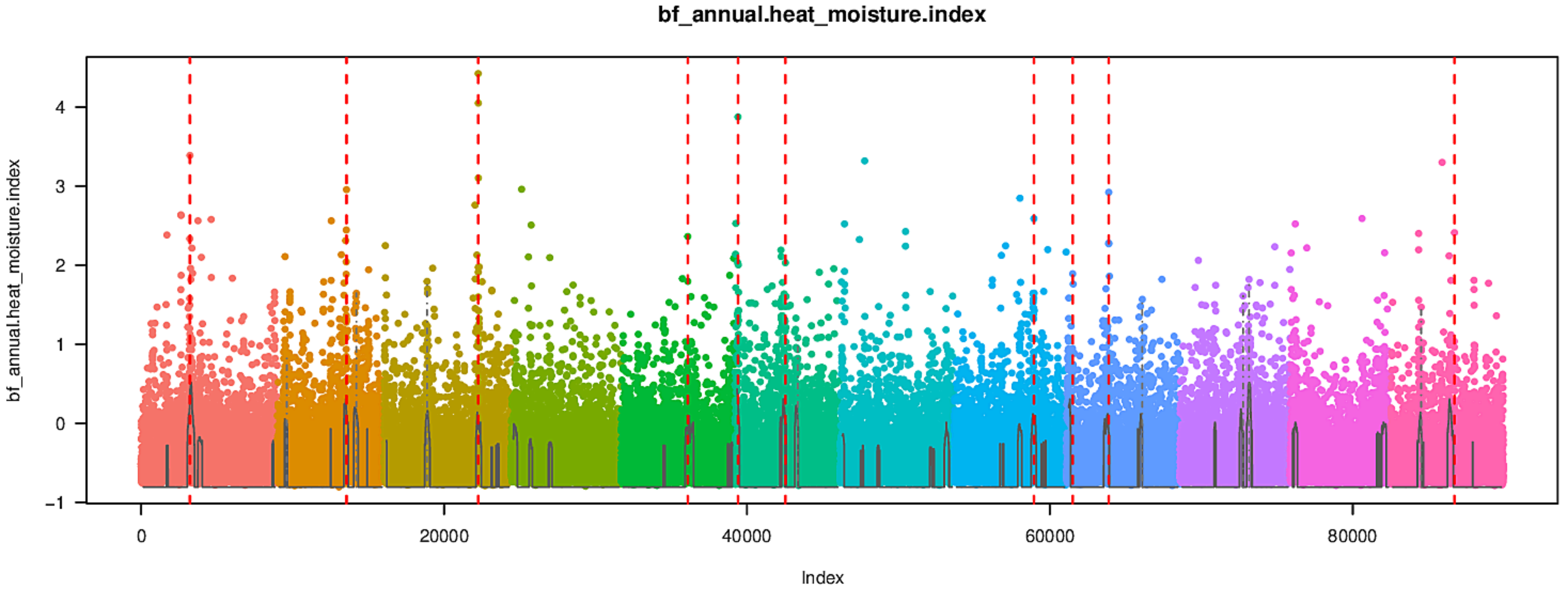
bf_annual.heat_moisture.index
bf_annual.heat_moisture.index
Index
4
3
2
1
0
−1
0
20000
40000
60000
80000

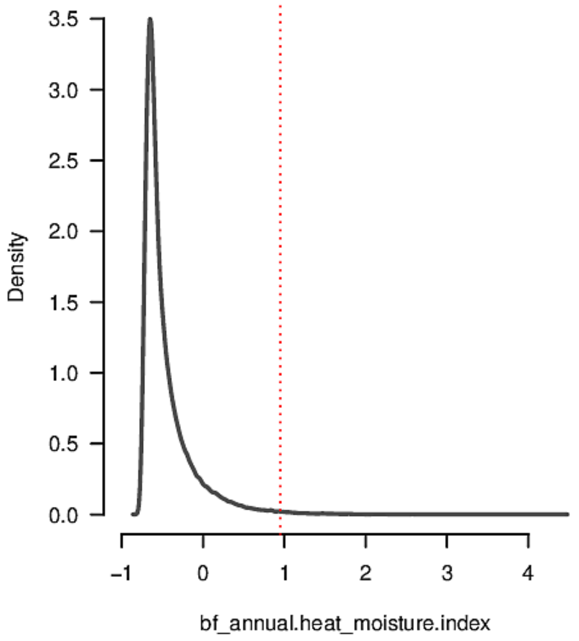
Density
bf_annual.heat_moisture.index
3.5
3.0
2.5
2.0
1.5
1.0
0.5
0.0
−1
0
1
2
3
4

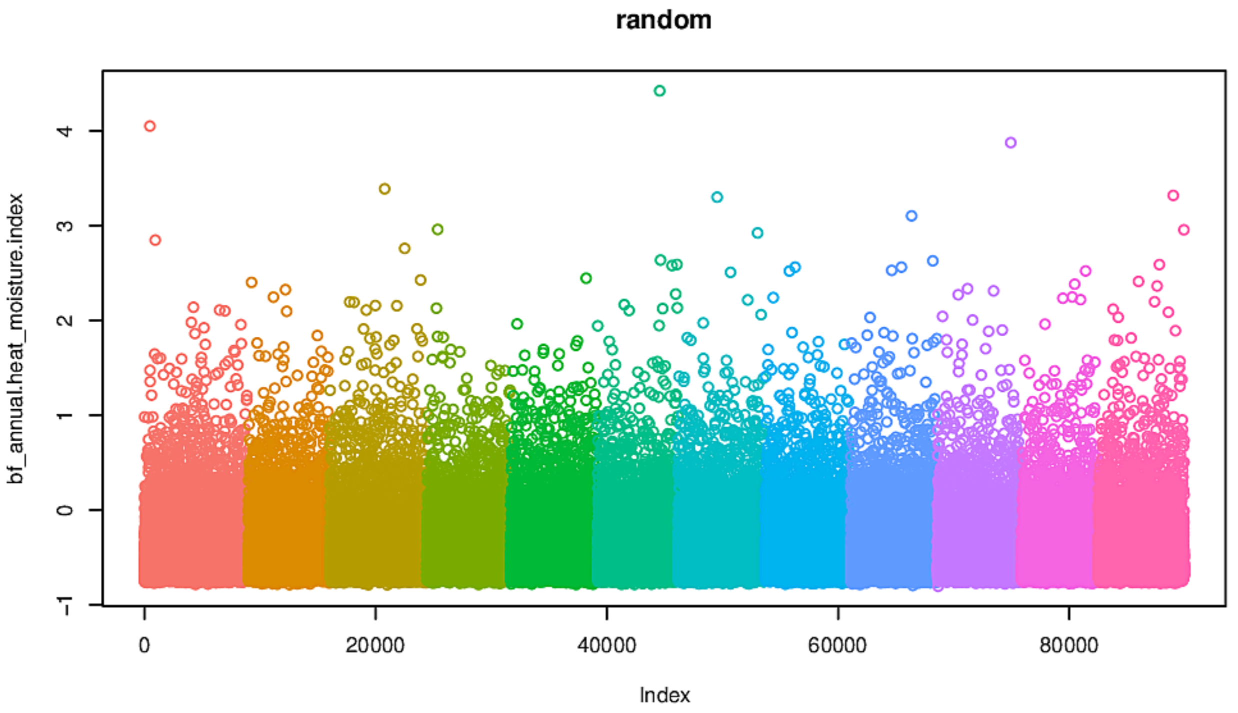
random
bf_annual.heat_moisture.index
Index
4
3
2
1
0
−1
0
20000
40000
60000
80000

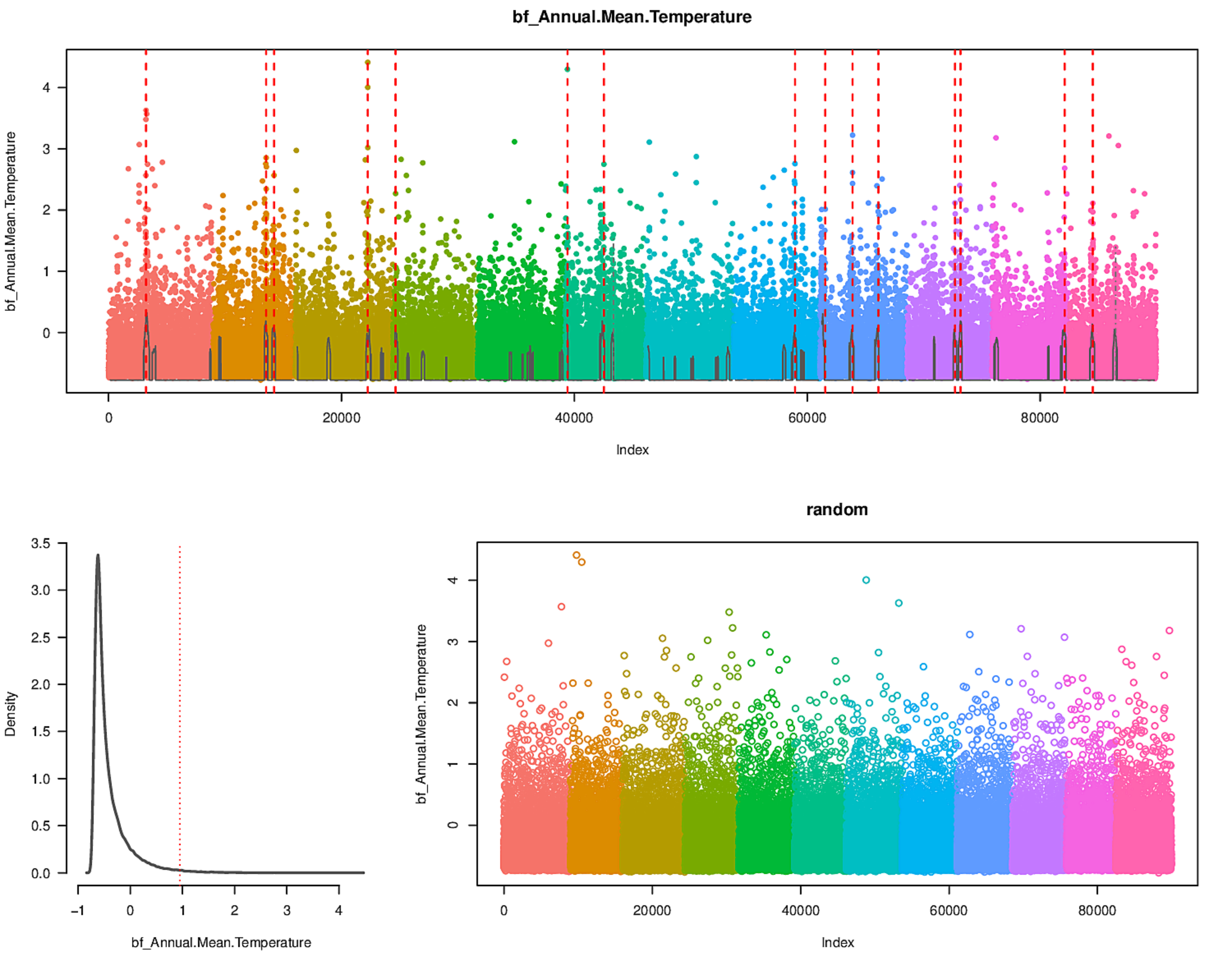
bf_Annual.Mean.Temperature
bf_Annual.Mean.Temperature
Index
0
20000
40000
60000
80000
0
1
2
3
4
Density
3.5
3.0
2.5
2.0
1.5
1.0
0.5
0.0
-1
0
1
2
3
4
bf_Annual.Mean.Temperature
random
bf_Annual.Mean.Temperature
Index

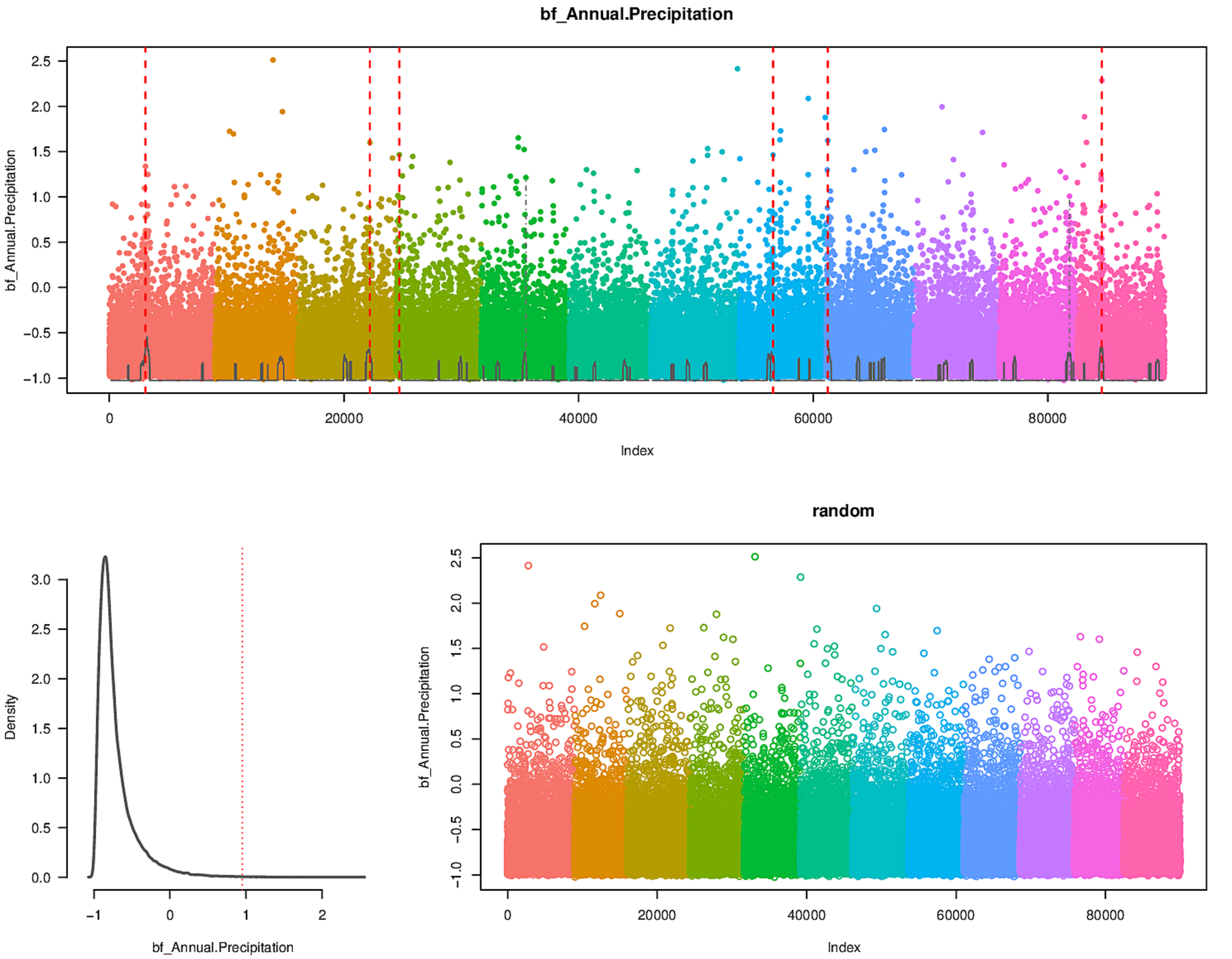
bf_Annual.Precipitation
bf_Annual.Precipitation
Index
Density
bf_Annual.Precipitation
random
bf_Annual.Precipitation
Index

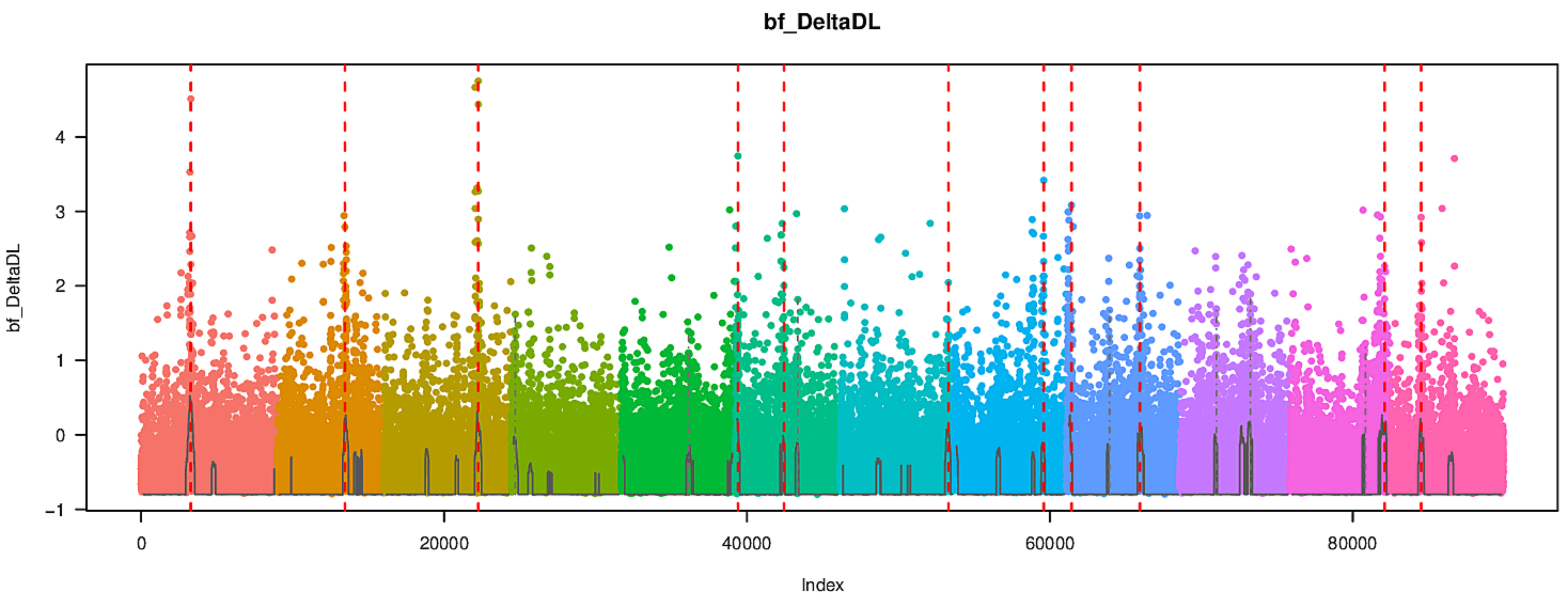
bf_DeltaDL
bf_DeltaDL
Index
0
20000
40000
60000
80000
-1
0
1
2
3
4

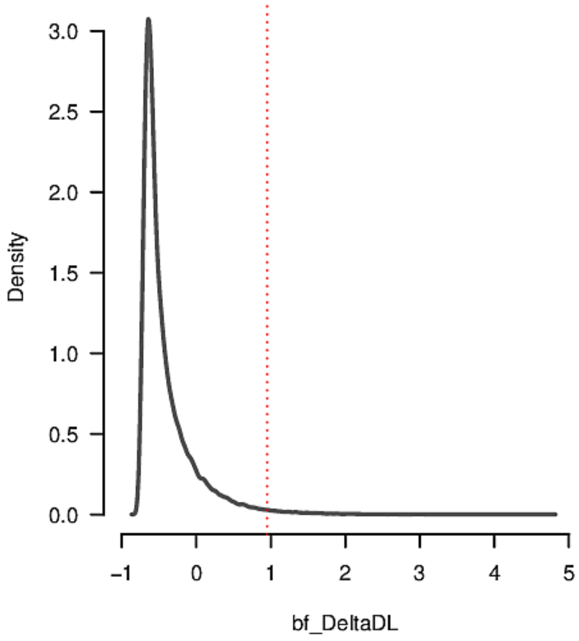
Density
bf_DeltaDL
0.0
0.5
1.0
1.5
2.0
2.5
3.0
-1
0
1
2
3
4
5

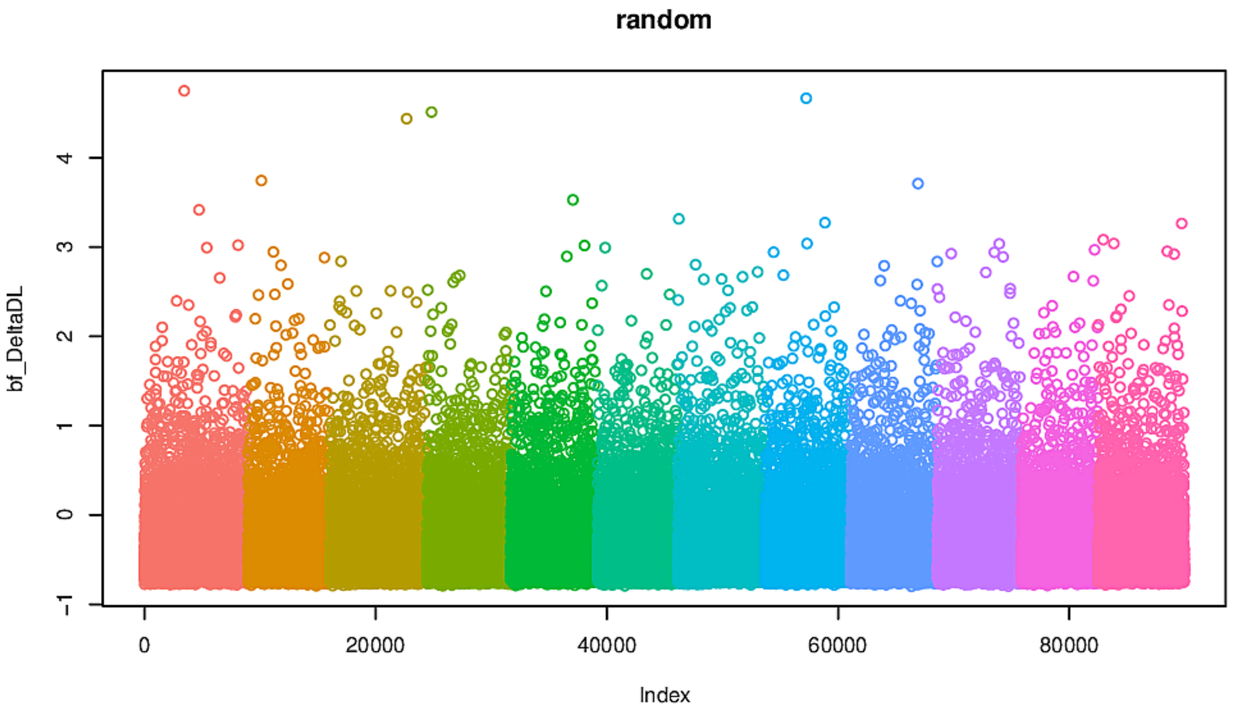
random
bf_DeltaDL
Index
0
20000
40000
60000
80000
-1
0
1
2
3
4

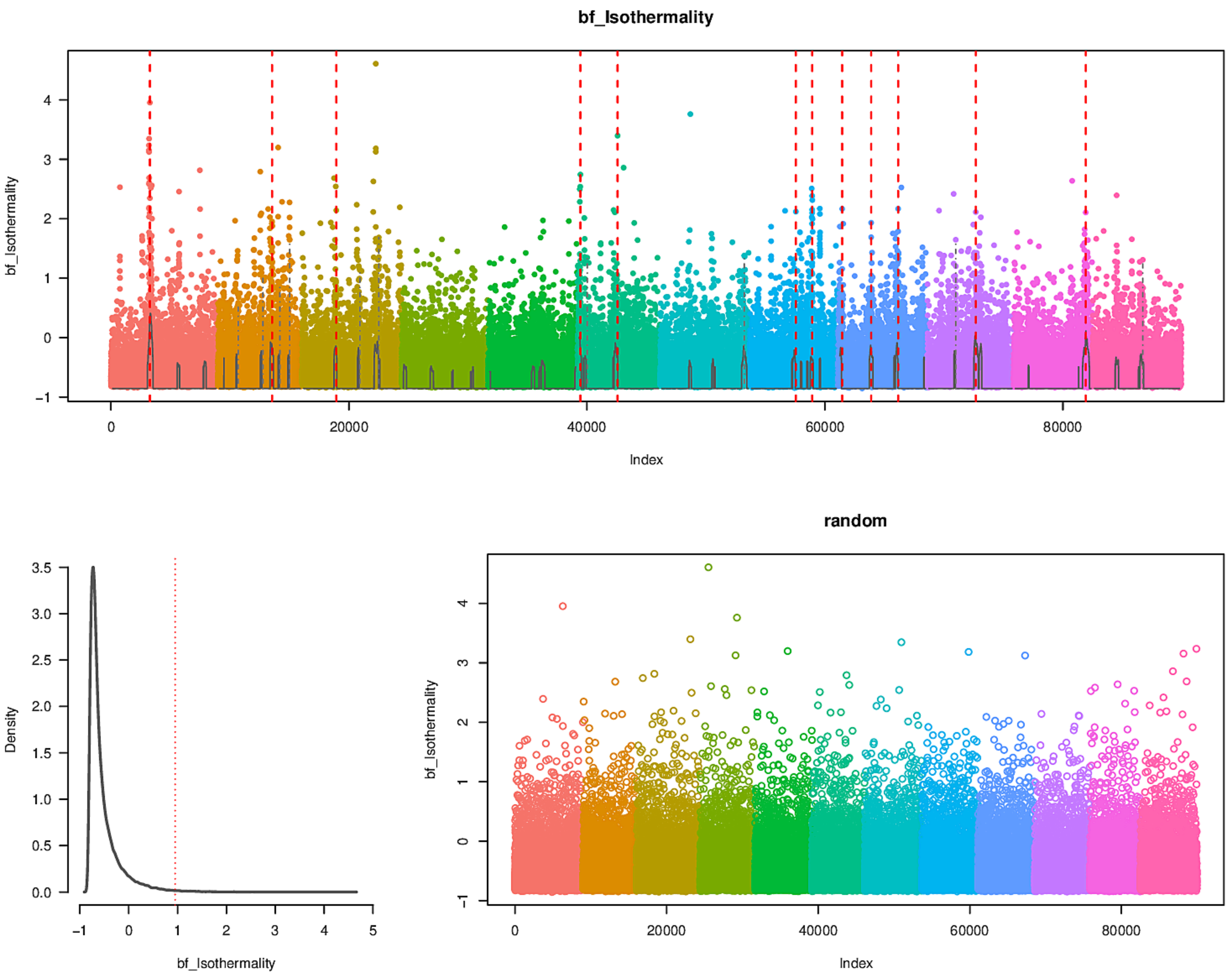
bf_Isothermality
bf_Isothermality
Index
0
20000
40000
60000
80000
-1
0
1
2
3
4
Density
bf_Isothermality
-1
0
1
2
3
4
5
0.0
0.5
1.0
1.5
2.0
2.5
3.0
3.5
random
bf_Isothermality
Index

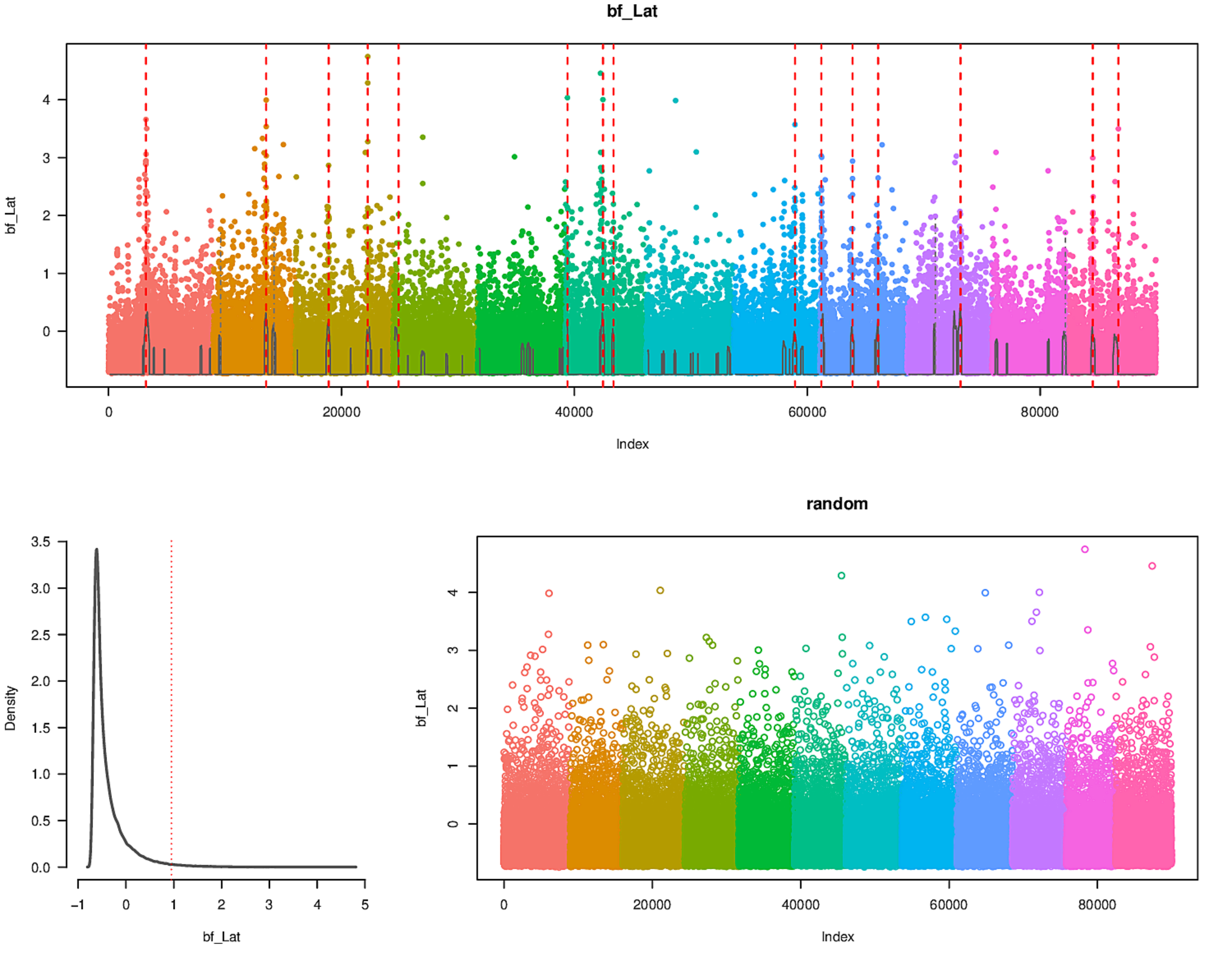
bf_Lat
Index
bf_Lat
0
20000
40000
60000
80000
0
1
2
3
4
Density
3.5
3.0
2.5
2.0
1.5
1.0
0.5
0.0
-1
5
random

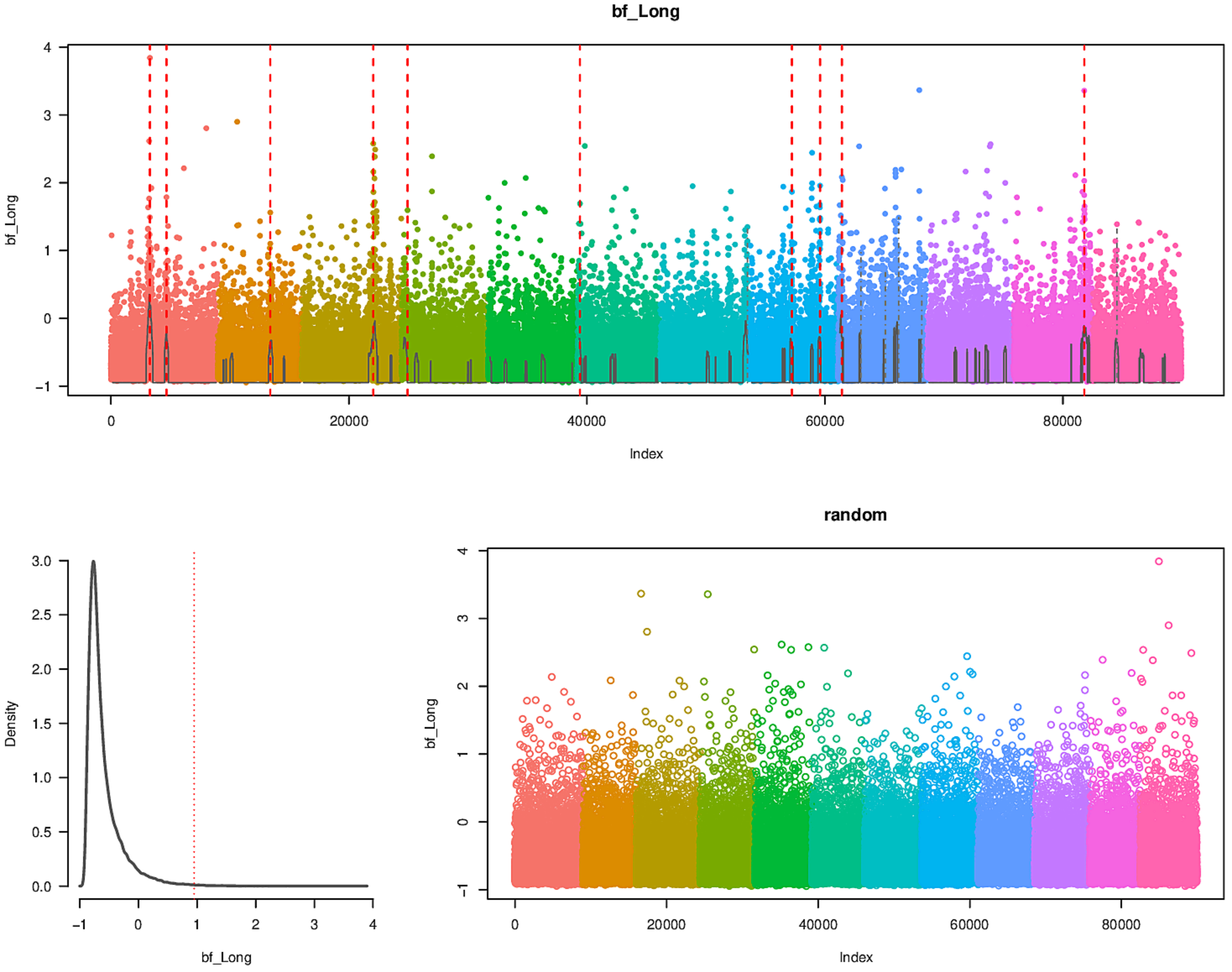

bf_Long
bf_Long
Index
0
20000
40000
60000
80000
-1
0
1
2
3
4
Density
bf_Long
0.0
0.5
1.0
1.5
2.0
2.5
3.0
-1
0
1
2
3
4
random
bf_Long
Index
0
20000
40000
60000
80000
-1
0
1
2
3
4

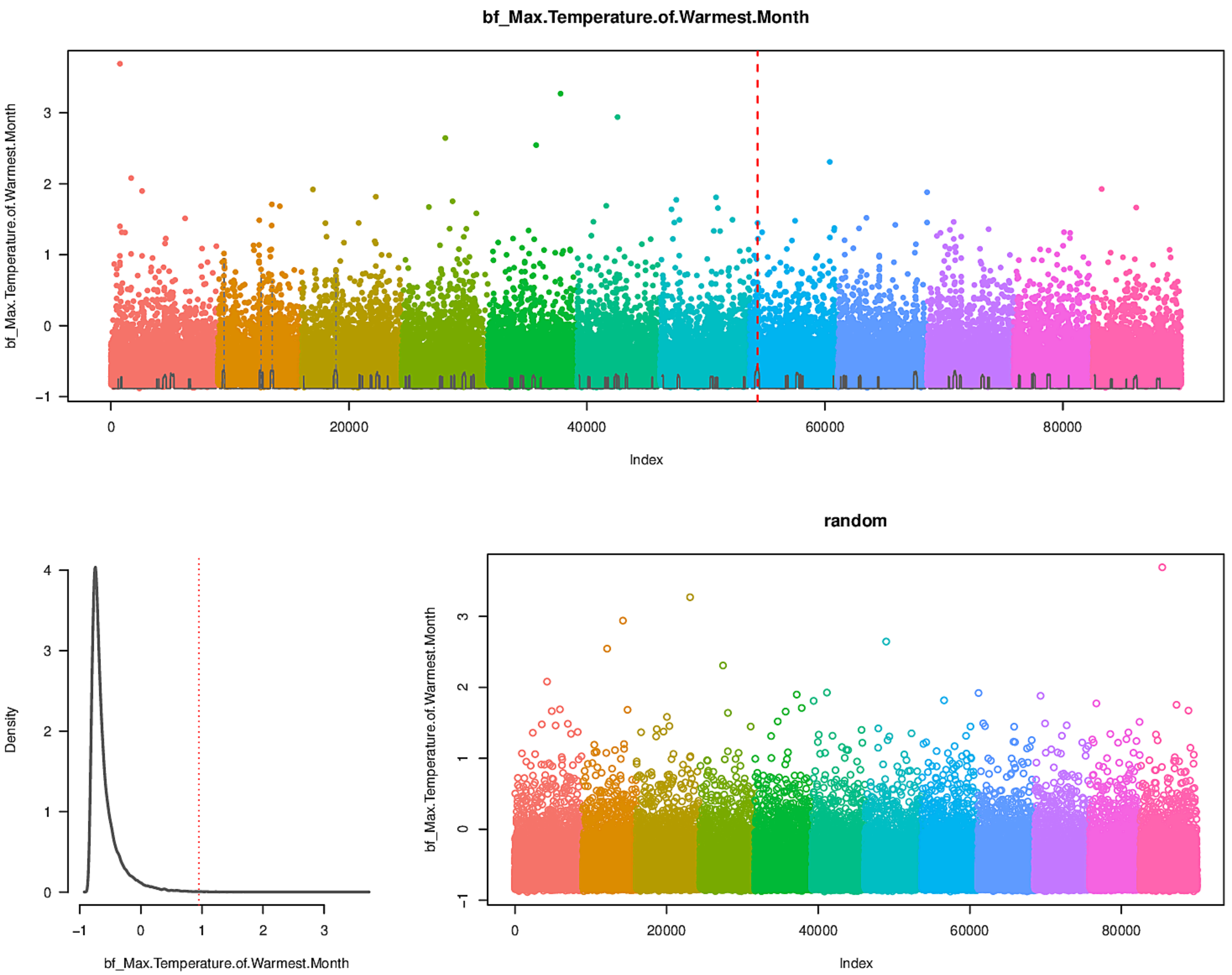
bf_Max.Temperature.of.Warmest.Month
bf_Max.Temperature.of.Warmest.Month
Index
0
20000
40000
60000
80000
−1
0
1
2
3
Density
bf_Max.Temperature.of.Warmest.Month
random
bf_Max.Temperature.of.Warmest.Month
Index

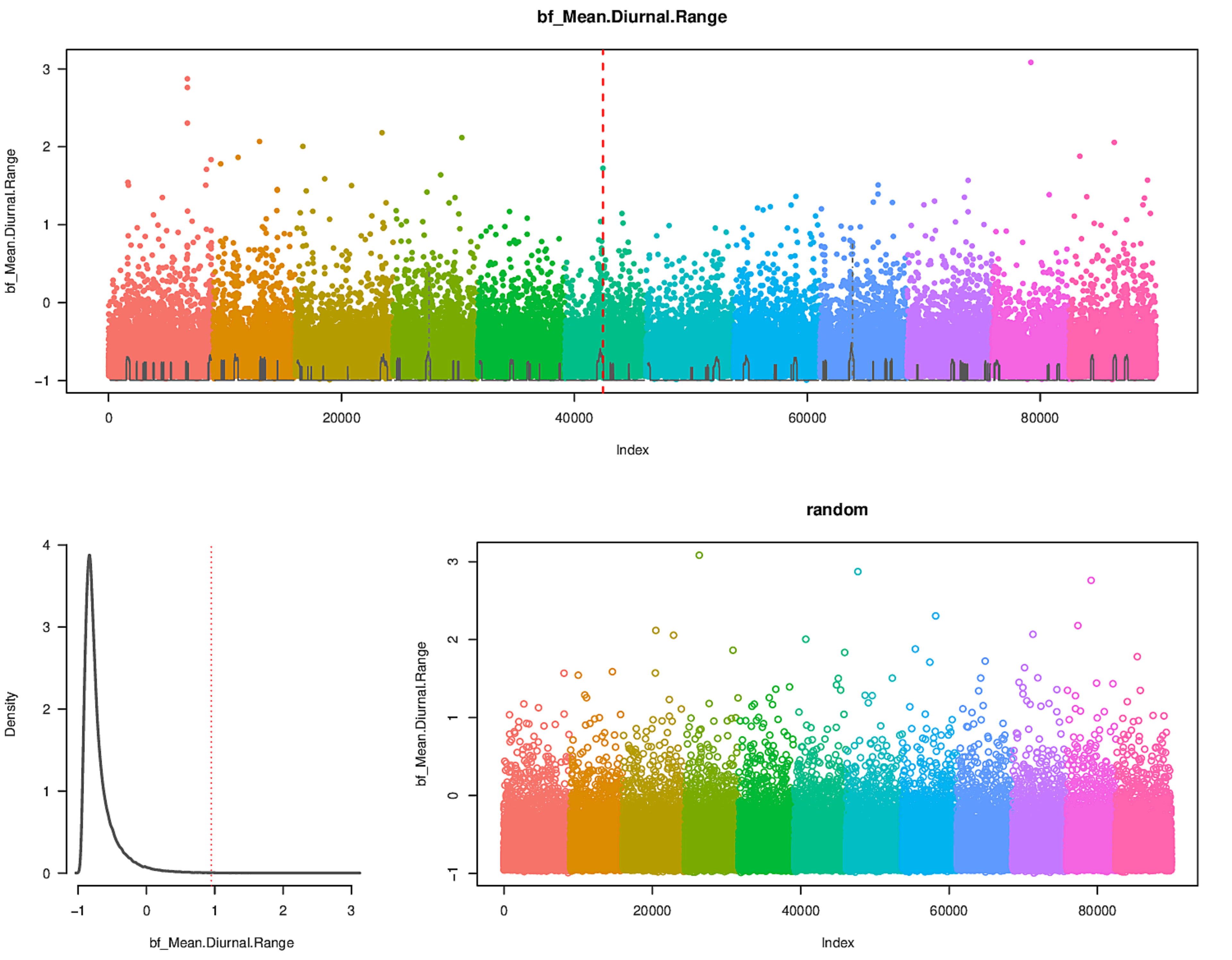
bf_Mean.Diurnal.Range
bf_Mean.Diurnal.Range
Index
Density
bf_Mean.Diurnal.Range
random
bf_Mean.Diurnal.Range
Index

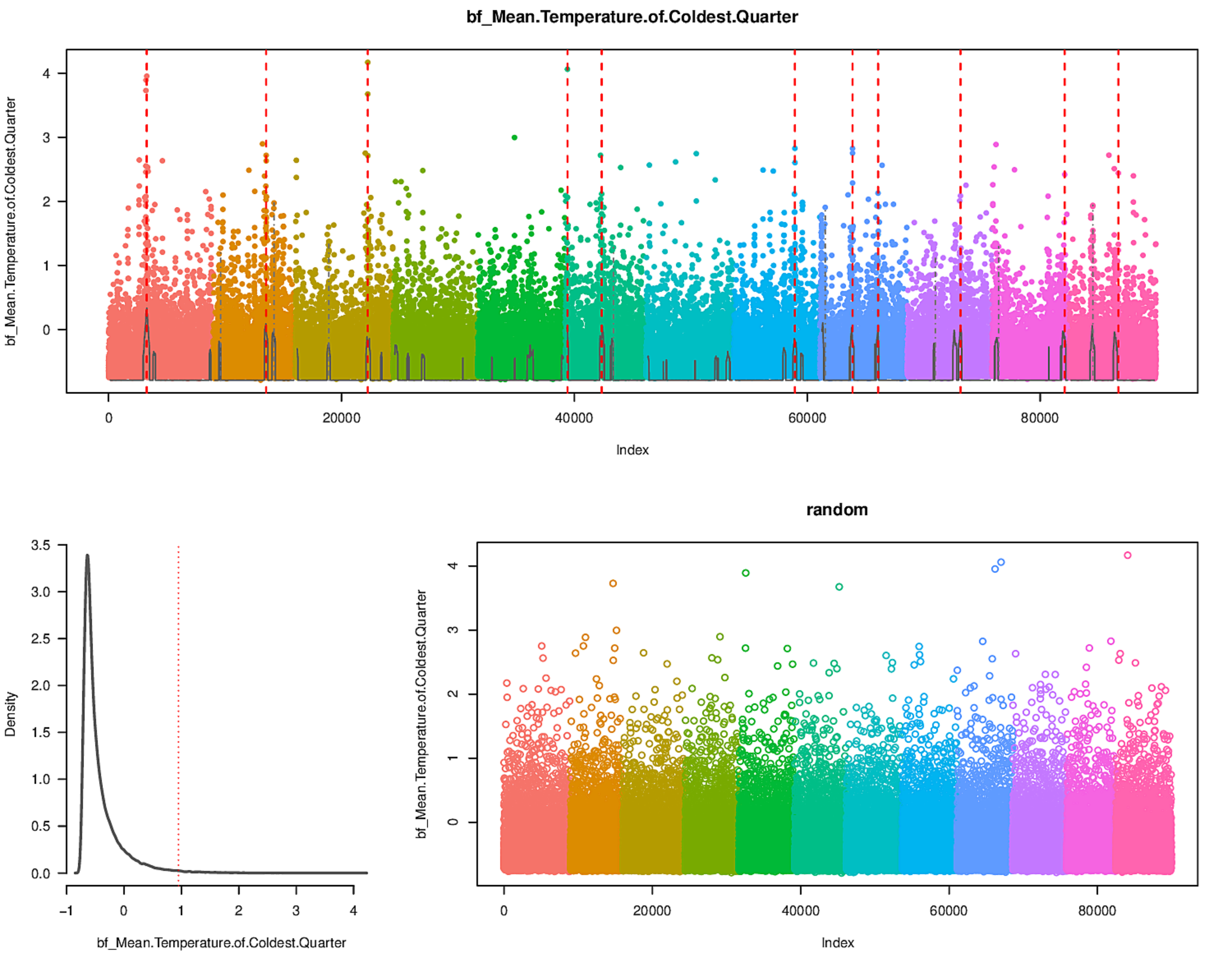

bf_Mean.Temperature.of.Coldest.Quarter
Index
Density
random

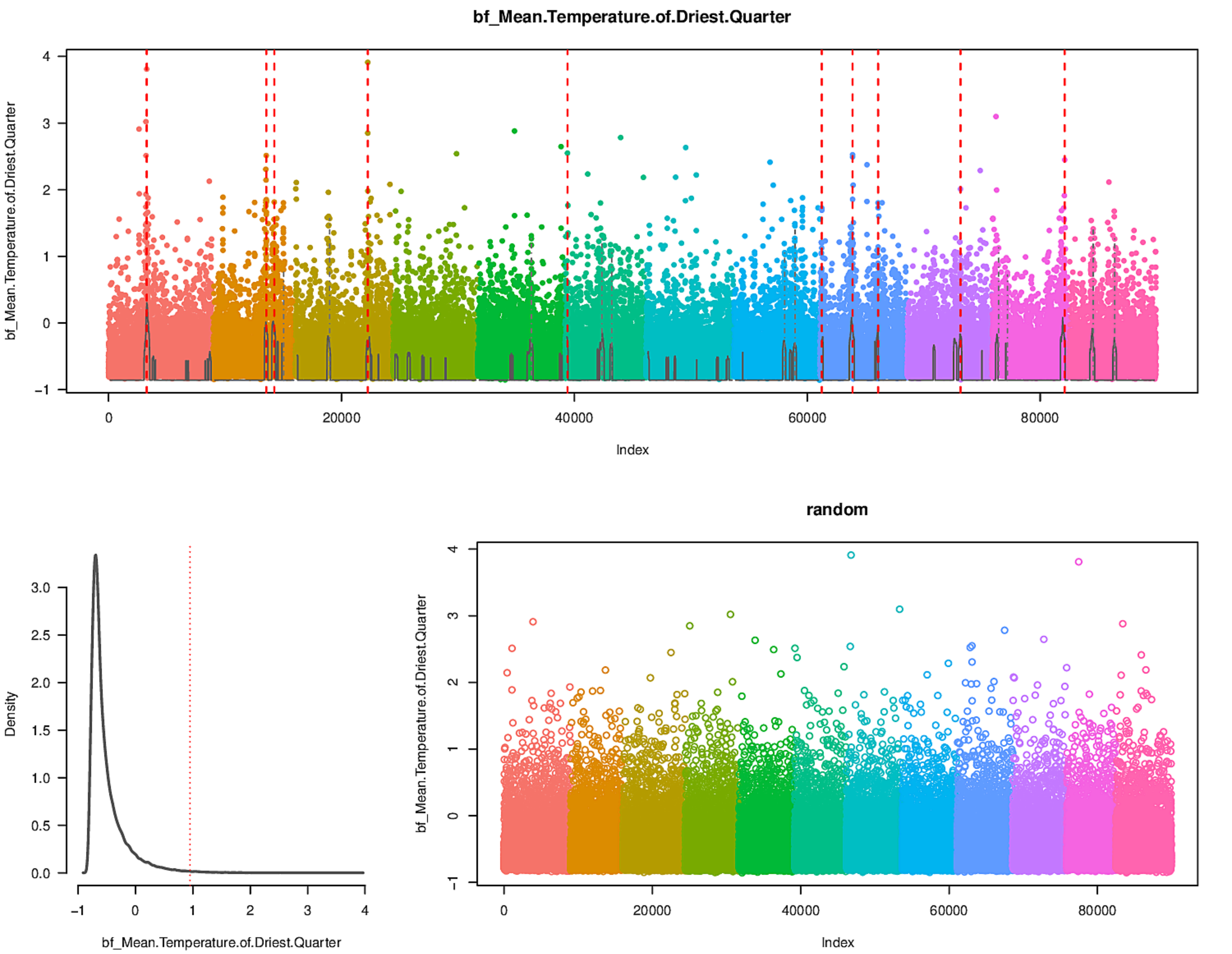

bf_Mean.Temperature.of.Driest.Quarter
bf_Mean.Temperature.of.Driest.Quarter
Index
0
20000
40000
60000
80000
−1
0
1
2
3
4
Density
bf_Mean.Temperature.of.Driest.Quarter
0.0
0.5
1.0
1.5
2.0
2.5
3.0
random
bf_Mean.Temperature.of.Driest.Quarter
Index

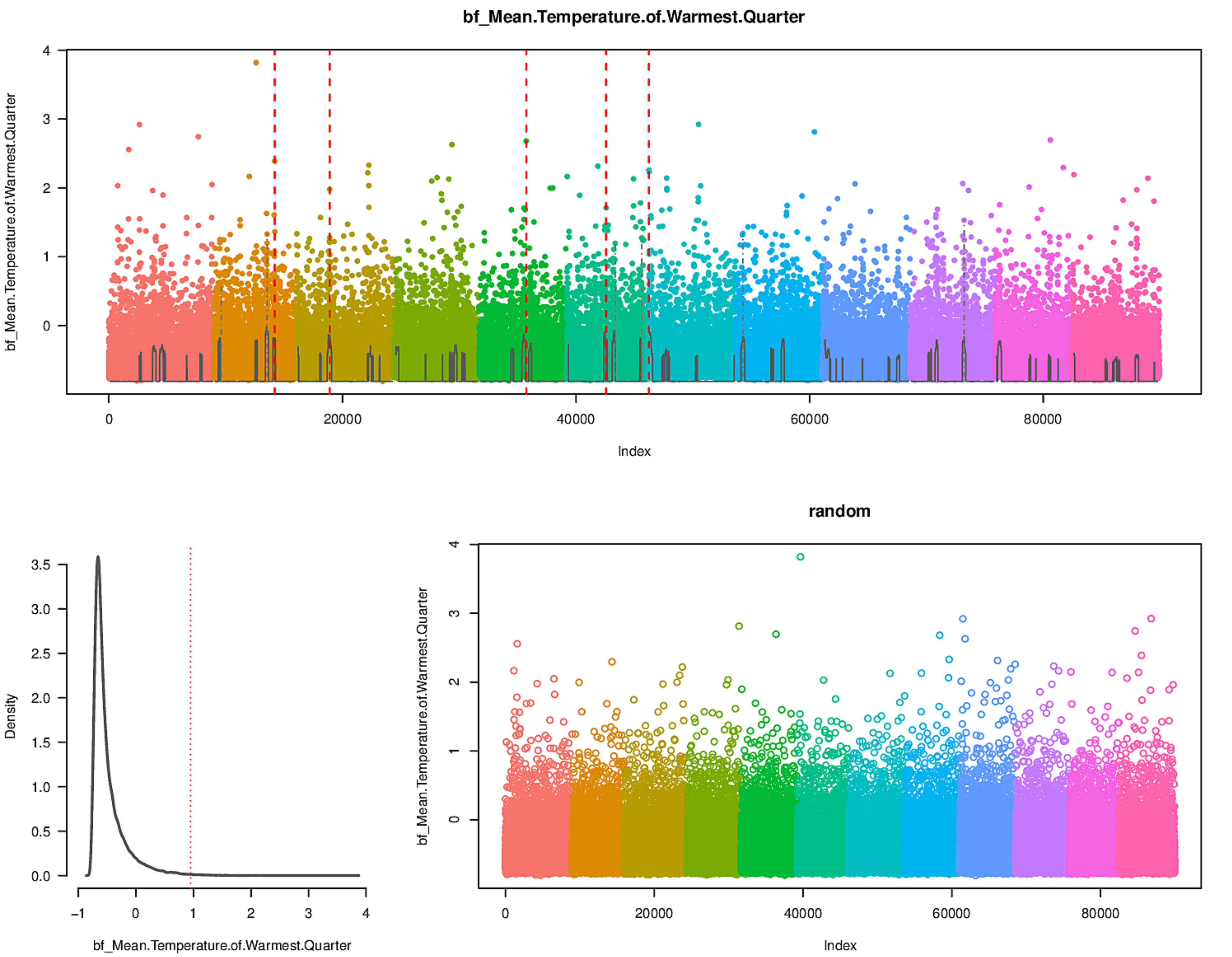
bf_Mean.Temperature.of.Warmest.Quarter
Index
Density
random

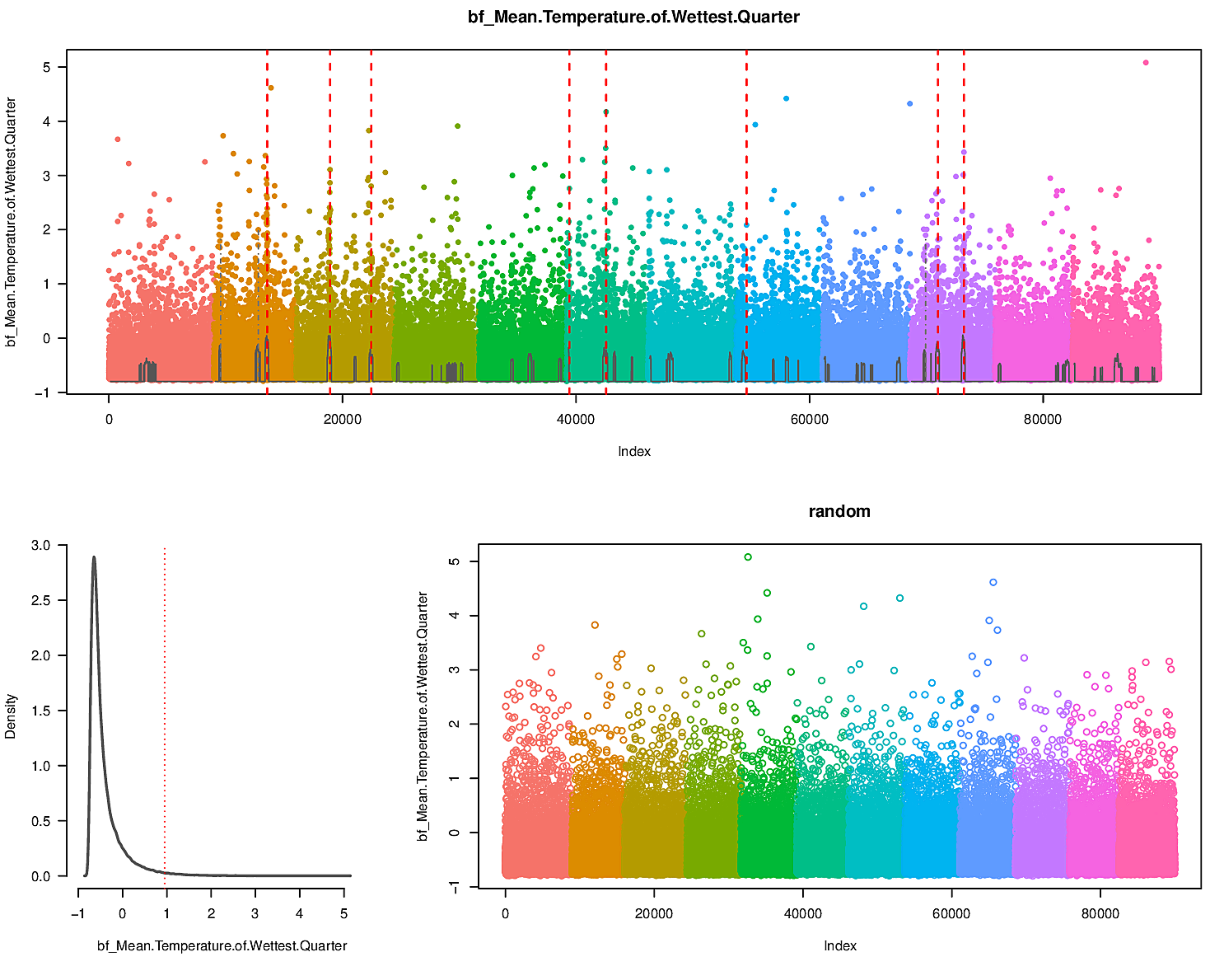
bf_Mean.Temperature.of.Wettest.Quarter
Index
Density
random
bf_Mean.Temperature.of.Wettest.Quarter
Index

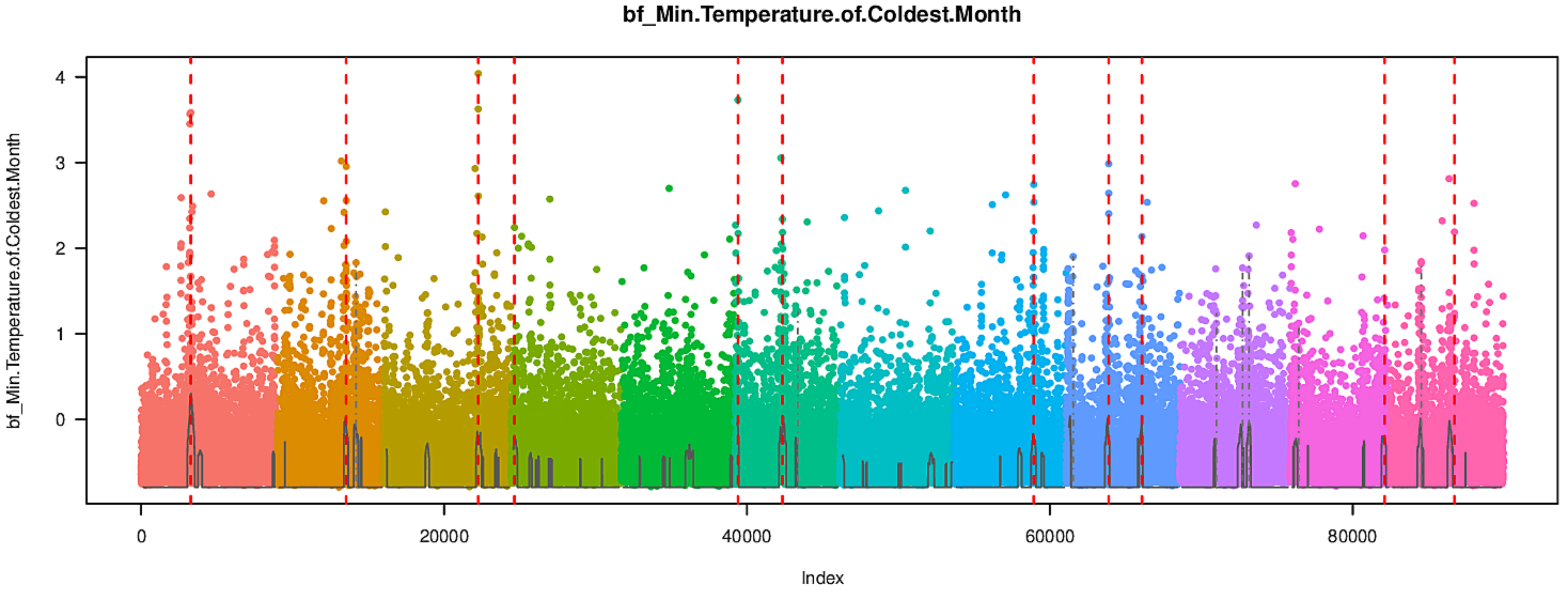
bf_Min.Temperature.of.Coldest.Month
bf_Min.Temperature.of.Coldest.Month
Index
0
20000
40000
60000
80000
0
1
2
3
4

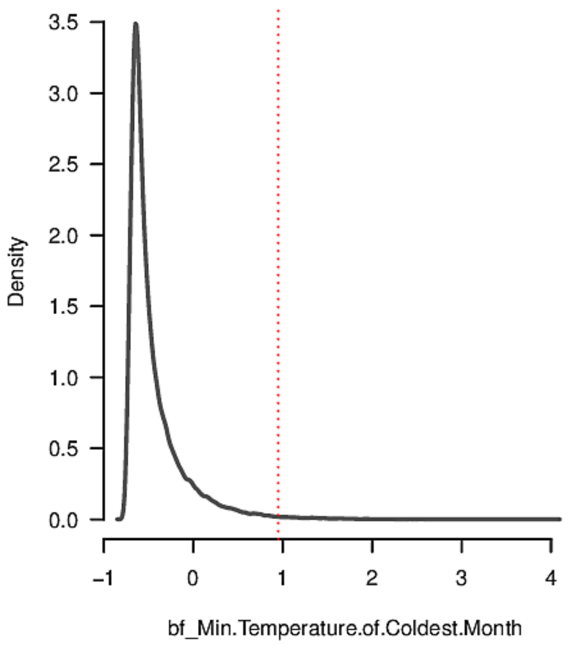
Density
bf_Min.Temperature.of.Coldest.Month
0.0
0.5
1.0
1.5
2.0
2.5
3.0
3.5
-1
0
1
2
3
4

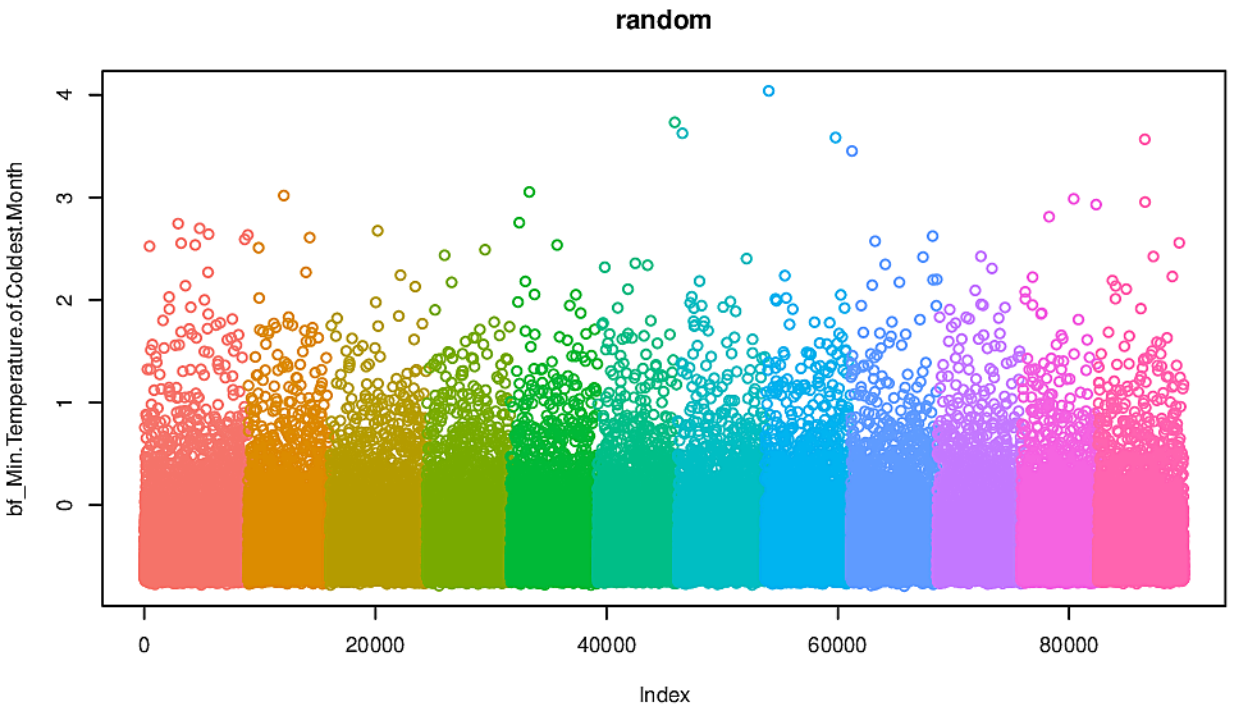
random
bf_Min.Temperature.of.Coldest.Month
Index
0
20000
40000
60000
80000
0
1
2
3
4

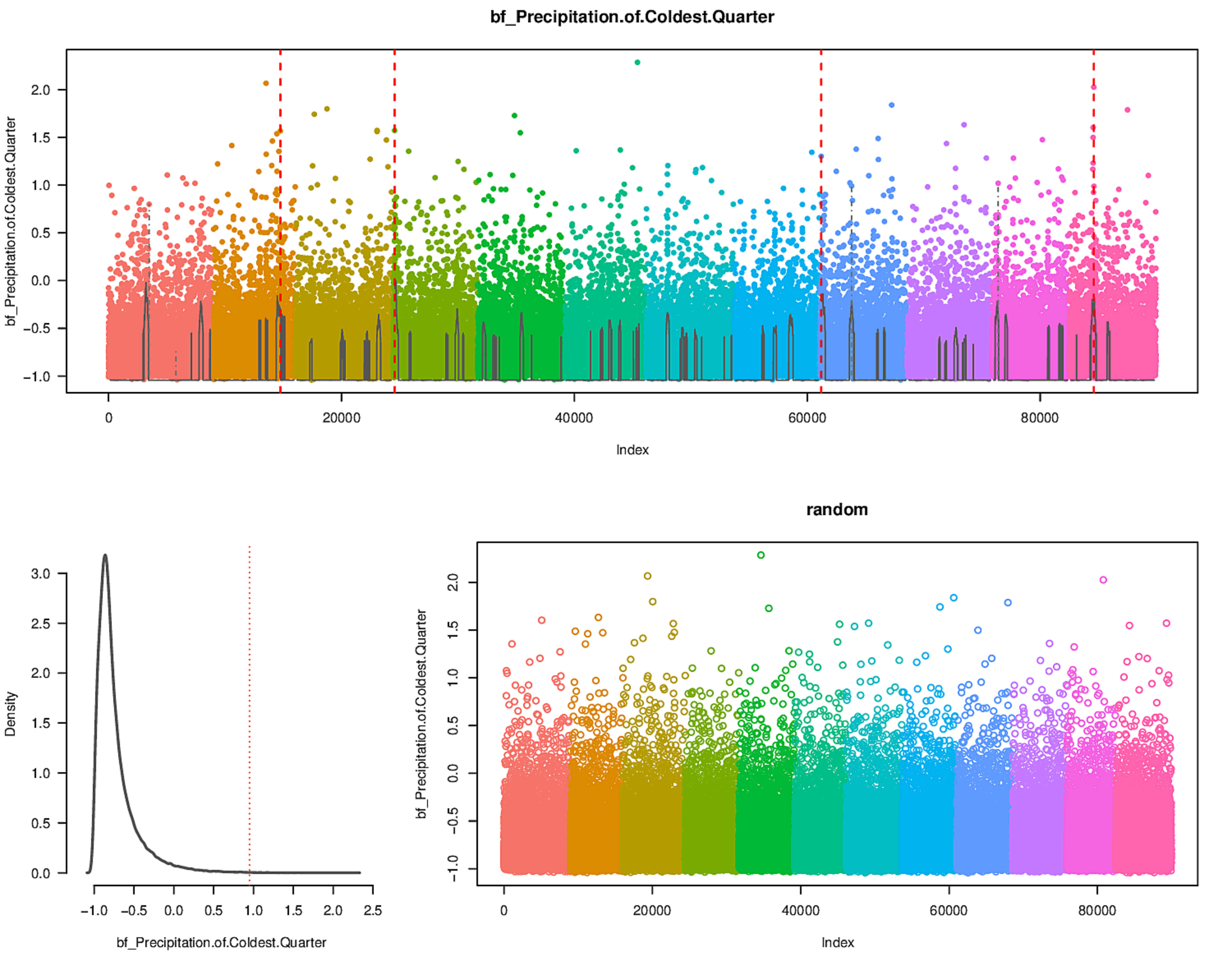
bf_Precipitation.of.Coldest.Quarter
bf_Precipitation.of.Coldest.Quarter
Index
Density
bf_Precipitation.of.Coldest.Quarter
random
bf_Precipitation.of.Coldest.Quarter
Index

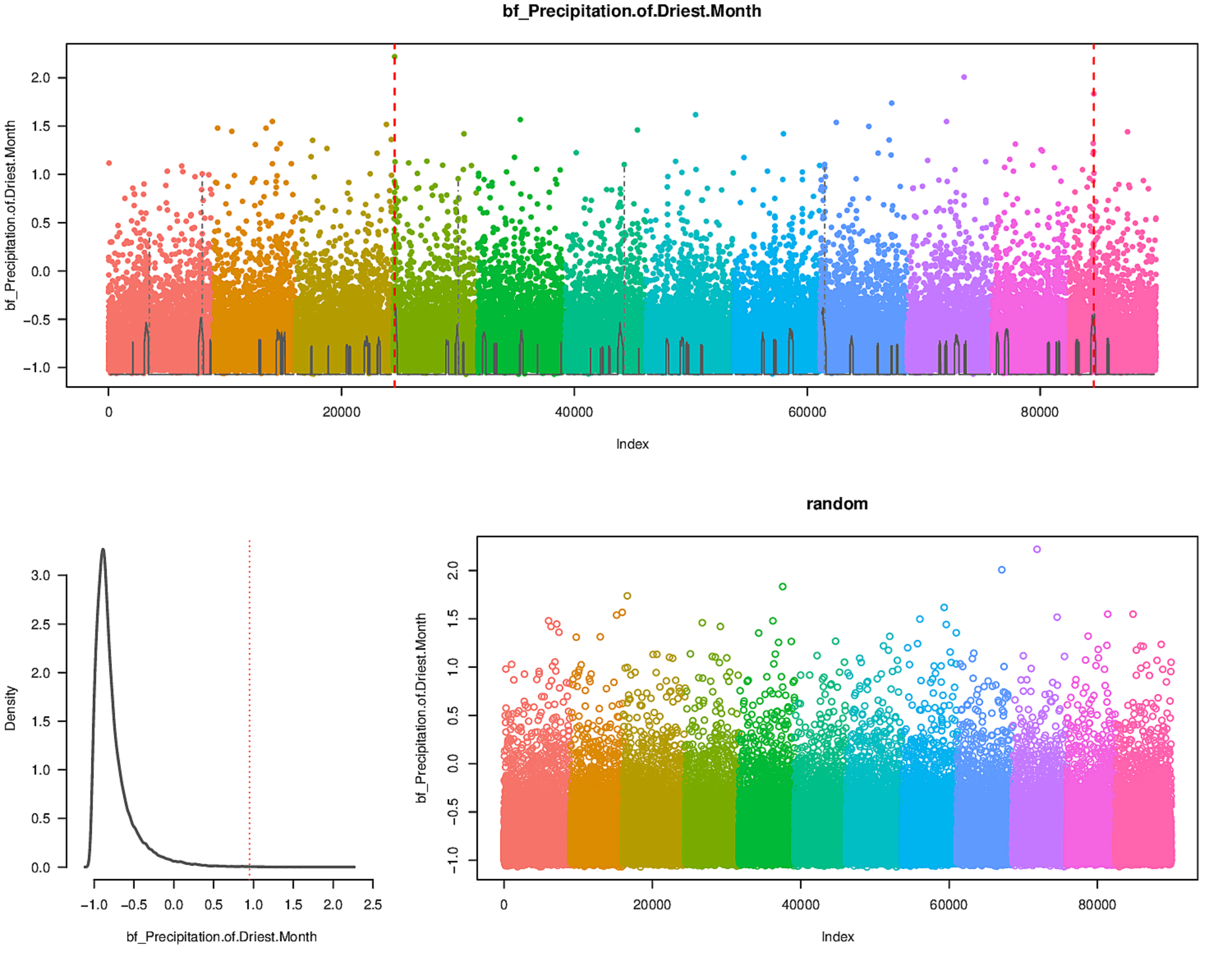
bf_Precipitation.of.Driest.Month
bf_Precipitation.of.Driest.Month
Index
Density
bf_Precipitation.of.Driest.Month
random
bf_Precipitation.of.Driest.Month
Index

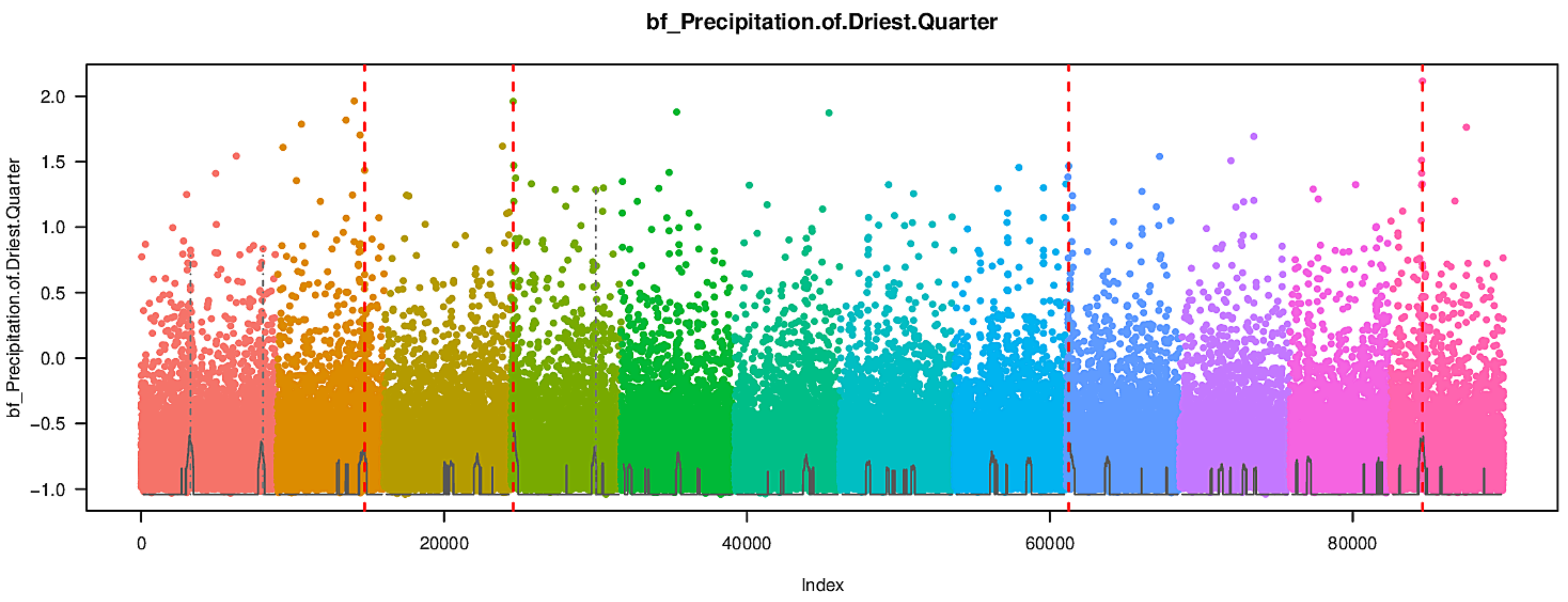

bf_Precipitation.of.Driest.Quarter
bf_Precipitation.of.Driest.Quarter
Index


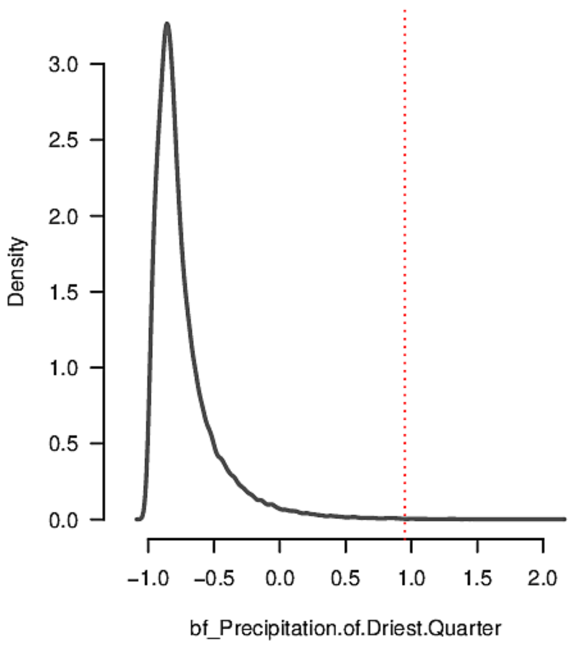

Density
bf_Precipitation.of.Driest.Quarter


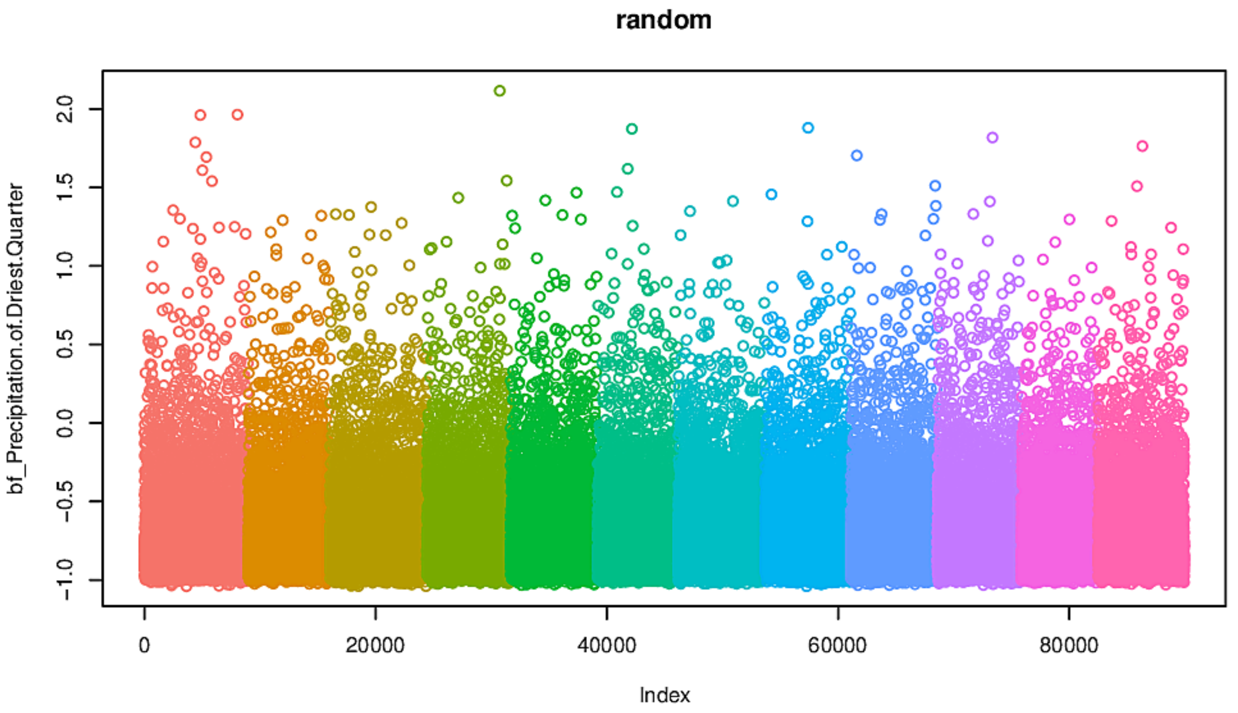

random
bf_Precipitation.of.Driest.Quarter
Index

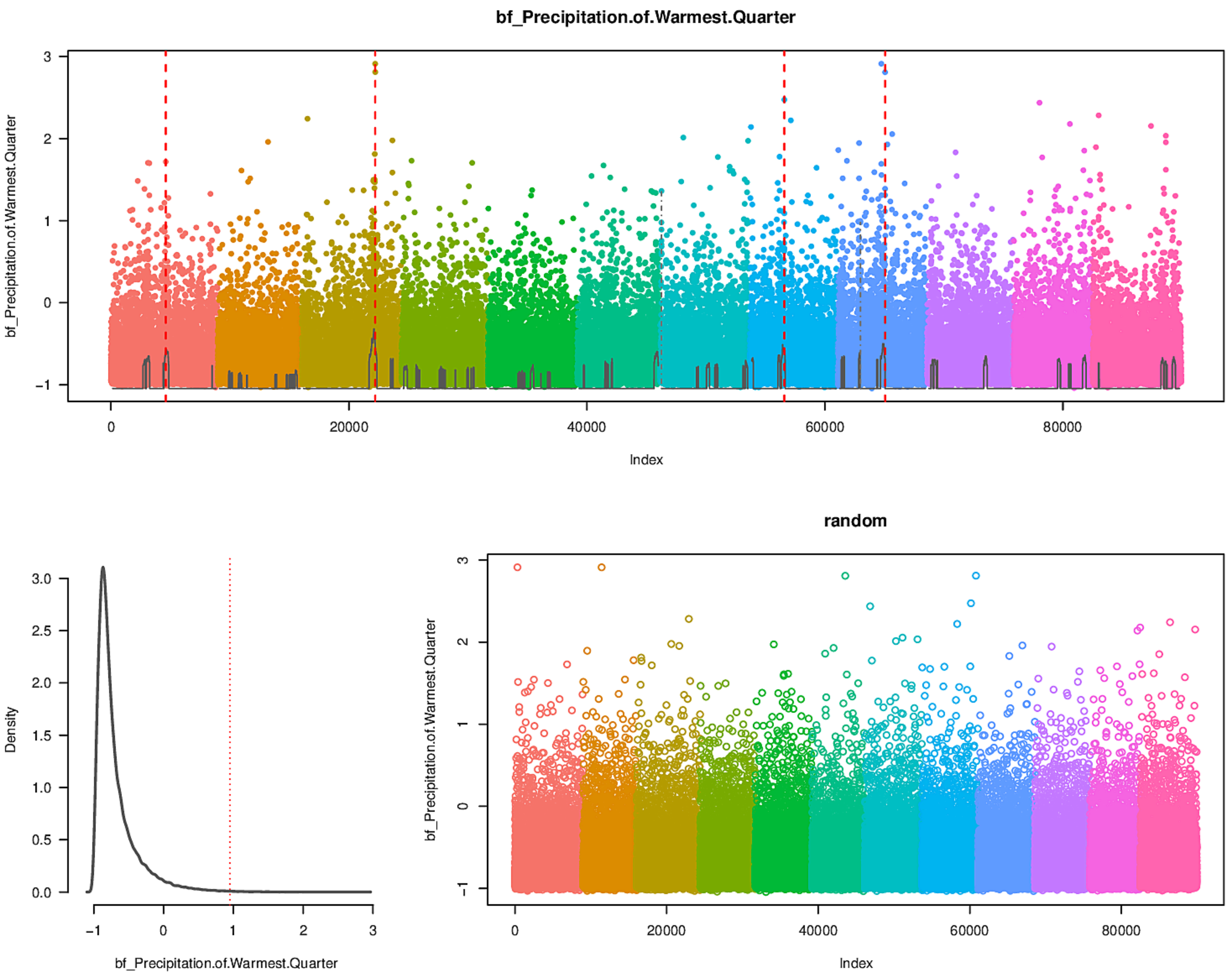
bf_Precipitation.of.Warmest.Quarter
bf_Precipitation.of.Warmest.Quarter
Index
0
20000
40000
60000
80000
-1
0
1
2
3
Density
0.0
0.5
1.0
1.5
2.0
2.5
3.0
bf_Precipitation.of.Warmest.Quarter
random
bf_Precipitation.of.Warmest.Quarter
Index

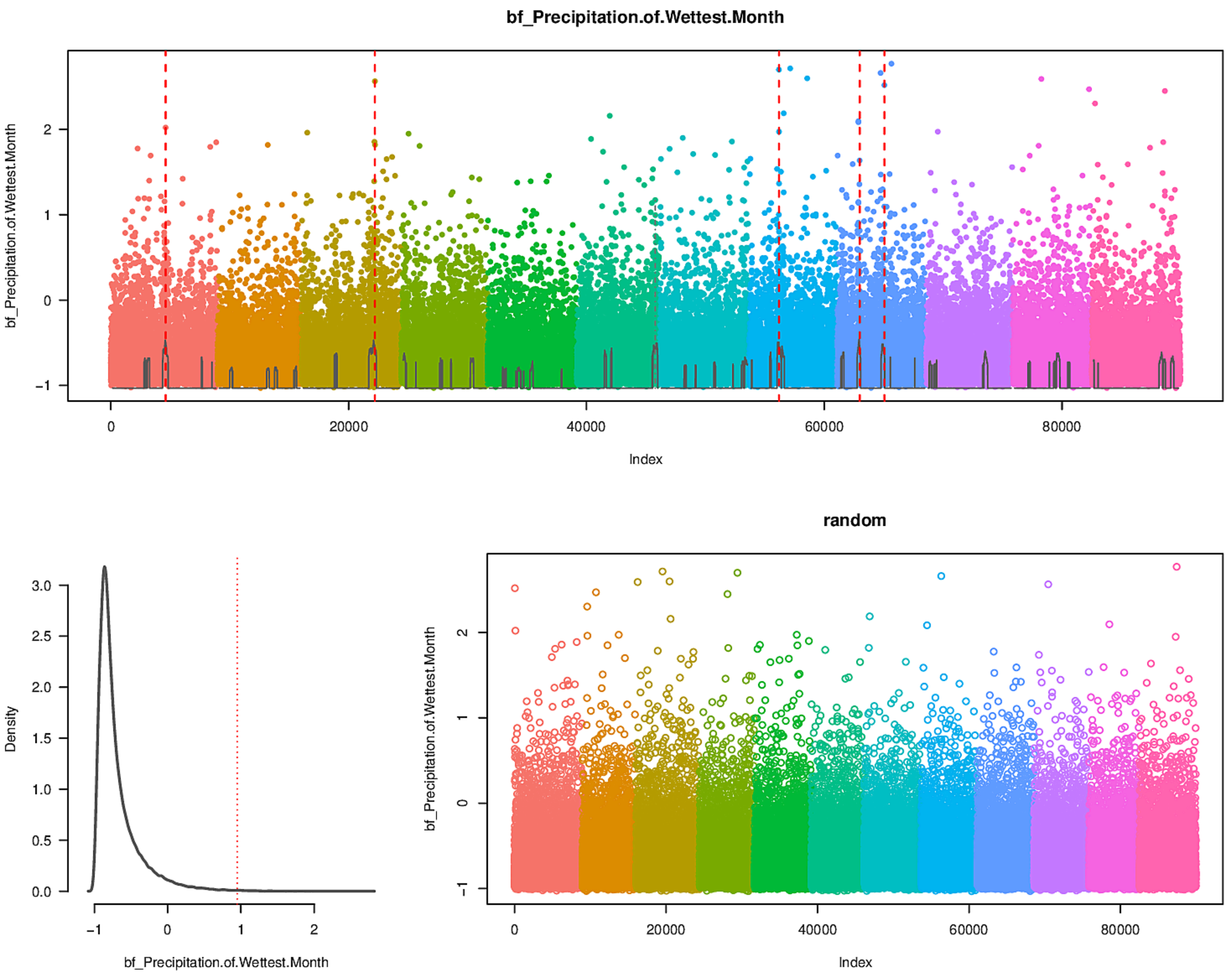
bf_Precipitation.of.Wettest.Month
bf_Precipitation.of.Wettest.Month
Index
0
20000
40000
60000
80000
−1
0
1
2
Density
3.0
2.5
2.0
1.5
1.0
0.5
0.0
bf_Precipitation.of.Wettest.Month
random
bf_Precipitation.of.Wettest.Month
Index

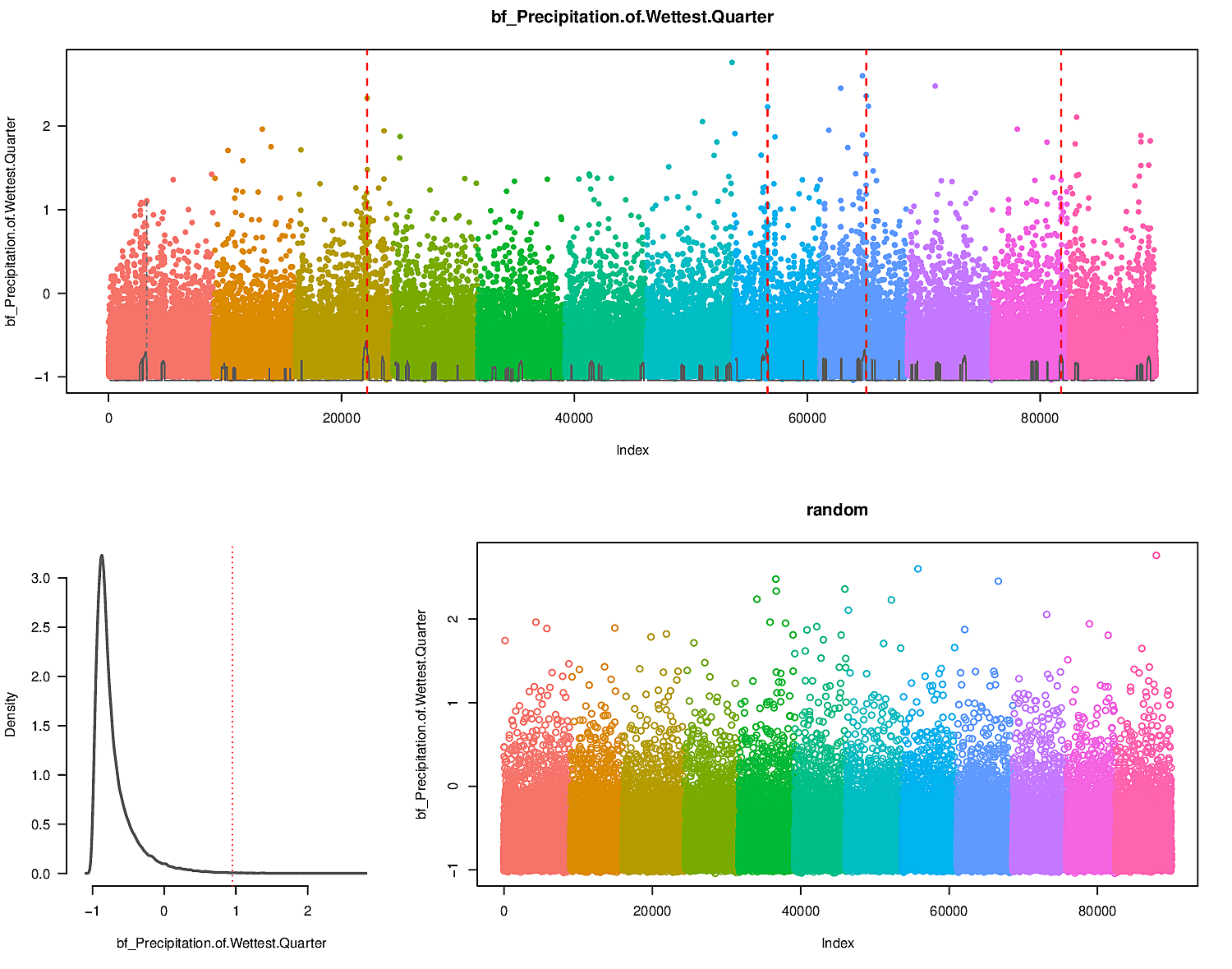
bf_Precipitation.of.Wettest.Quarter
bf_Precipitation.of.Wettest.Quarter
Index
0
20000
40000
60000
80000
2
1
0
−1
Density
3.0
2.5
2.0
1.5
1.0
0.5
0.0
−1
0
1
2
bf_Precipitation.of.Wettest.Quarter
random
bf_Precipitation.of.Wettest.Quarter
Index

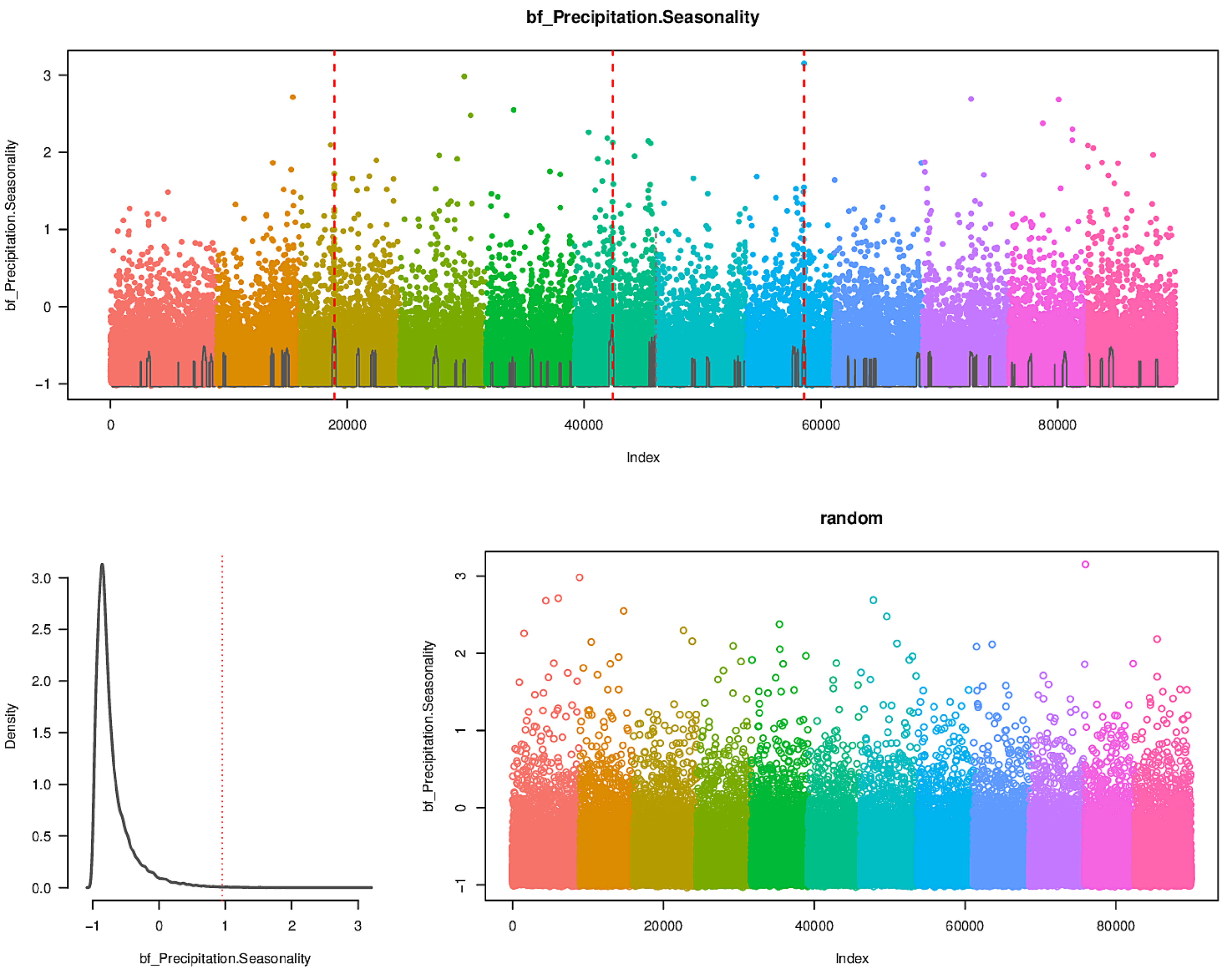

bf_Precipitation.Seasonality
Index
Density
random

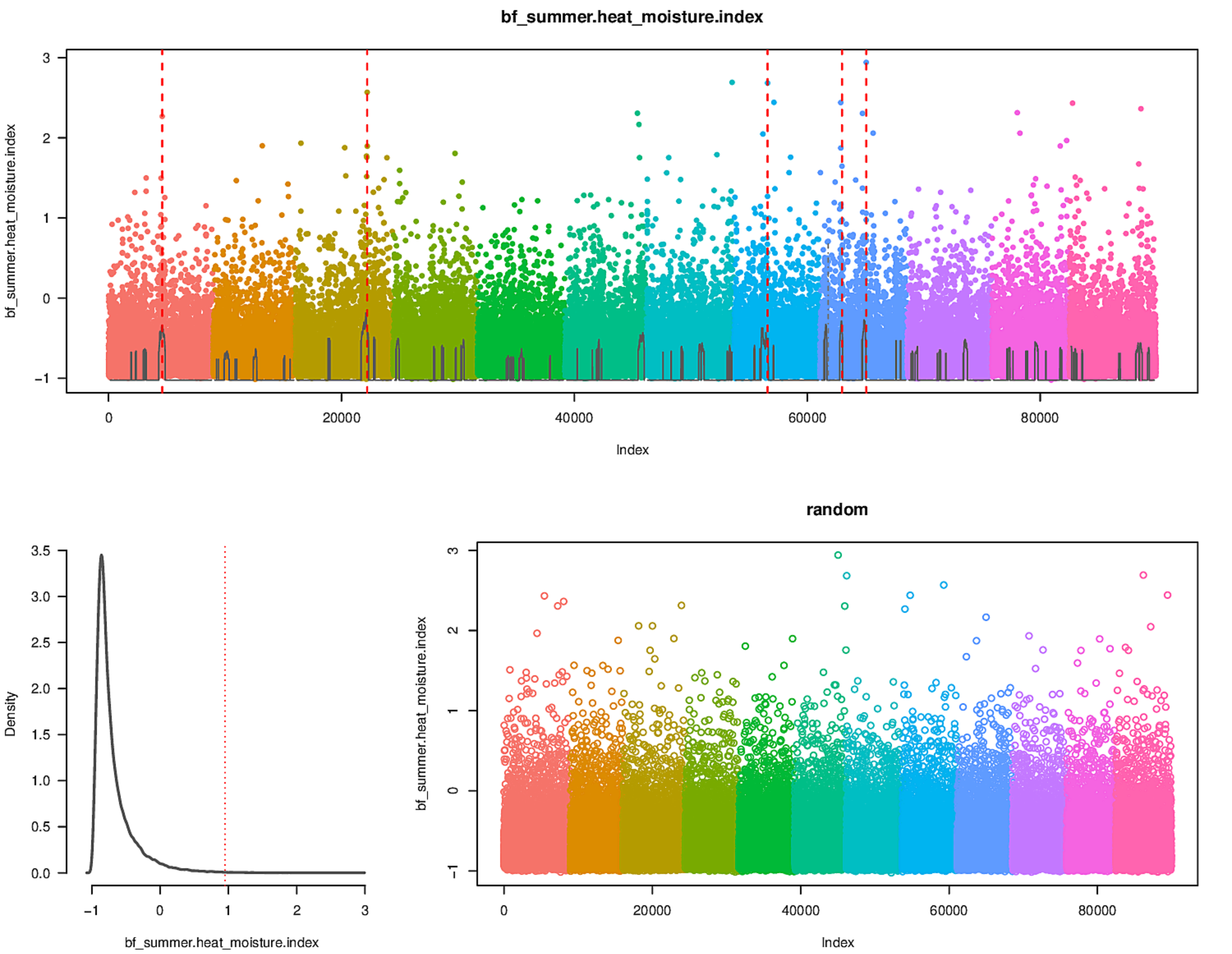
bf_summer.heat_moisture.index
bf_summer.heat_moisture.index
Index
0
20000
40000
60000
80000
-1
0
1
2
3
Density
bf_summer.heat_moisture.index
3.5
3.0
2.5
2.0
1.5
1.0
0.5
0.0
-1
0
1
2
3
random
bf_summer.heat_moisture.index
Index
0
20000
40000
60000
80000
-1
0
1
2
3

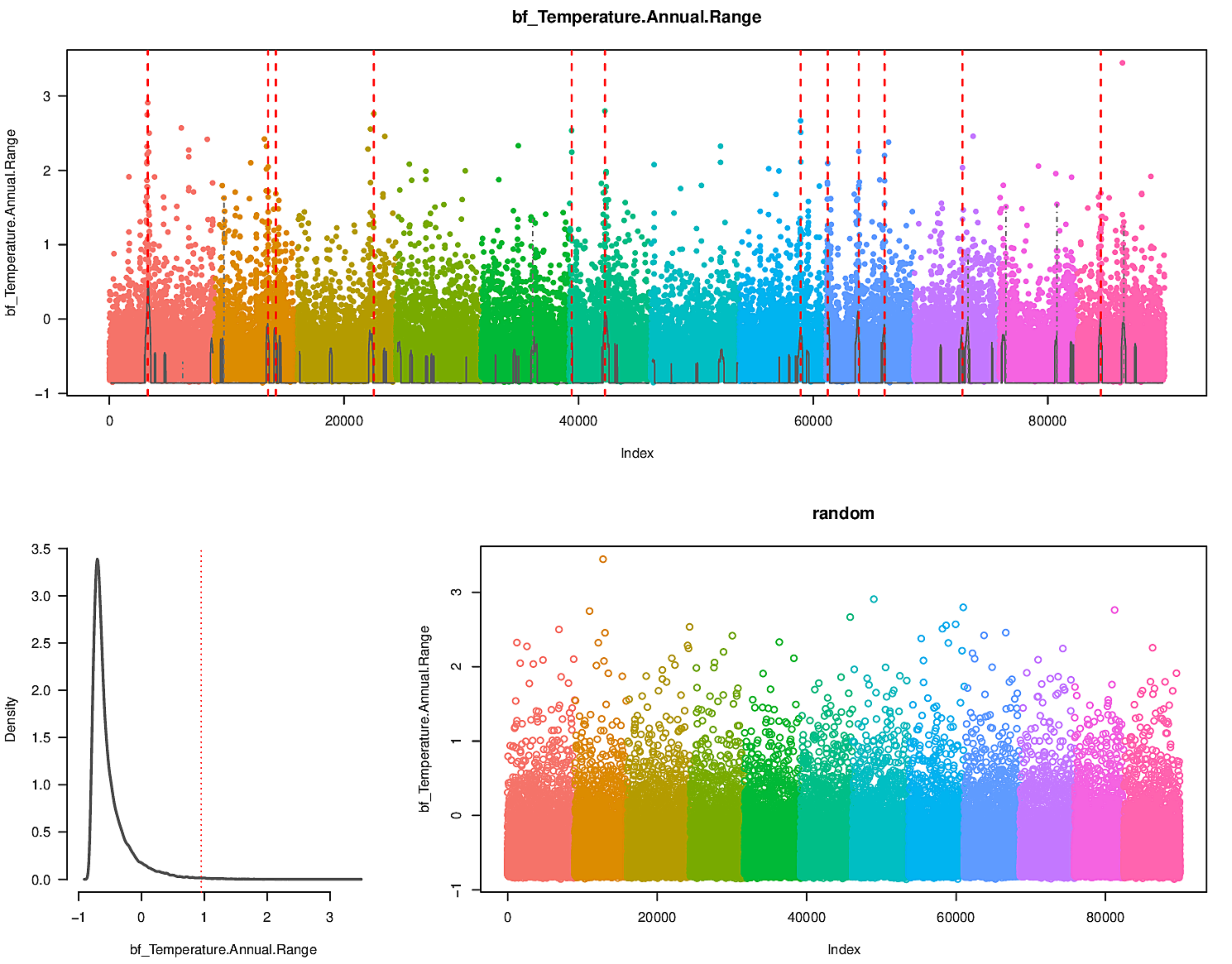

bf_Temperature.Annual.Range
bf_Temperature.Annual.Range
Index
0
20000
40000
60000
80000
-1
0
1
2
3
Density
0.0
0.5
1.0
1.5
2.0
2.5
3.0
3.5
random

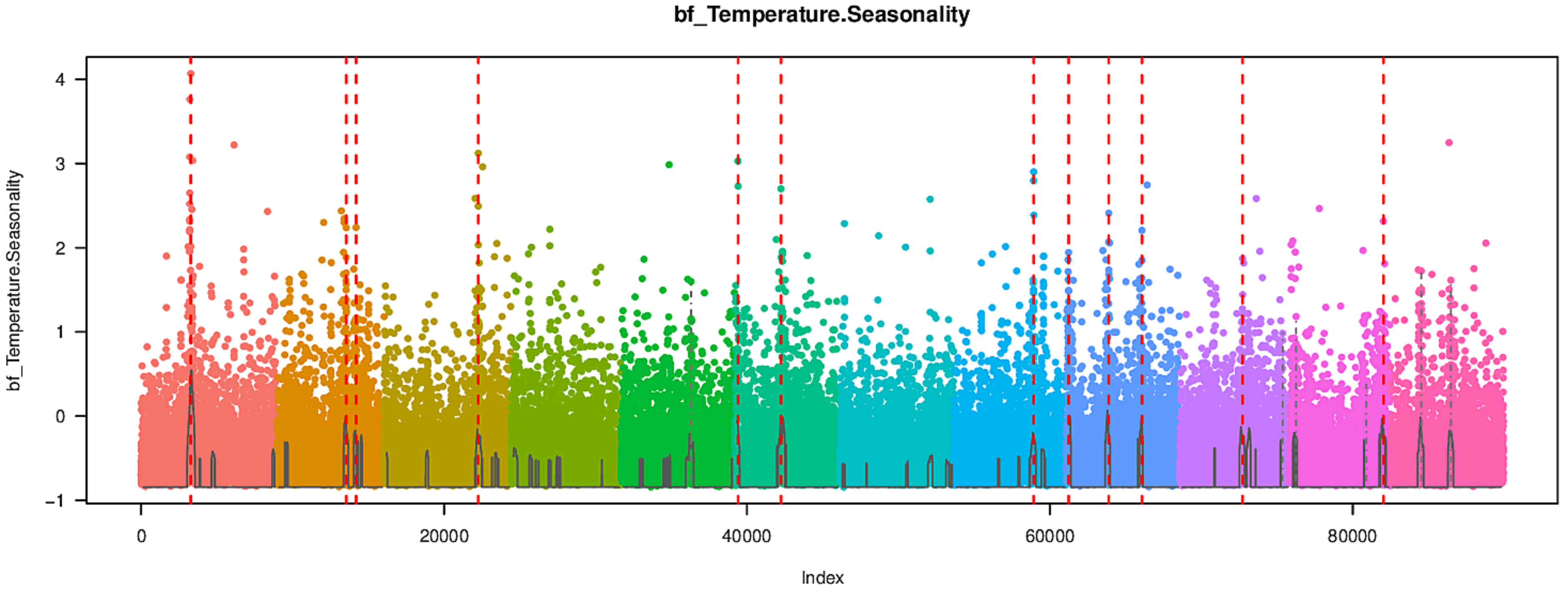
bf_Temperature.Seasonality
bf_Temperature.Seasonality
Index
0
20000
40000
60000
80000
-1
0
1
2
3
4

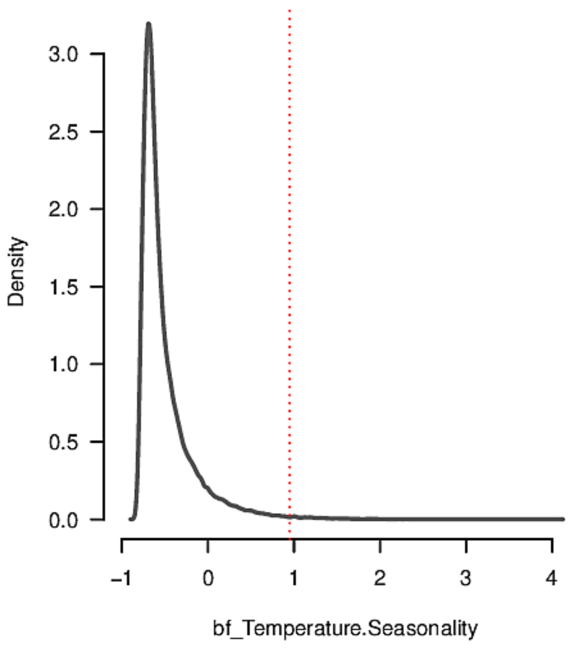
Density
bf_Temperature.Seasonality
0.0
0.5
1.0
1.5
2.0
2.5
3.0
-1
0
1
2
3
4

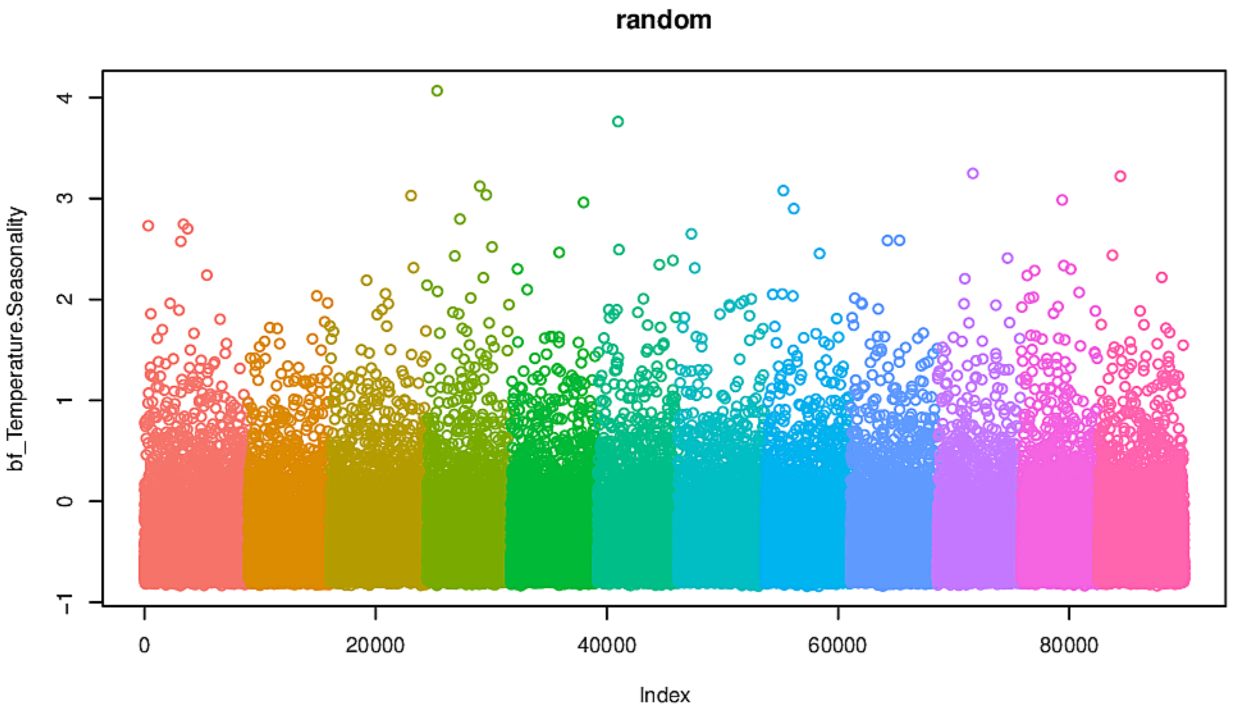
random
bf_Temperature.Seasonality
Index
0
20000
40000
60000
80000
-1
0
1
2
3
4

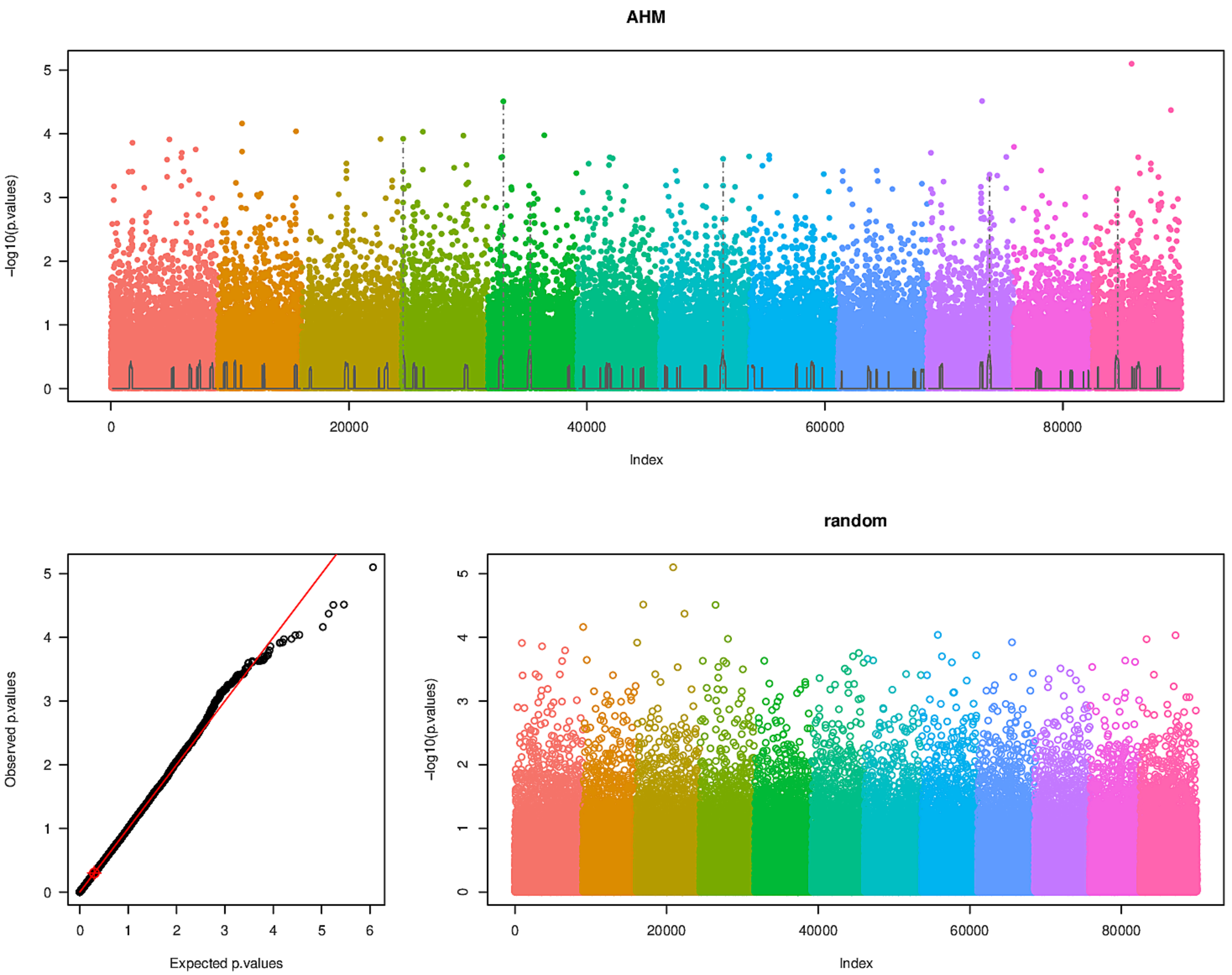
AHM
-log10(p.values)
0
1
2
3
4
5
0
20000
40000
60000
80000
Index
Observed p.values
Expected p.values
0
1
2
3
4
5
6
random
-log10(p.values)
Index

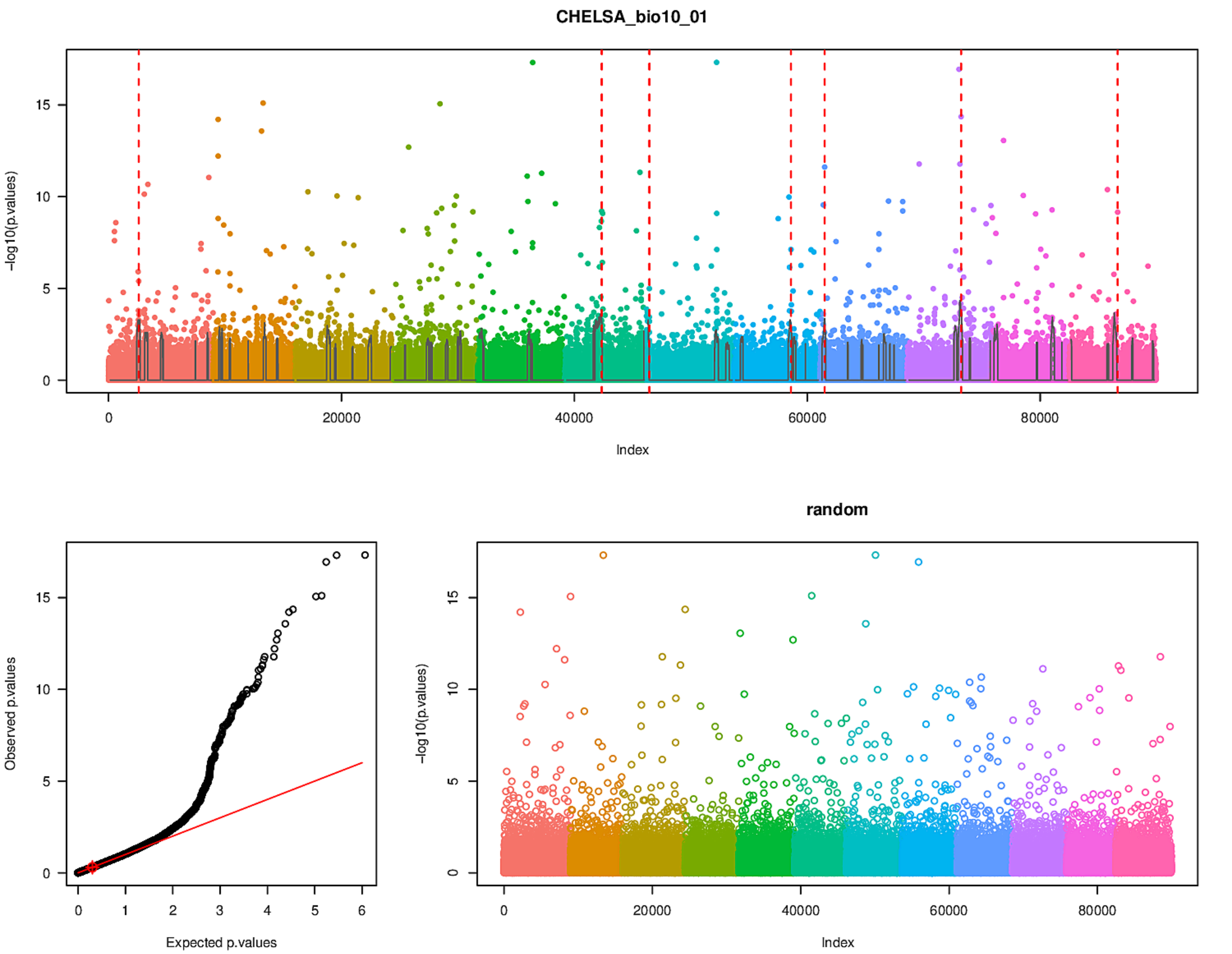
CHELSA_bio10_01
-log10(p.values)
Index
Observed p.values
Expected p.values
random
-log10(p.values)
Index

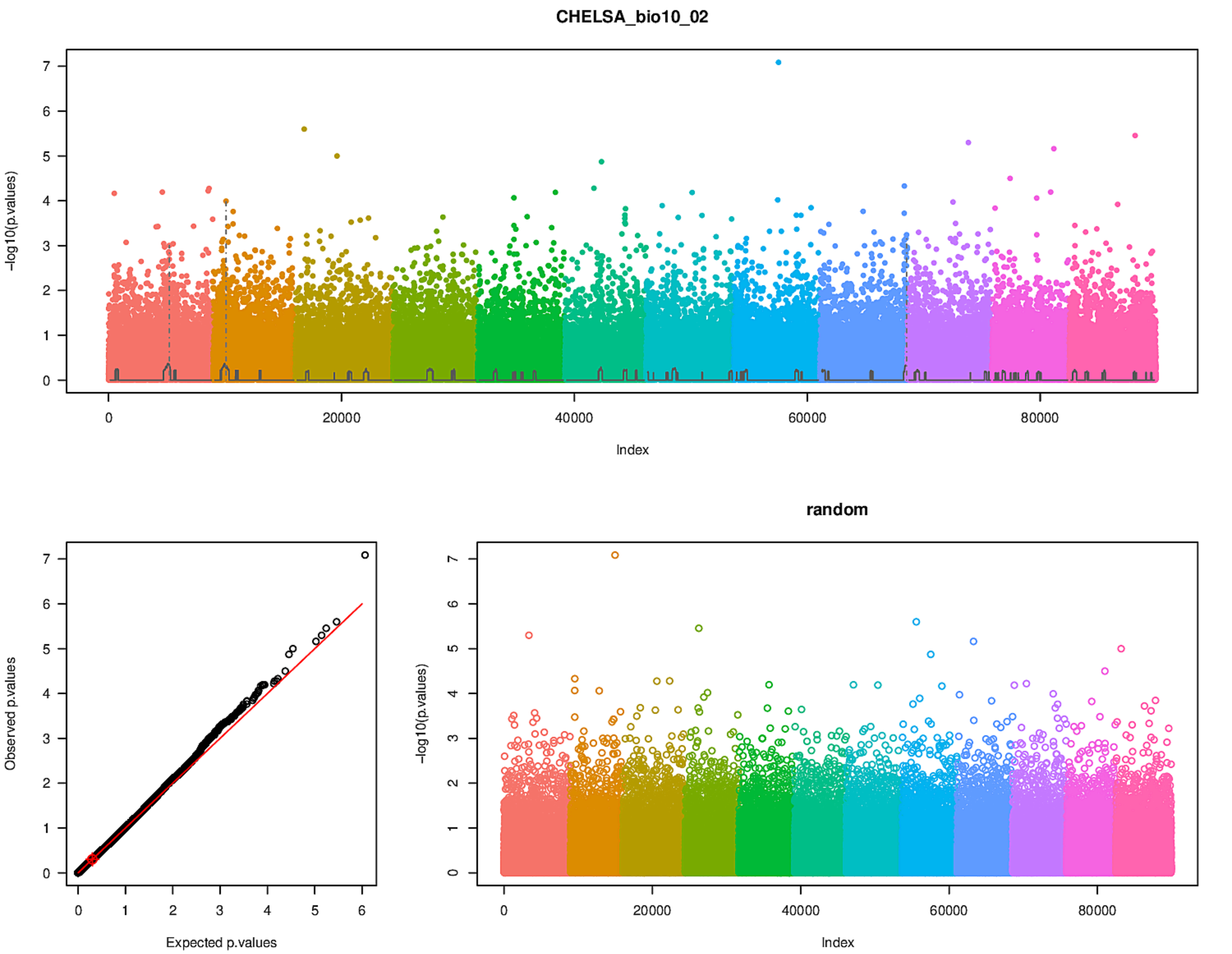
CHELSA_bio10_02
-log10(p.values)
Index
Observed p.values
Expected p.values
random
-log10(p.values)
Index

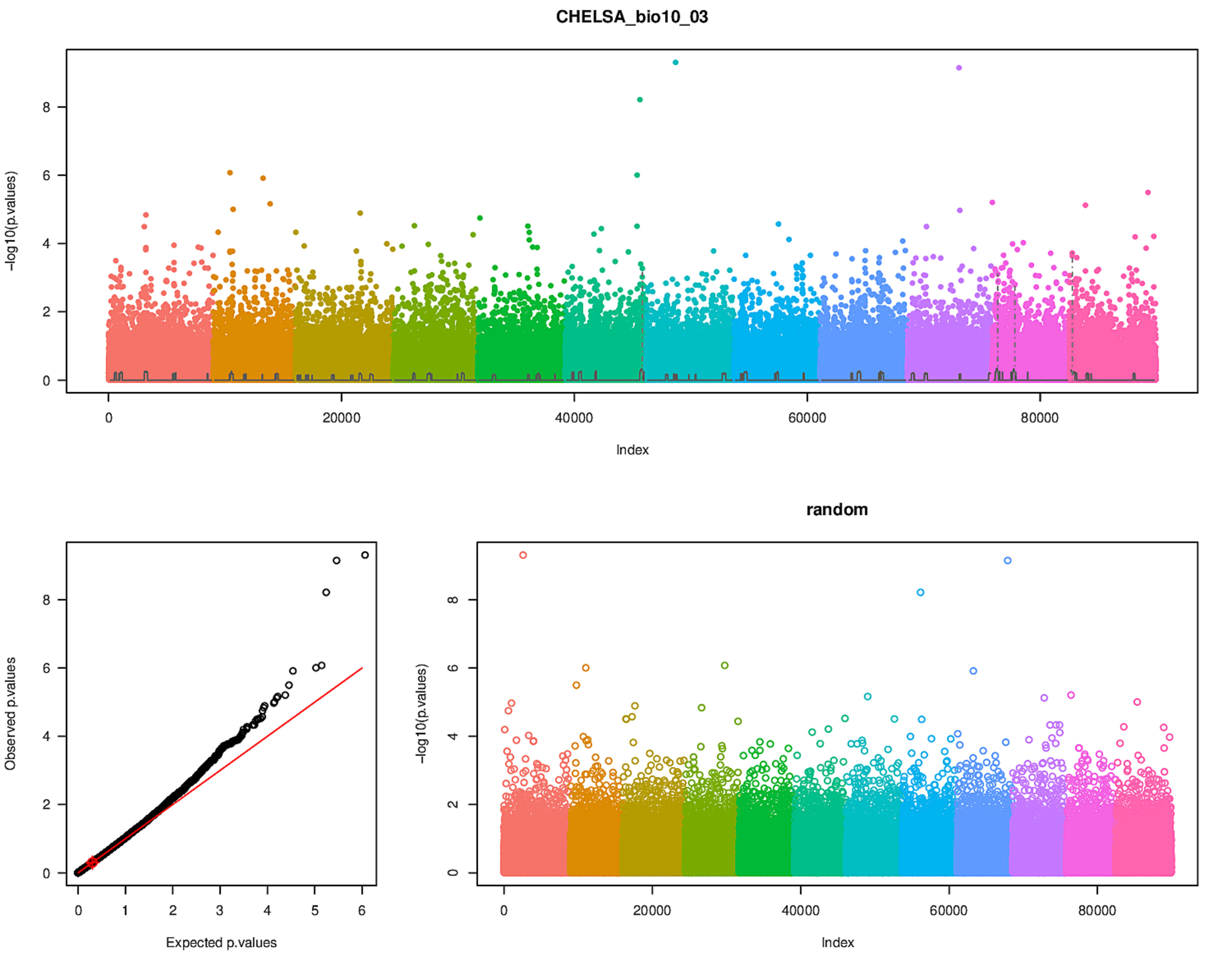
CHELSA_bio10_03
-log10(p.values)
Index
Observed p.values
Expected p.values
random
-log10(p.values)
Index

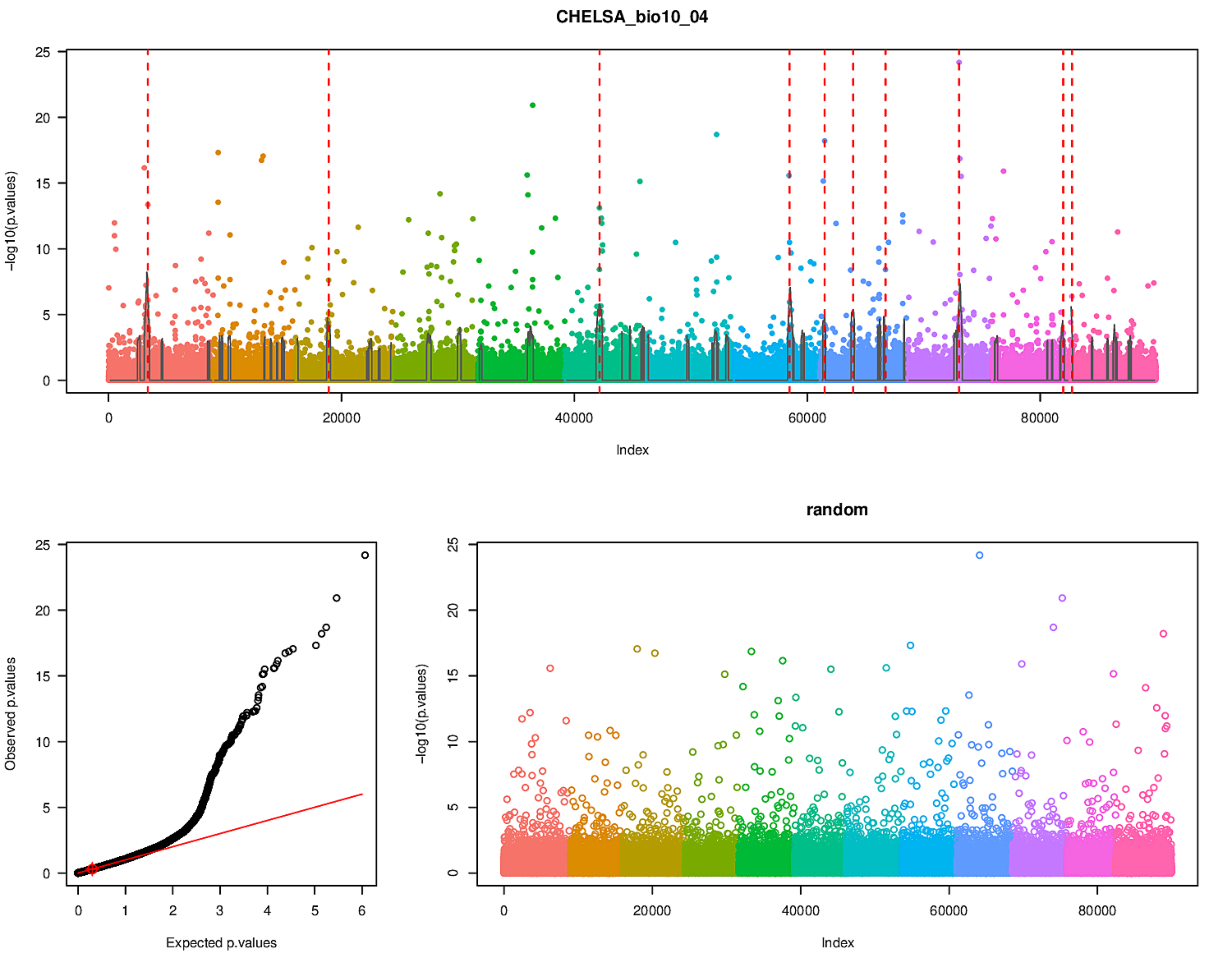

CHELSA_bio10_04
-log10(p.values)
Index
Observed p.values
Expected p.values
random
-log10(p.values)
Index

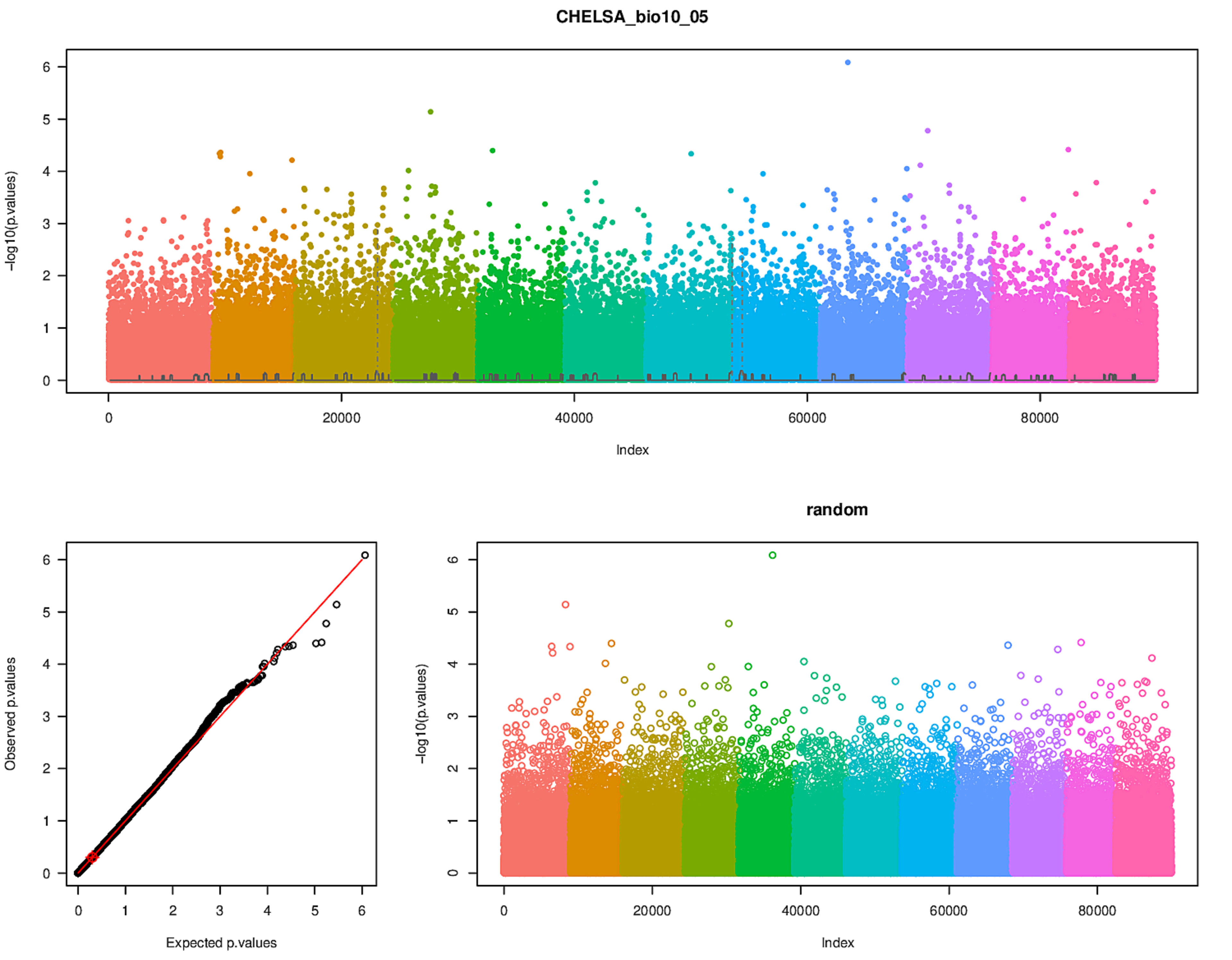
CHELSA_bio10_05
-log10(p.values)
Index
Observed p.values
Expected p.values
random
-log10(p.values)
Index

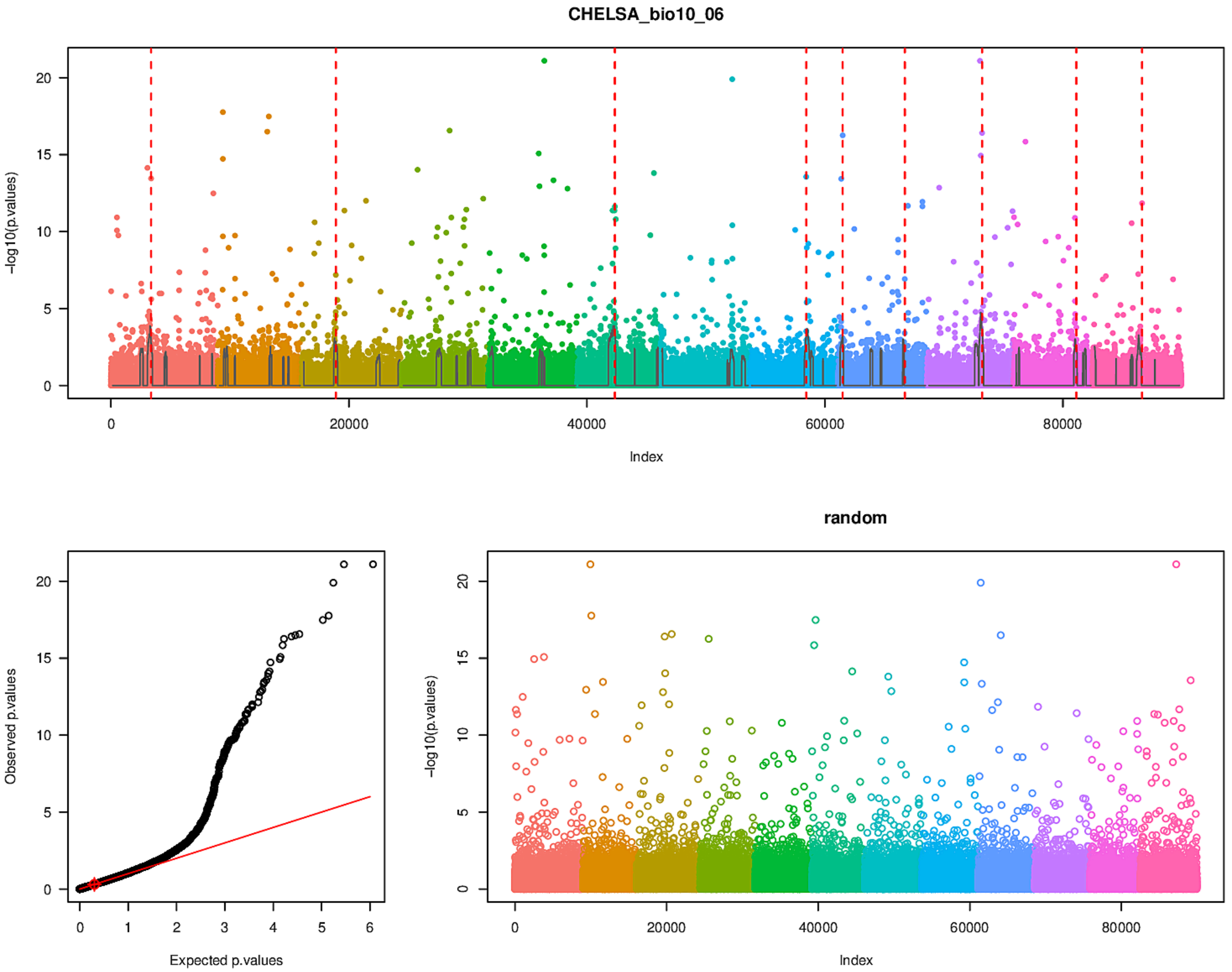
CHELSA_bio10_06
-log10(p.values)
Index
0
5
10
15
20
0
20000
40000
60000
80000
random
Observed p.values
Expected p.values
0
1
2
3
4
5
6

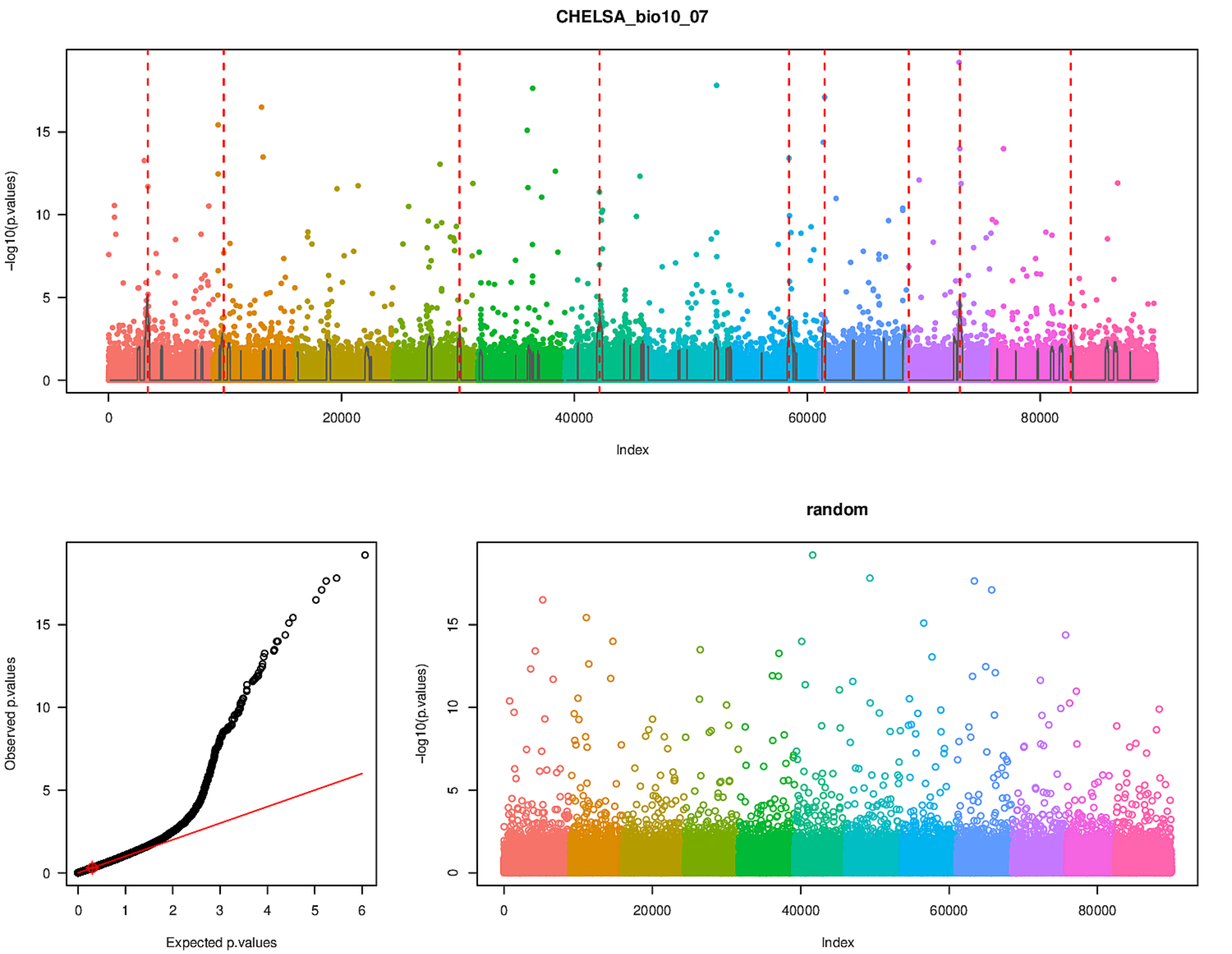

CHELSA_bio10_07
-log10(p.values)
Index
0
5
10
15
20000
40000
60000
80000
Observed p.values
Expected p.values
0
1
2
3
4
5
6
random

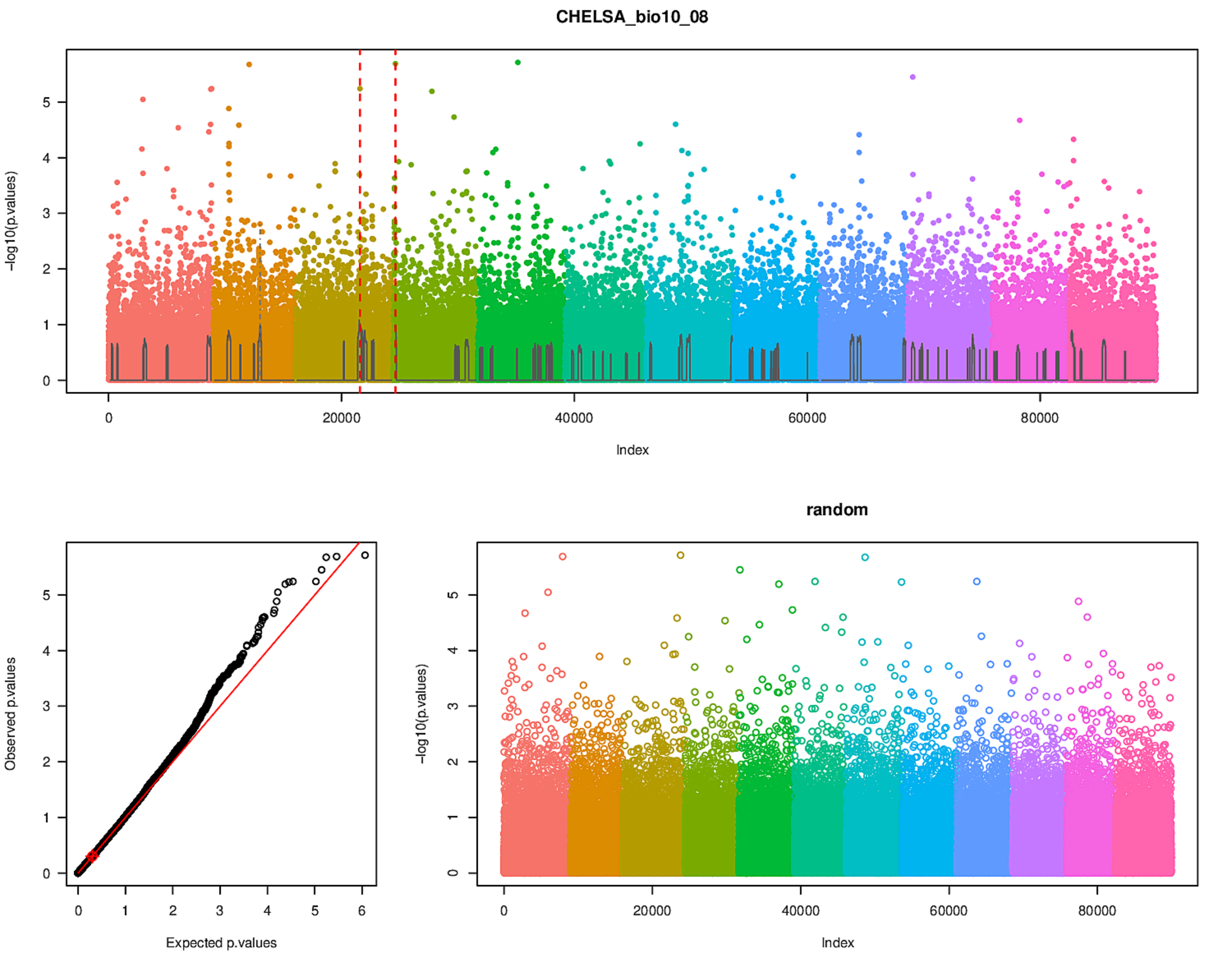
CHELSA_bio10_08
-log10(p.values)
Index
Observed p.values
Expected p.values
random
-log10(p.values)
Index

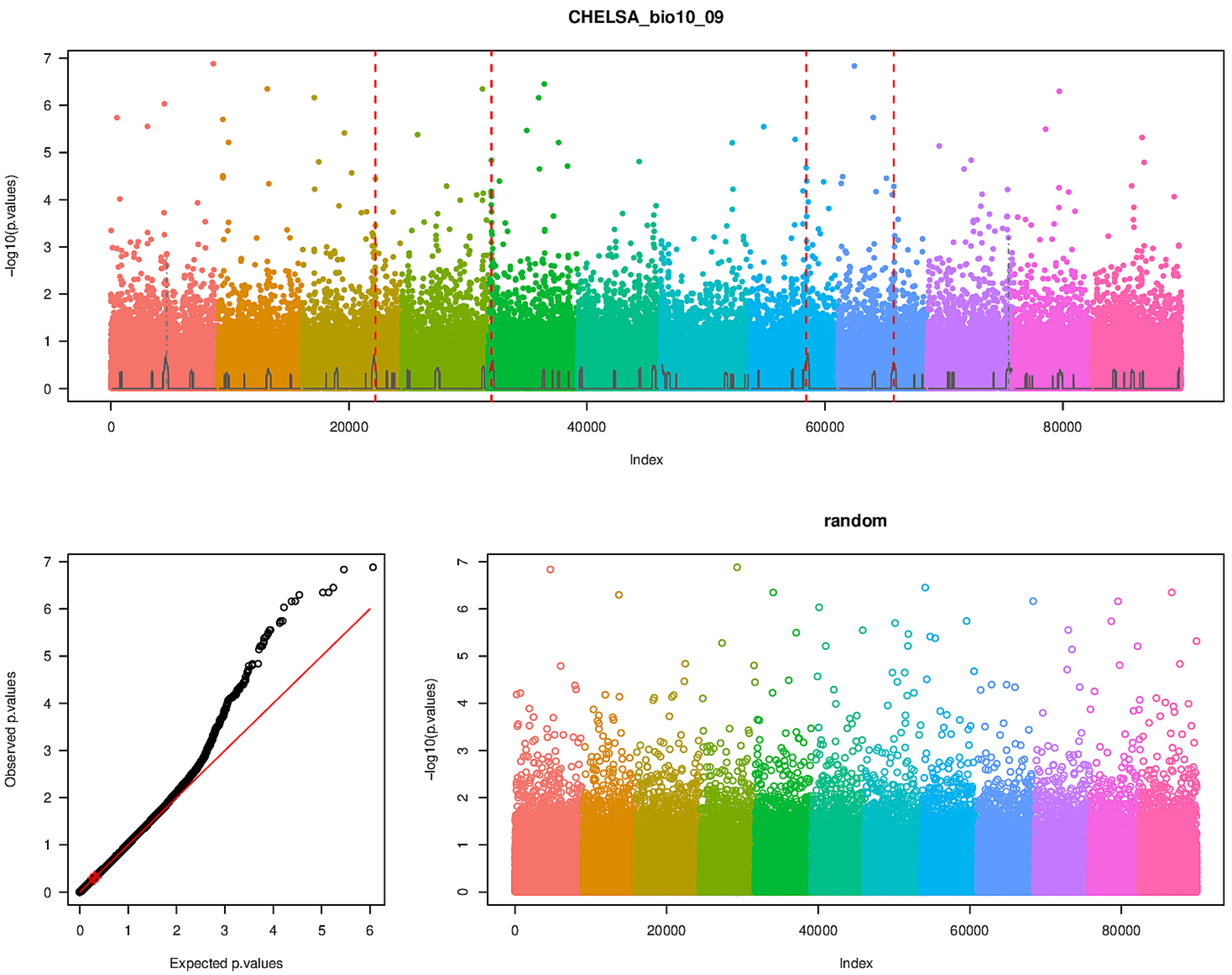
CHELSA_bio10_09
-log10(p.values)
Index
0
20000
40000
60000
80000
Observed p.values
Expected p.values
random
-log10(p.values)
Index

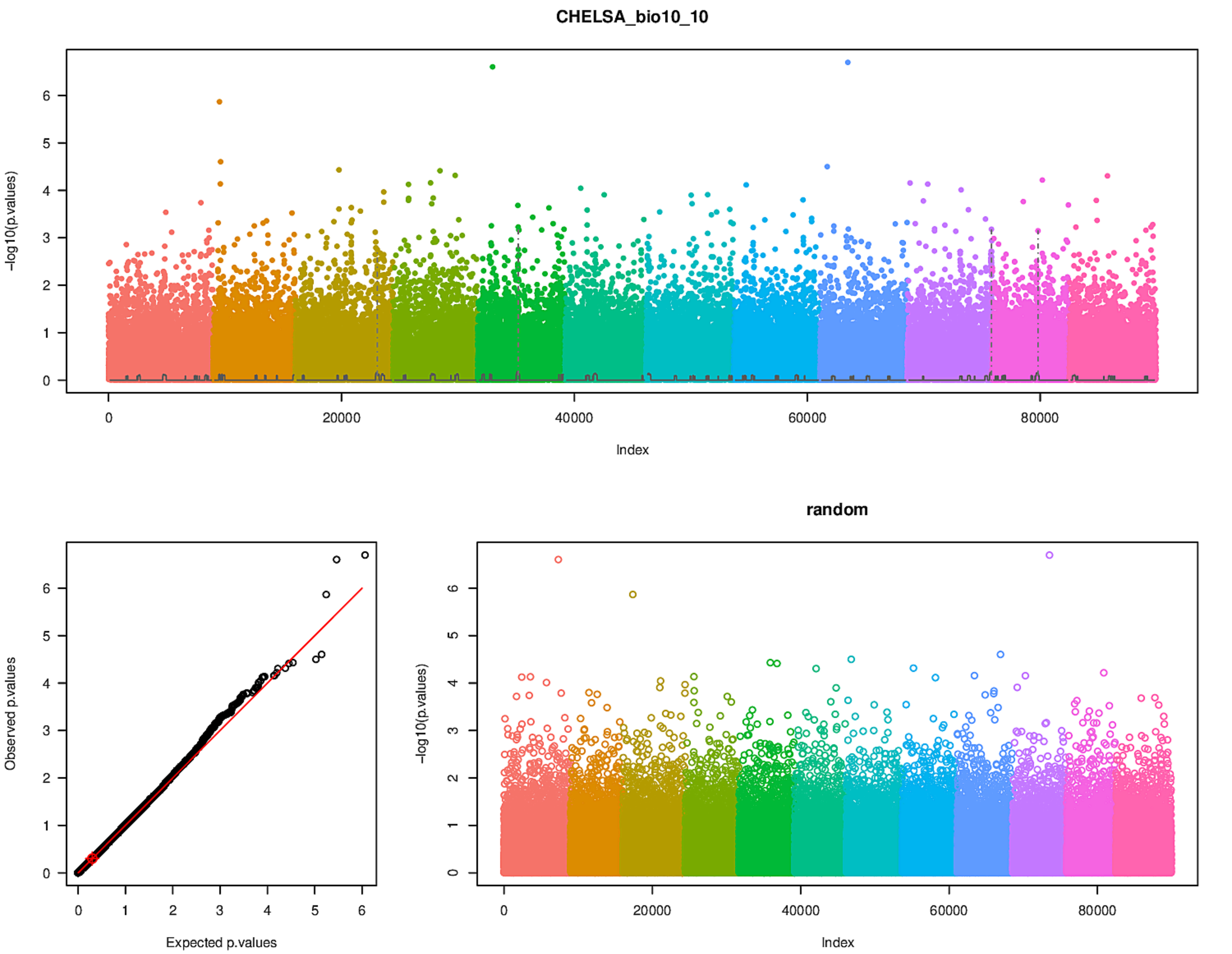
CHELSA_bio10_10
−log10(p.values)
Index
Observed p.values
Expected p.values
random
−log10(p.values)
Index

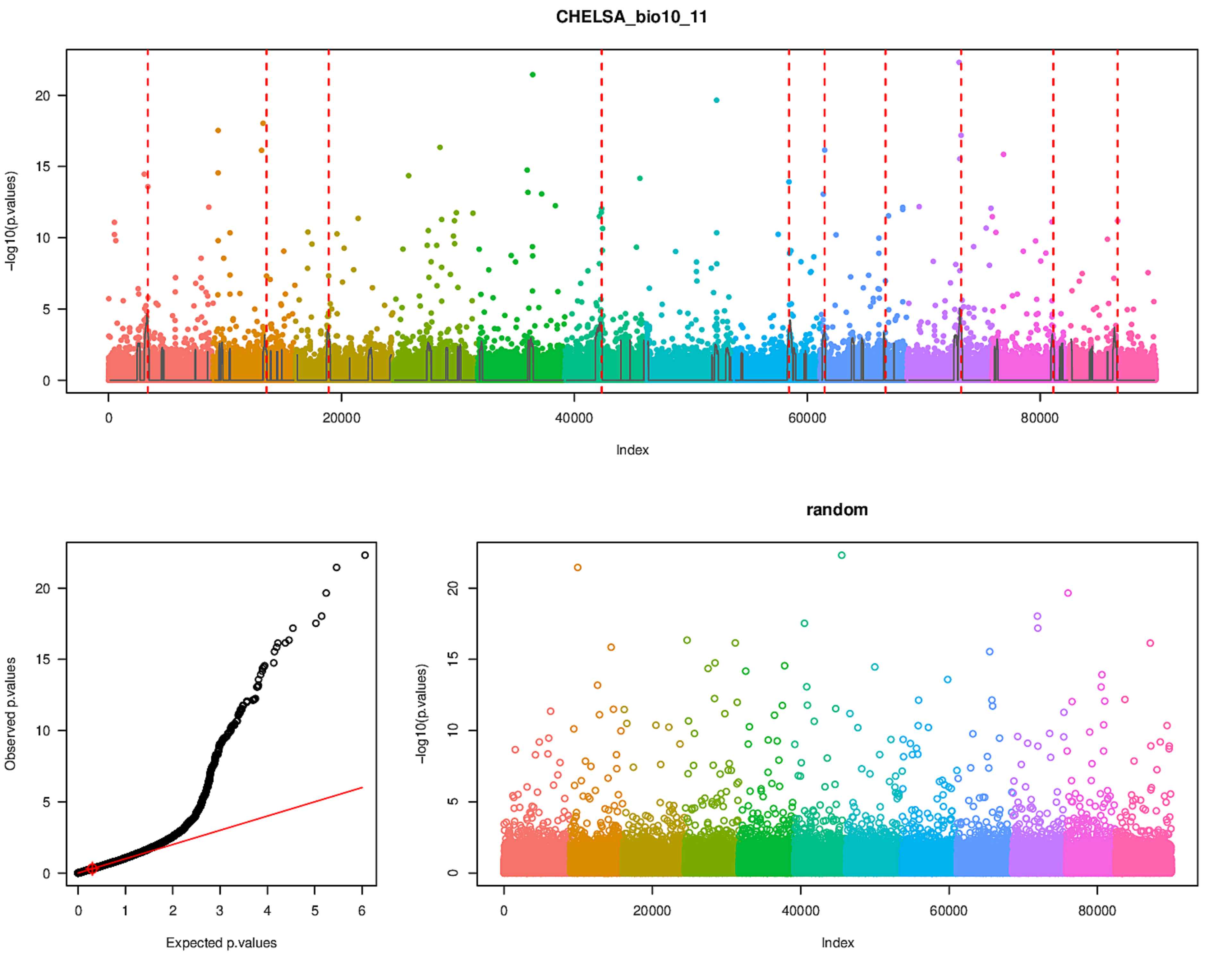
CHELSA_bio10_11
-log10(p.values)
Index
Observed p.values
Expected p.values
random
-log10(p.values)
Index

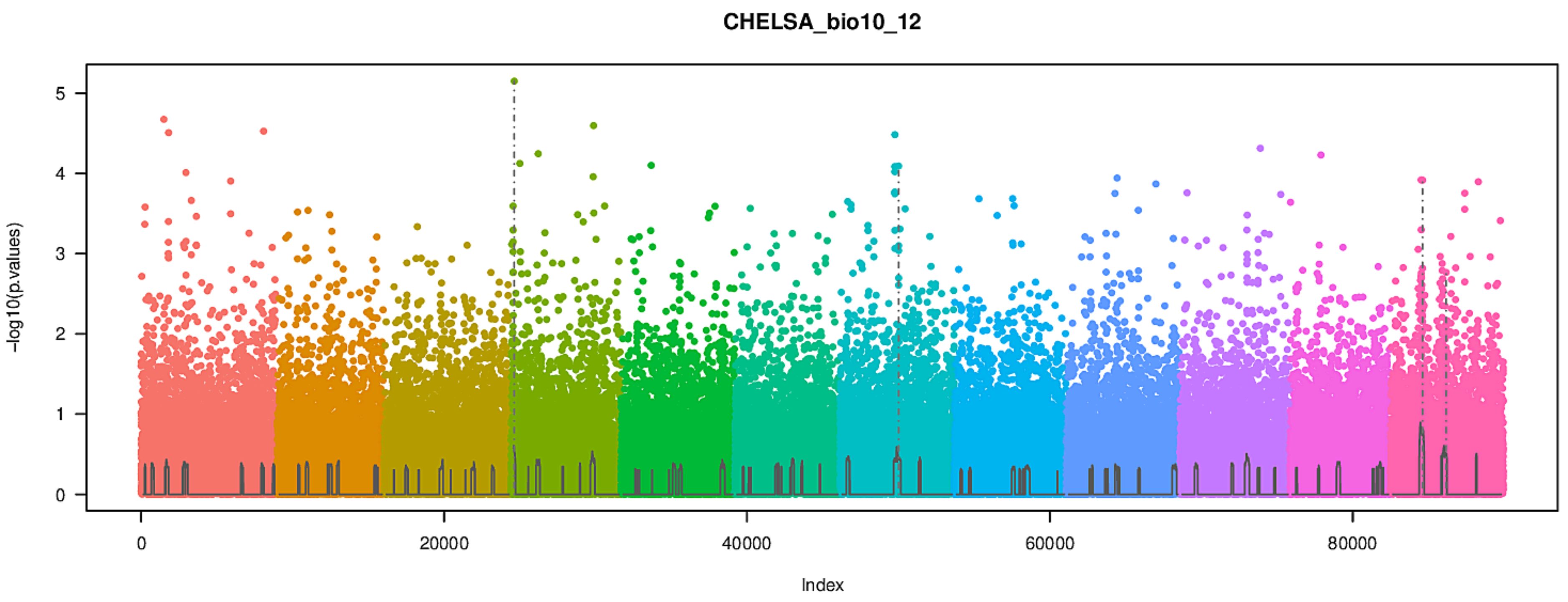
CHELSA_bio10_12
-log10(p.values)
Index
0
20000
40000
60000
80000

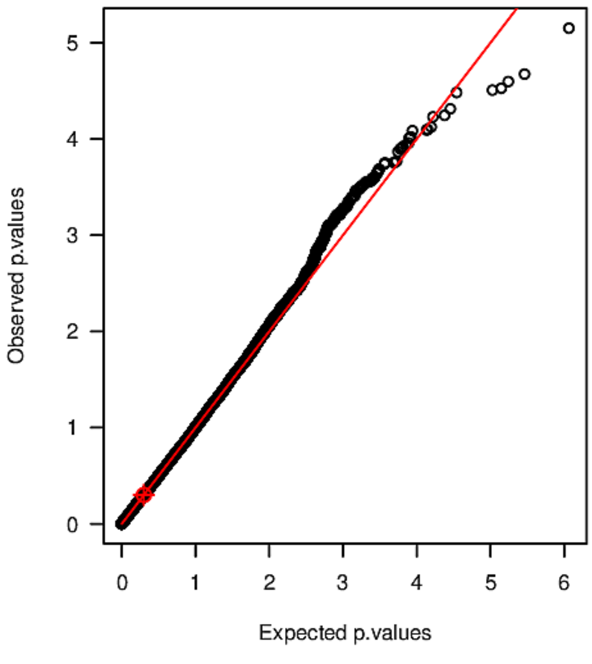
Observed p.values
Expected p.values

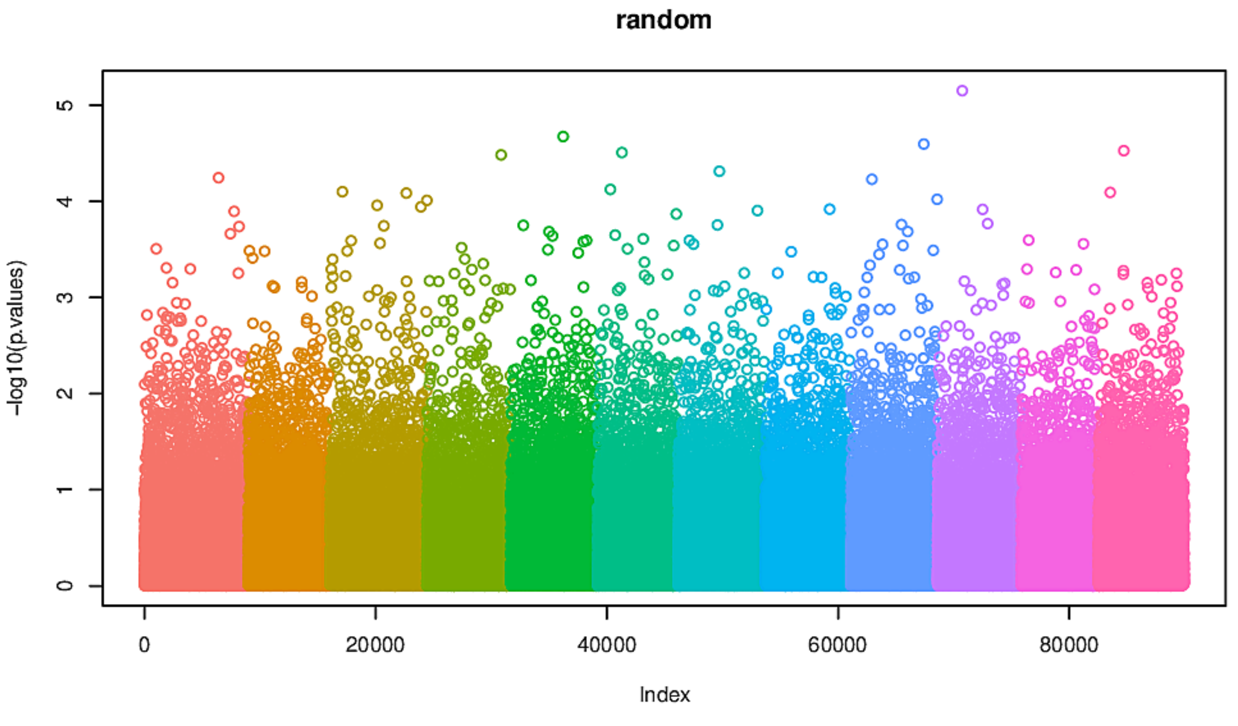
random
-log10(p.values)
Index
0
20000
40000
60000
80000

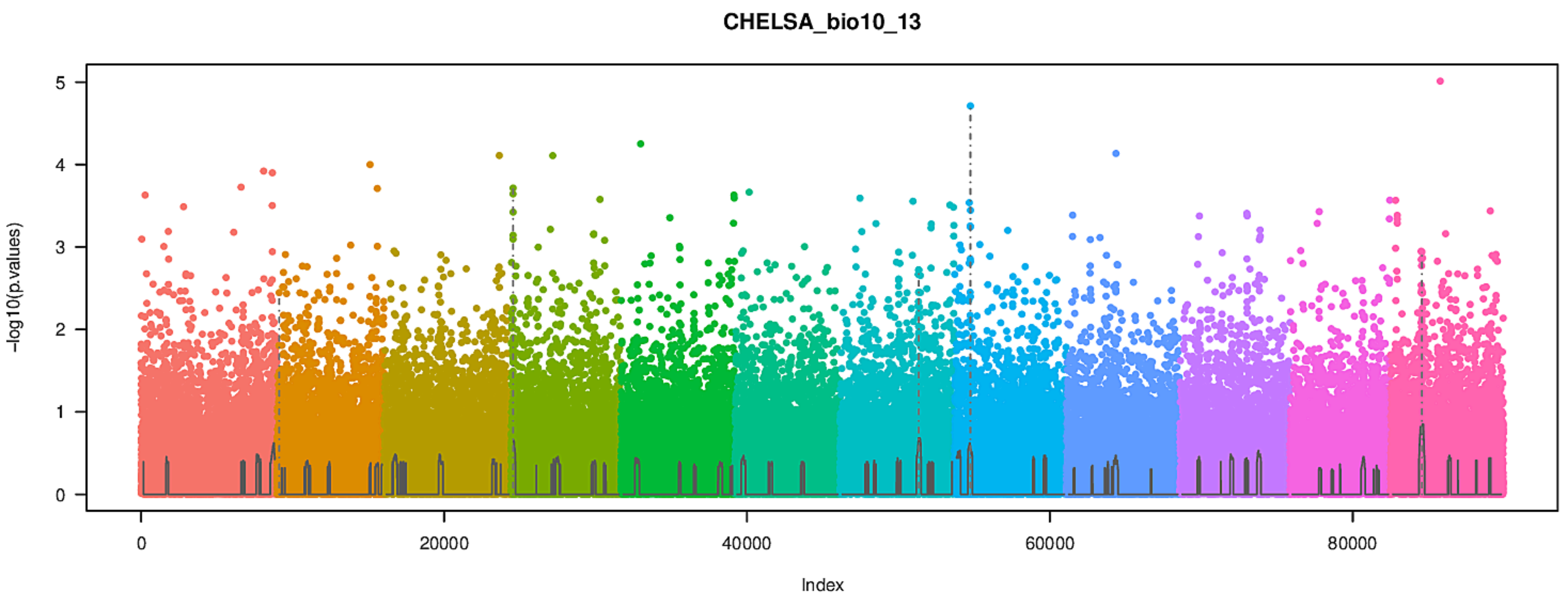
CHELSA_bio10_13
-log10(p.values)
Index
0
20000
40000
60000
80000
0
1
2
3
4
5

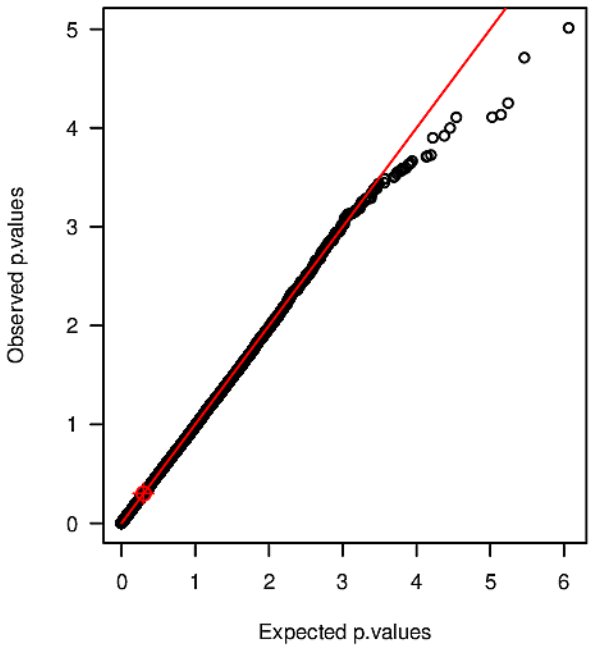
Observed p.values
Expected p.values
0
1
2
3
4
5
6

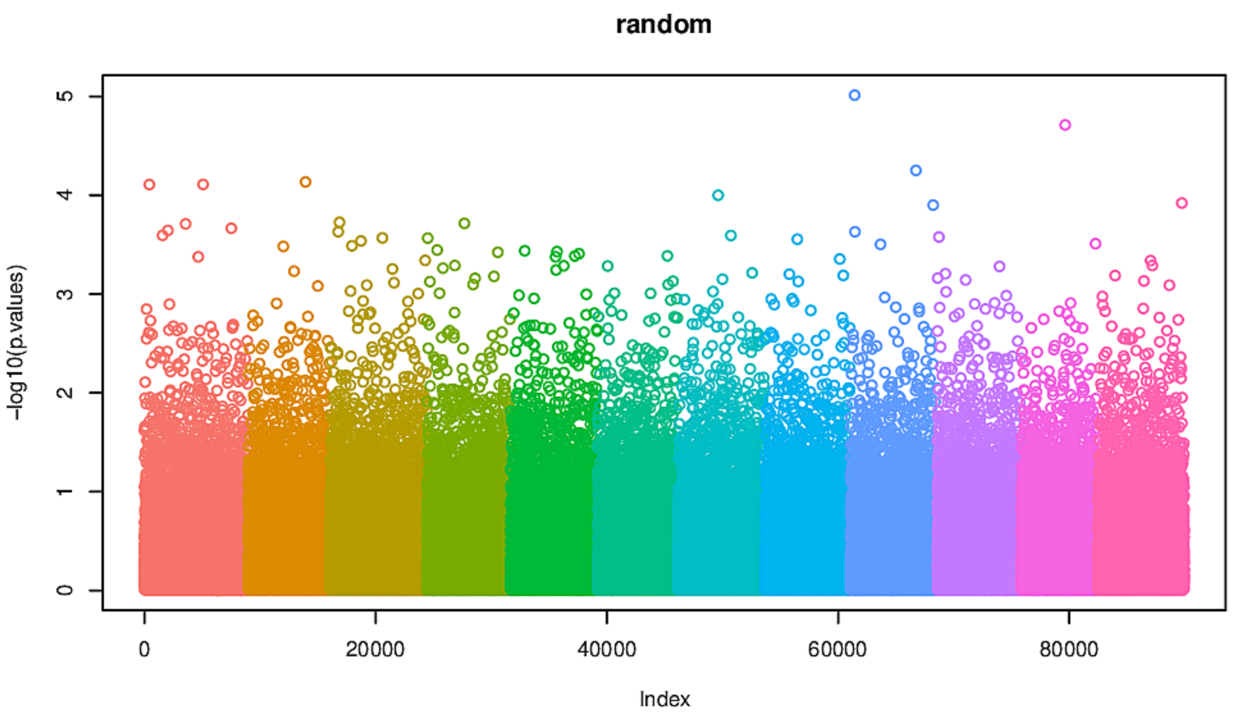
random
-log10(p.values)
Index
0
20000
40000
60000
80000
0
1
2
3
4
5

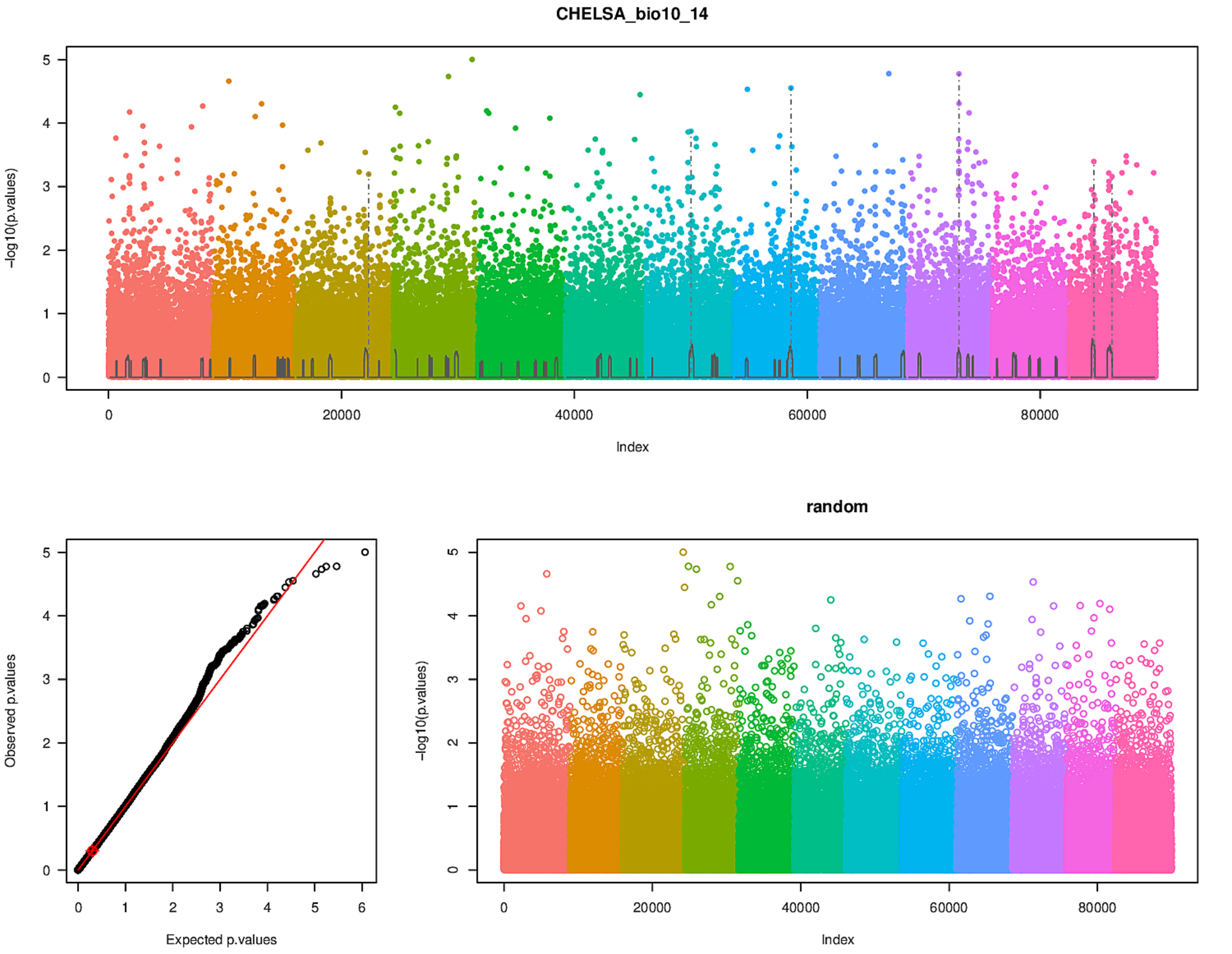
CHELSA_bio10_14
-log10(p.values)
0
1
2
3
4
5
0
20000
40000
60000
80000
Index
Observed p.values
Expected p.values
0
1
2
3
4
5
6
random
-log10(p.values)
Index

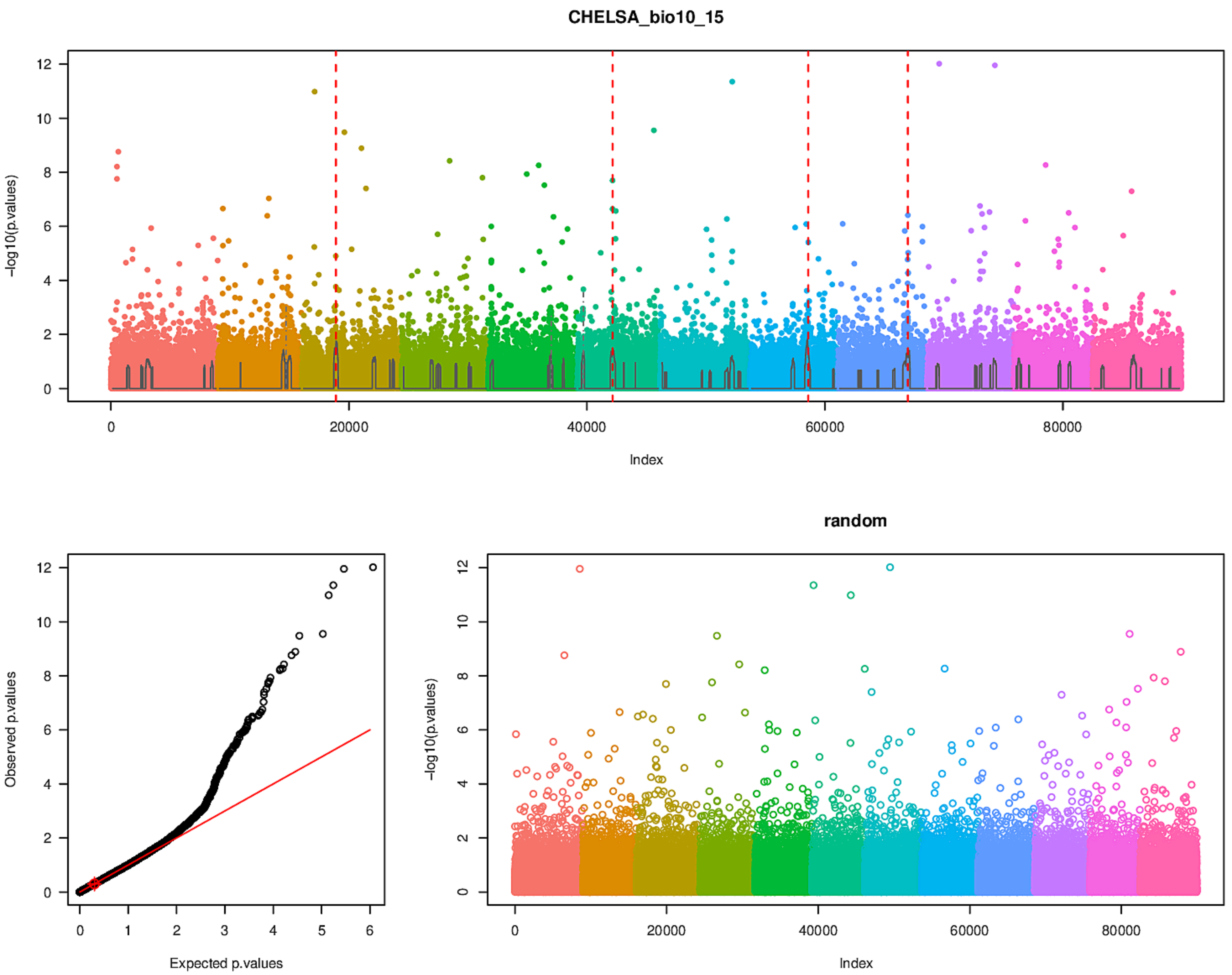
CHELSA_bio10_15
-log10(p.values)
0
2
4
6
8
10
12
0
20000
40000
60000
80000
Index
random
Observed p.values
Expected p.values
0
1
2
3
4
5
6

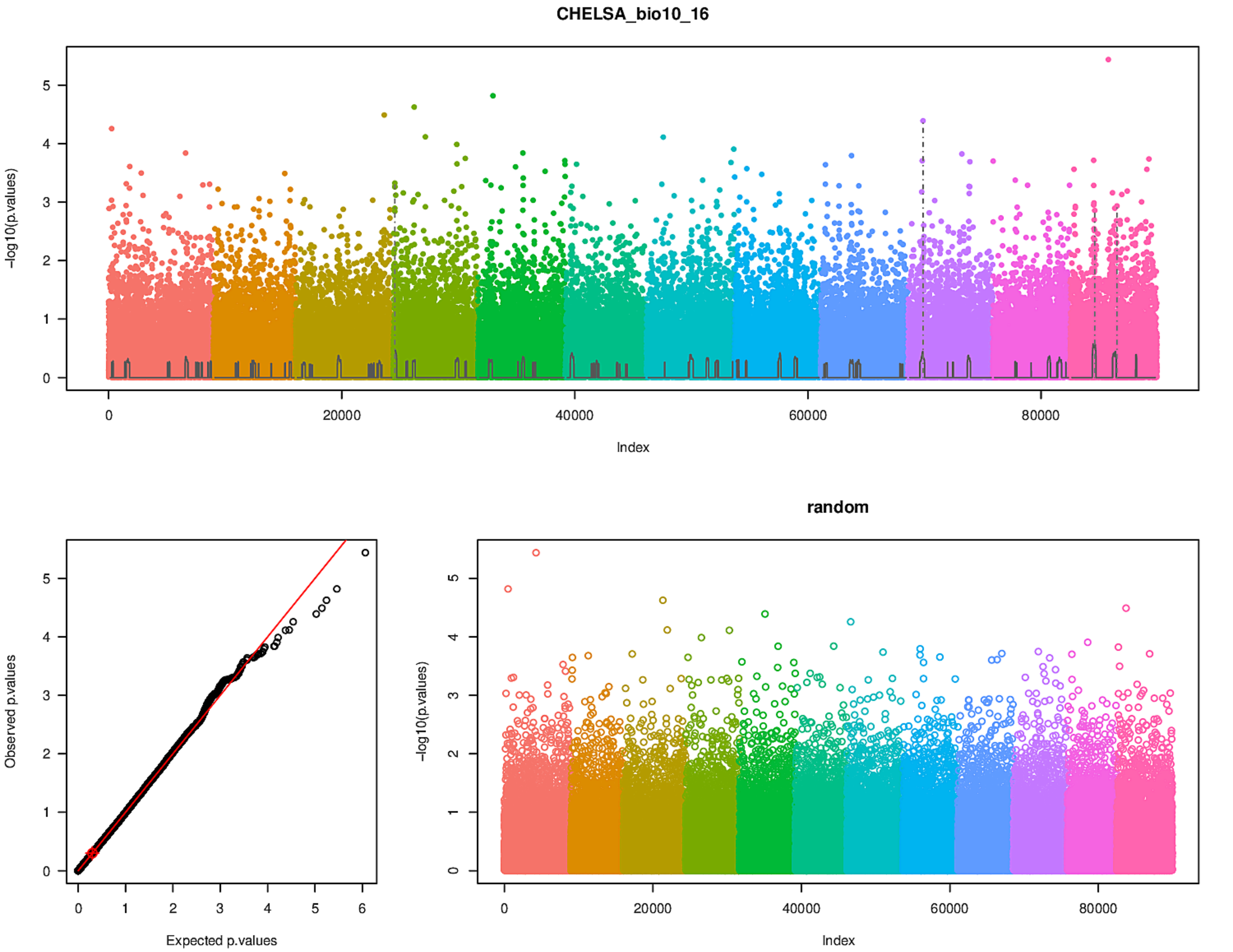
CHELSA_bio10_16
−log10(p.values)
Index
0
20000
40000
60000
80000
0
1
2
3
4
5
Observed p.values
Expected p.values
0
1
2
3
4
5
6
random
−log10(p.values)
Index

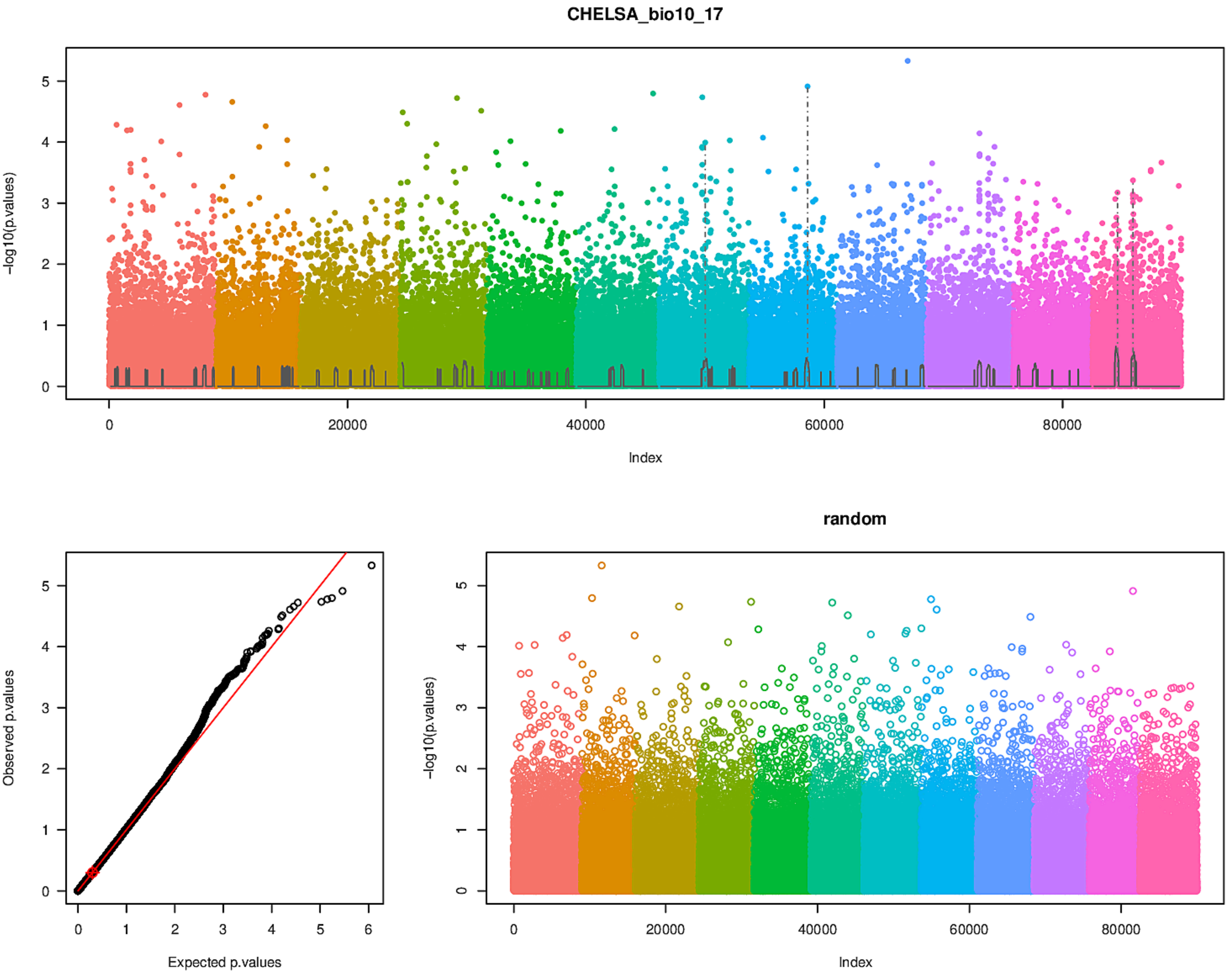
CHELSA_bio10_17
-log10(p.values)
Index
Observed p.values
Expected p.values
random
-log10(p.values)
Index

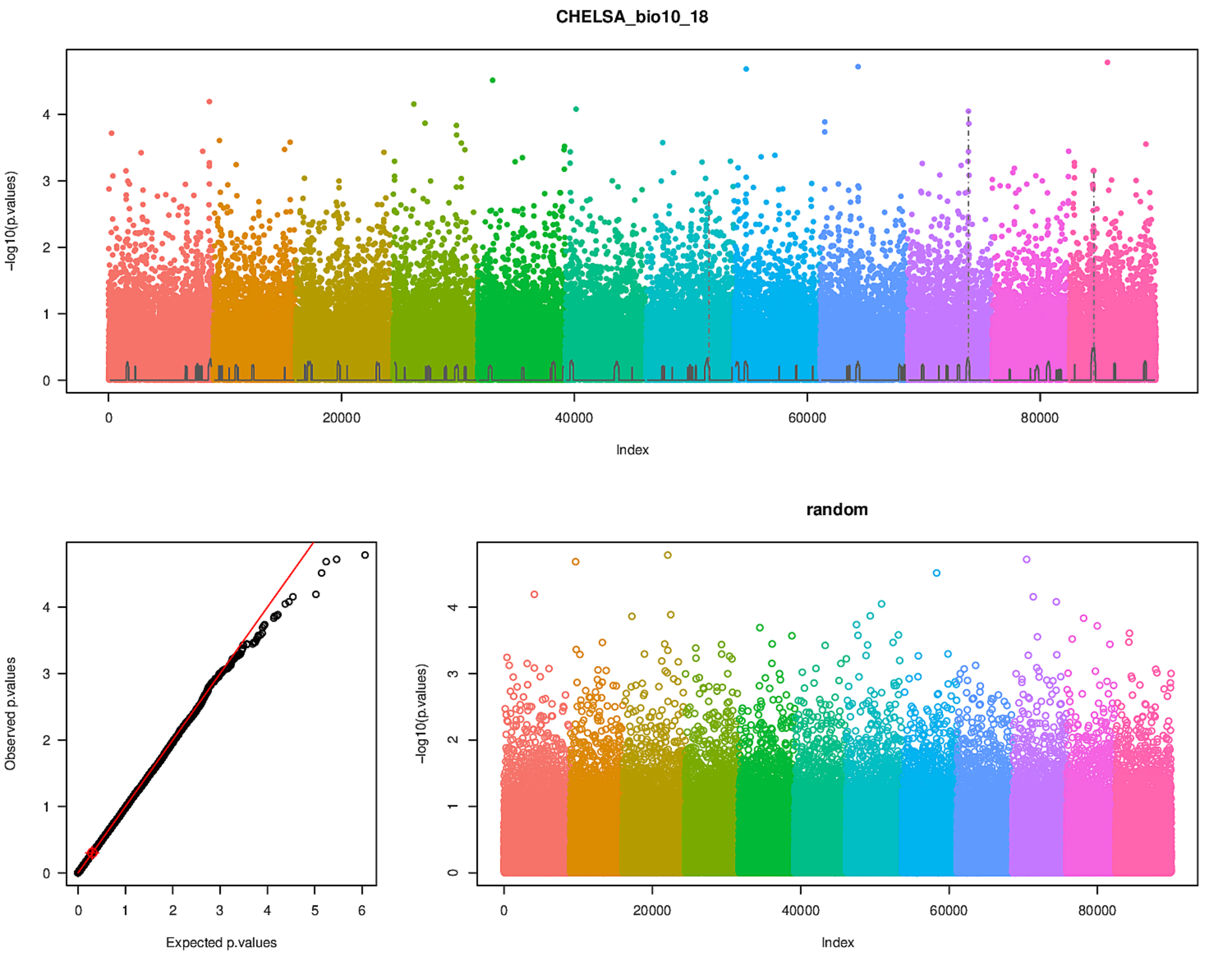
CHELSA_bio10_18
-log10(p.values)
Index
Observed p.values
Expected p.values
random
-log10(p.values)
Index

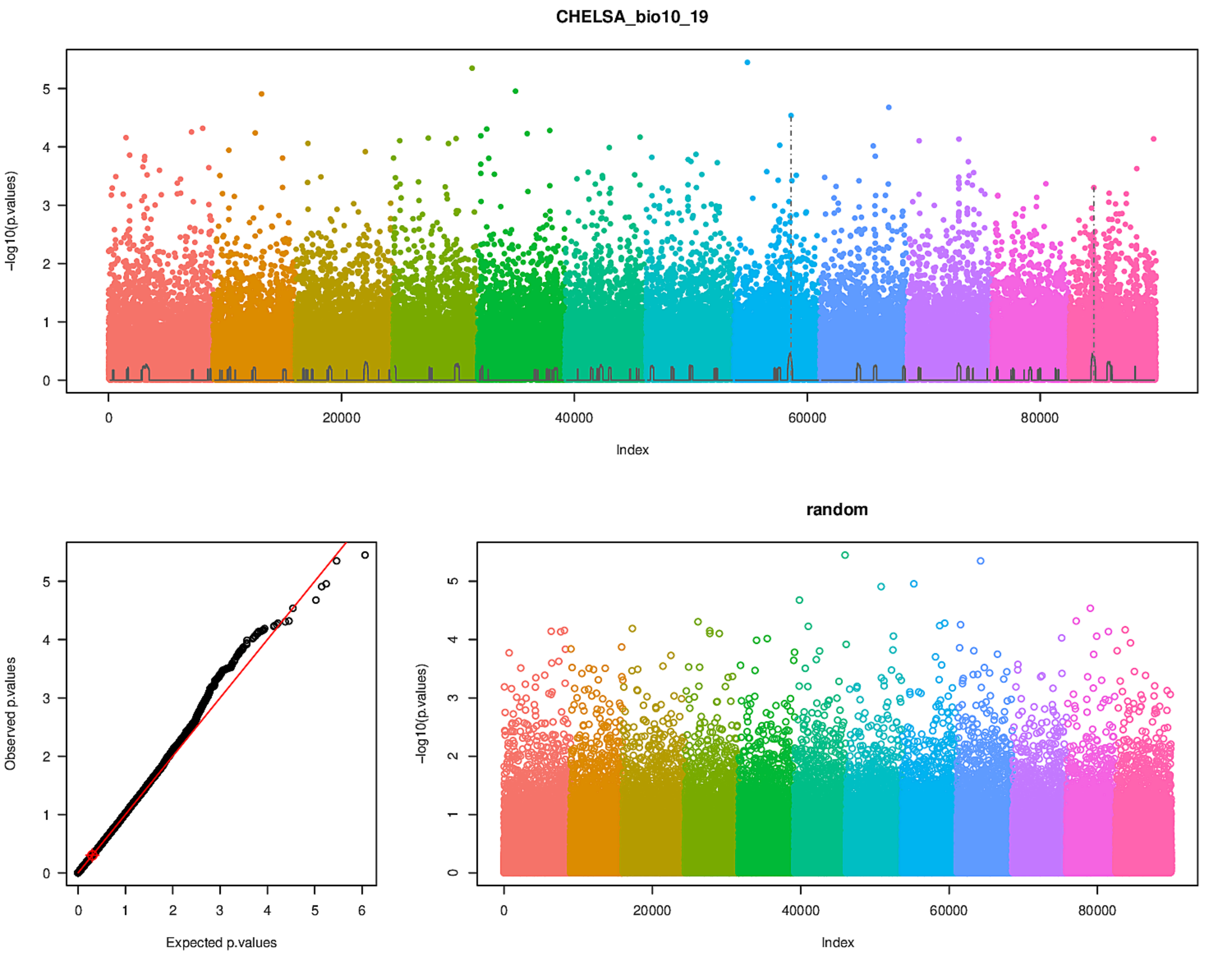

CHELSA_bio10_19
-log10(p.values)
Index
Observed p.values
Expected p.values
random
-log10(p.values)
Index

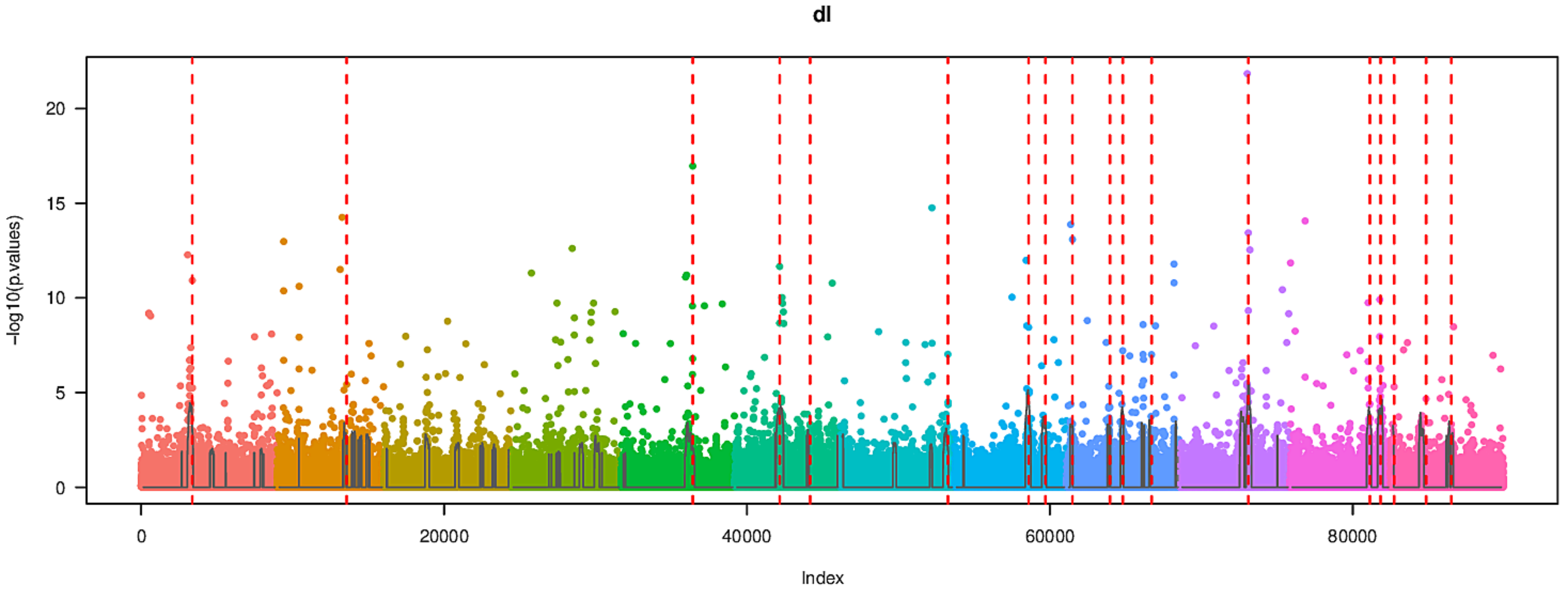
dl
-log10(p.values)
Index

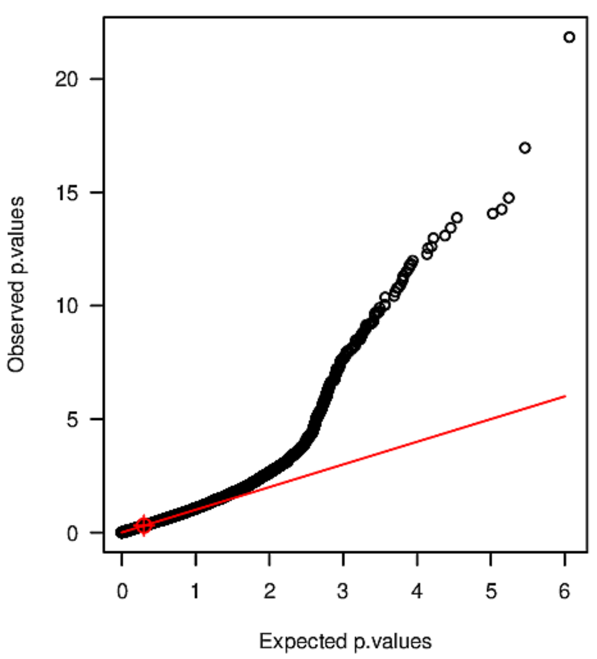
Observed p.values
Expected p.values

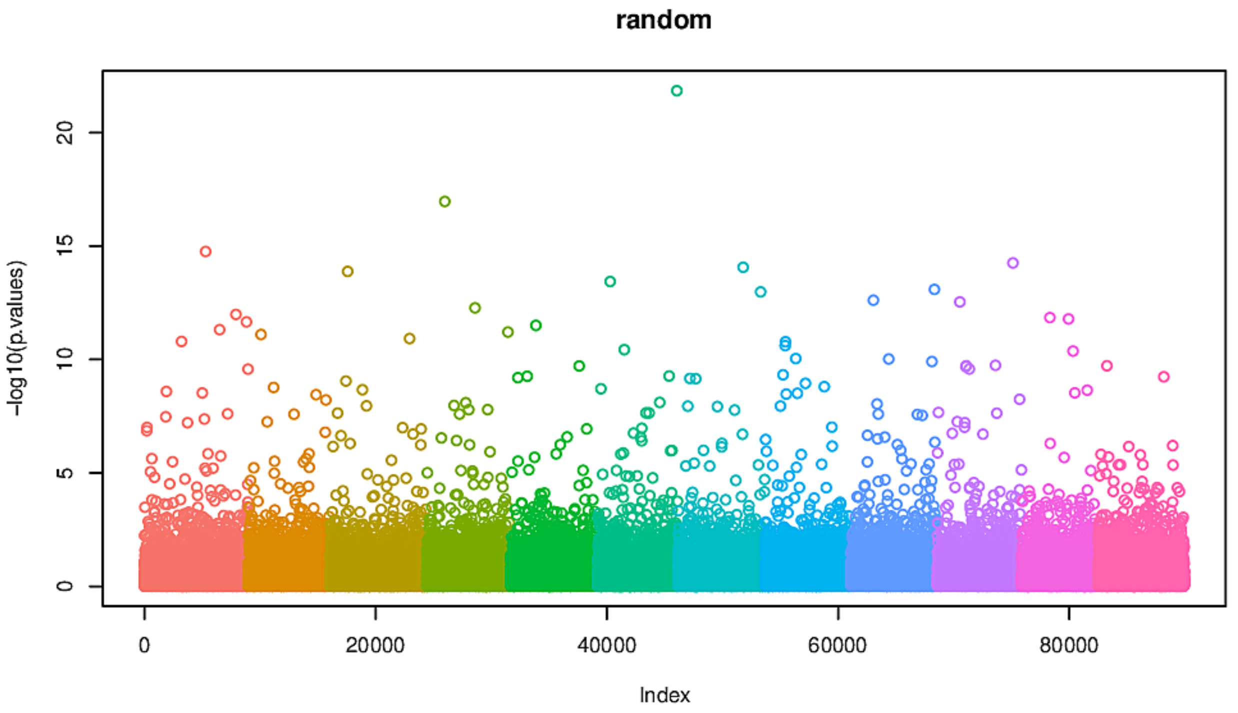
random
-log10(p.values)
Index

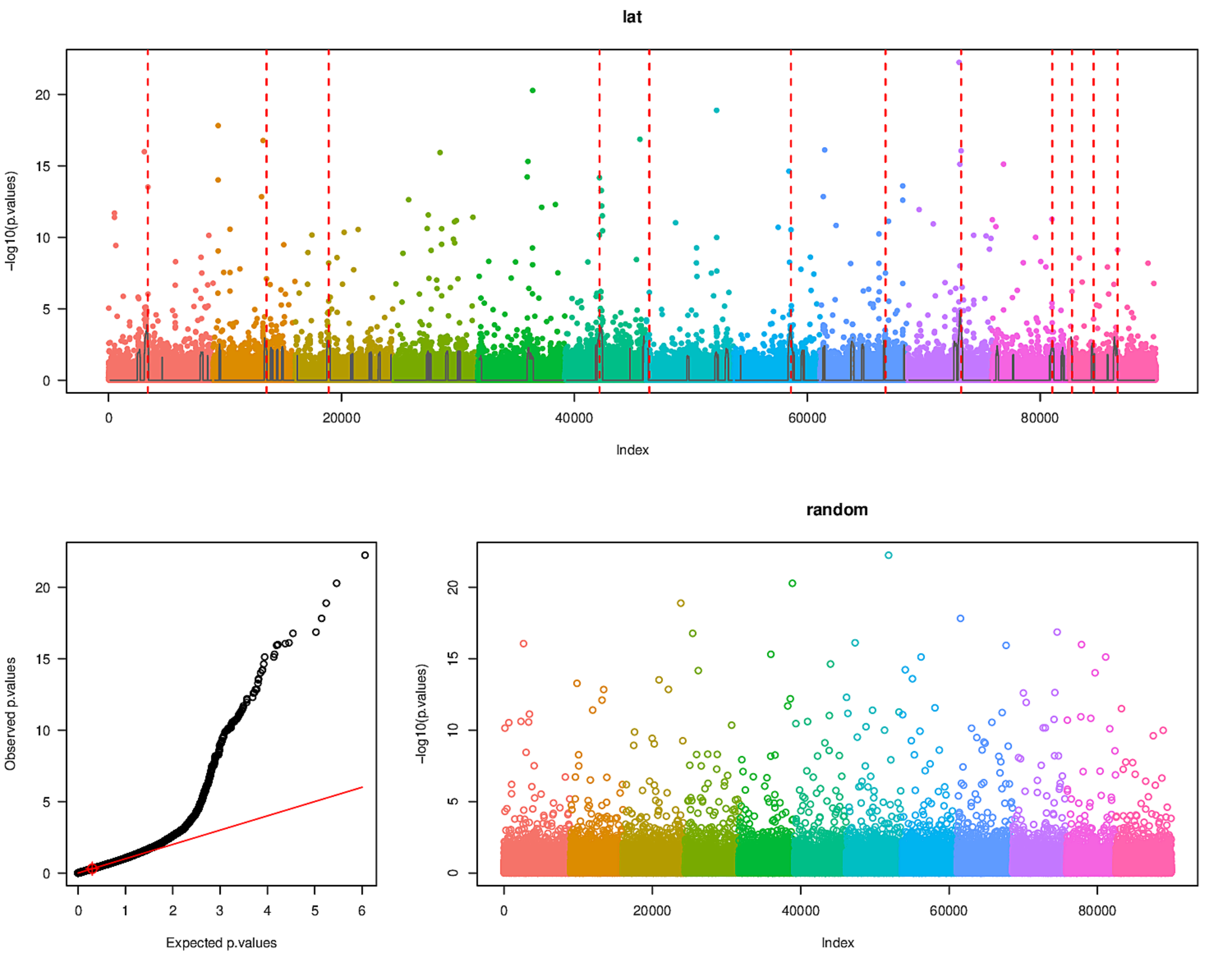
lat
-log10(p.values)
Index
0
20000
40000
60000
80000
0
5
10
15
20
Observed p.values
Expected p.values
0
1
2
3
4
5
6
random
-log10(p.values)
Index

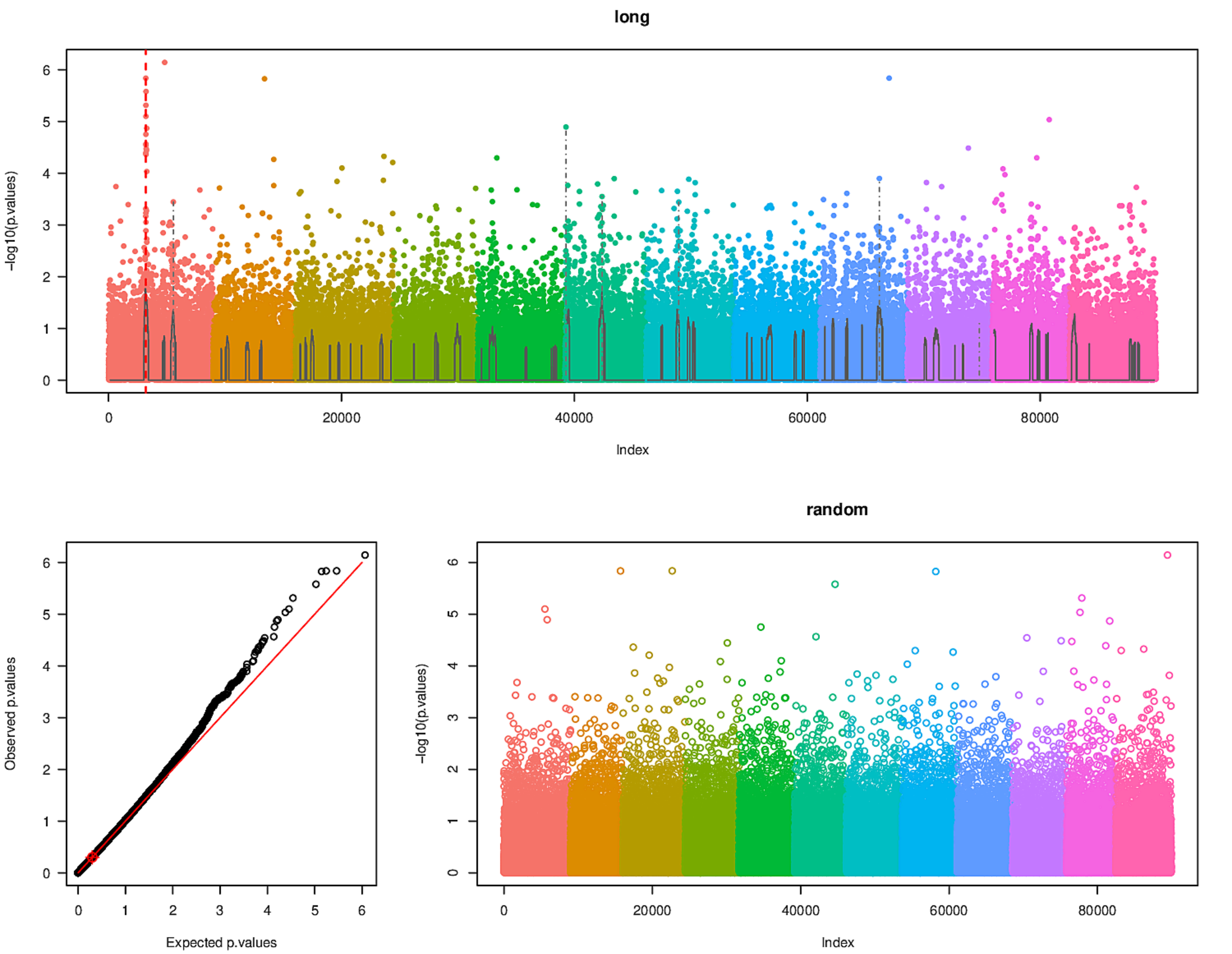
long
-log10(p.values)
Index
Observed p.values
Expected p.values
random
-log10(p.values)
Index

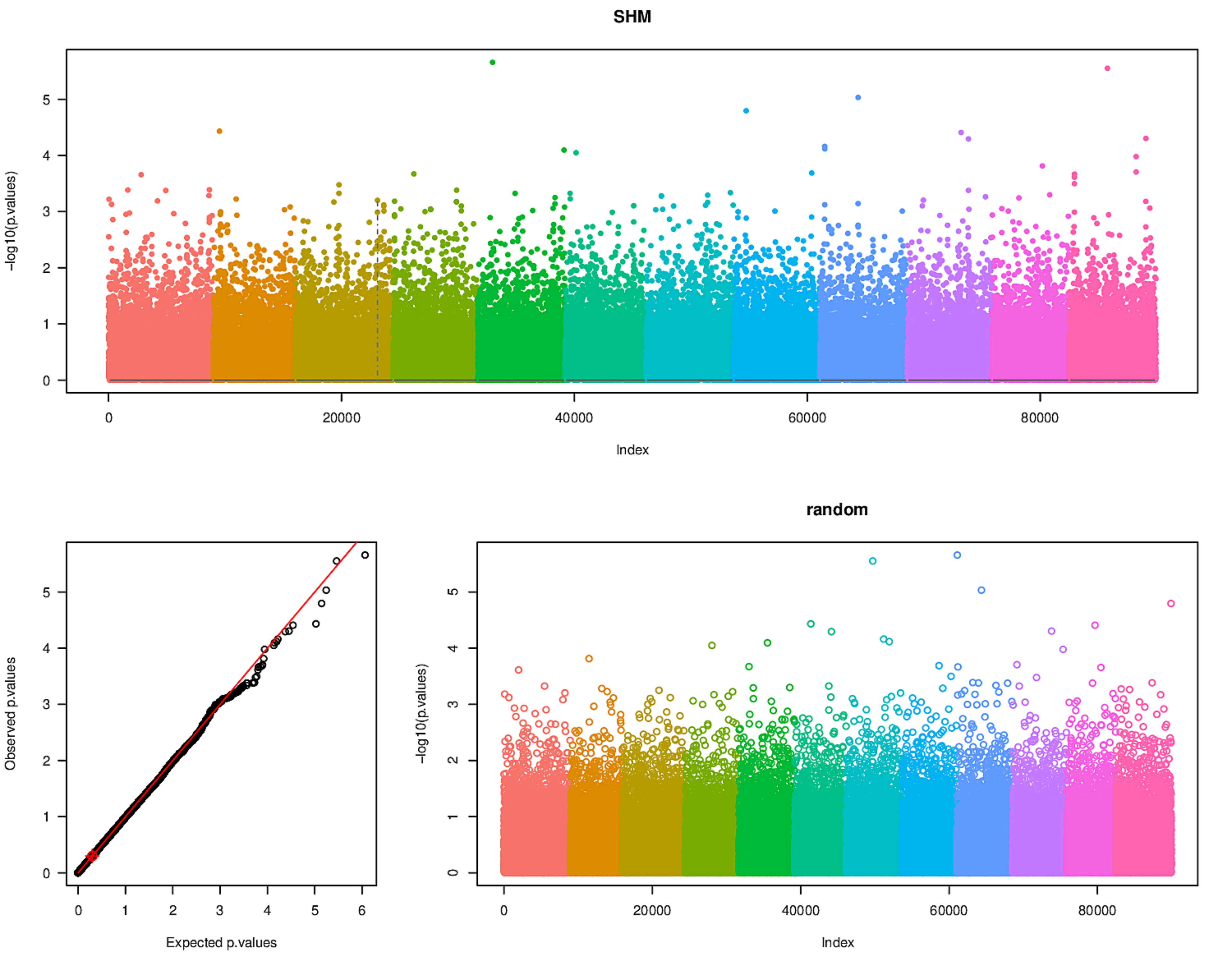

SHM
-log10(p.values)
Index
0
20000
40000
60000
80000
Observed p.values
Expected p.values
random

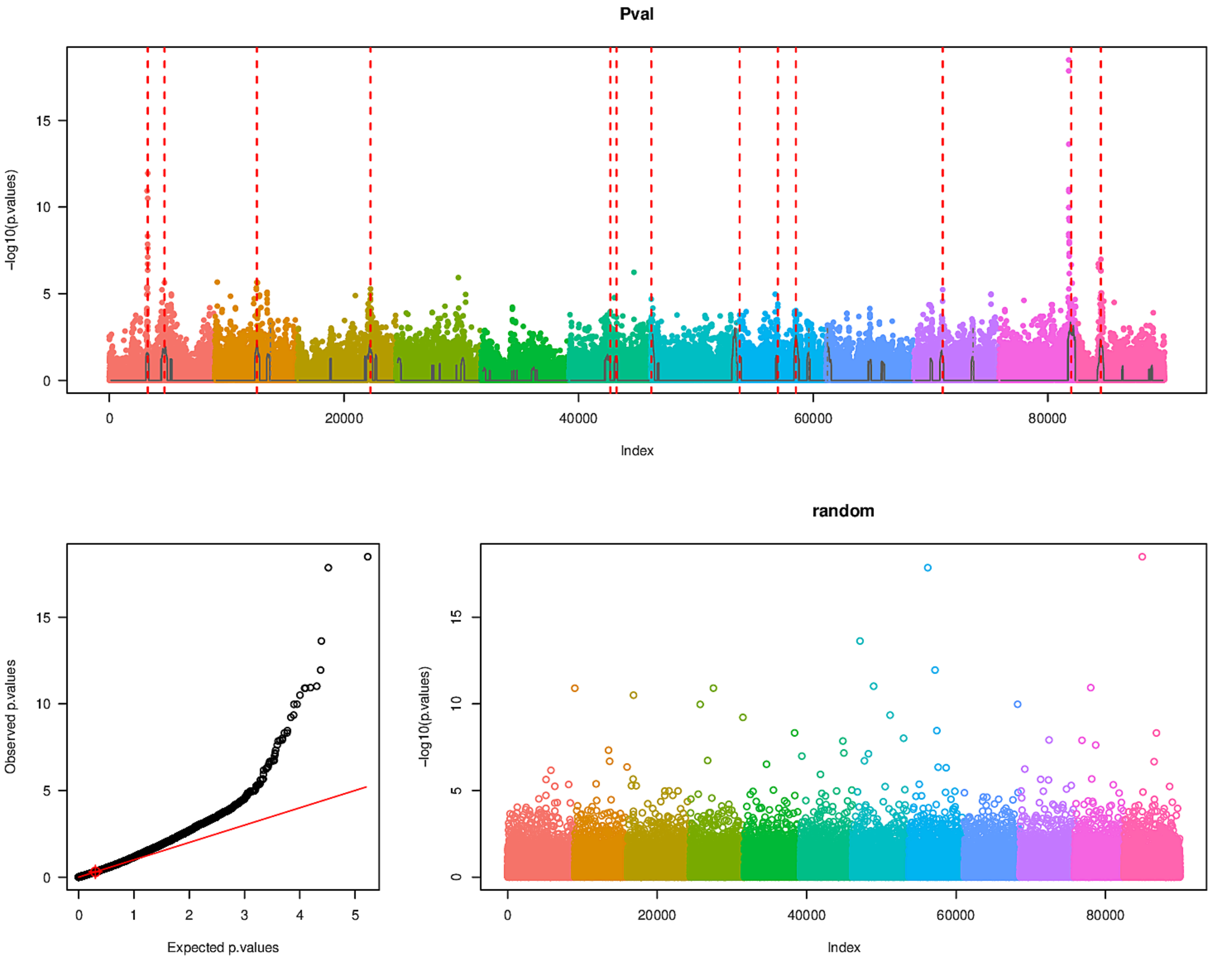
Pval
-log10(p.values)
Index
Observed p.values
Expected p.values
random
-log10(p.values)
Index

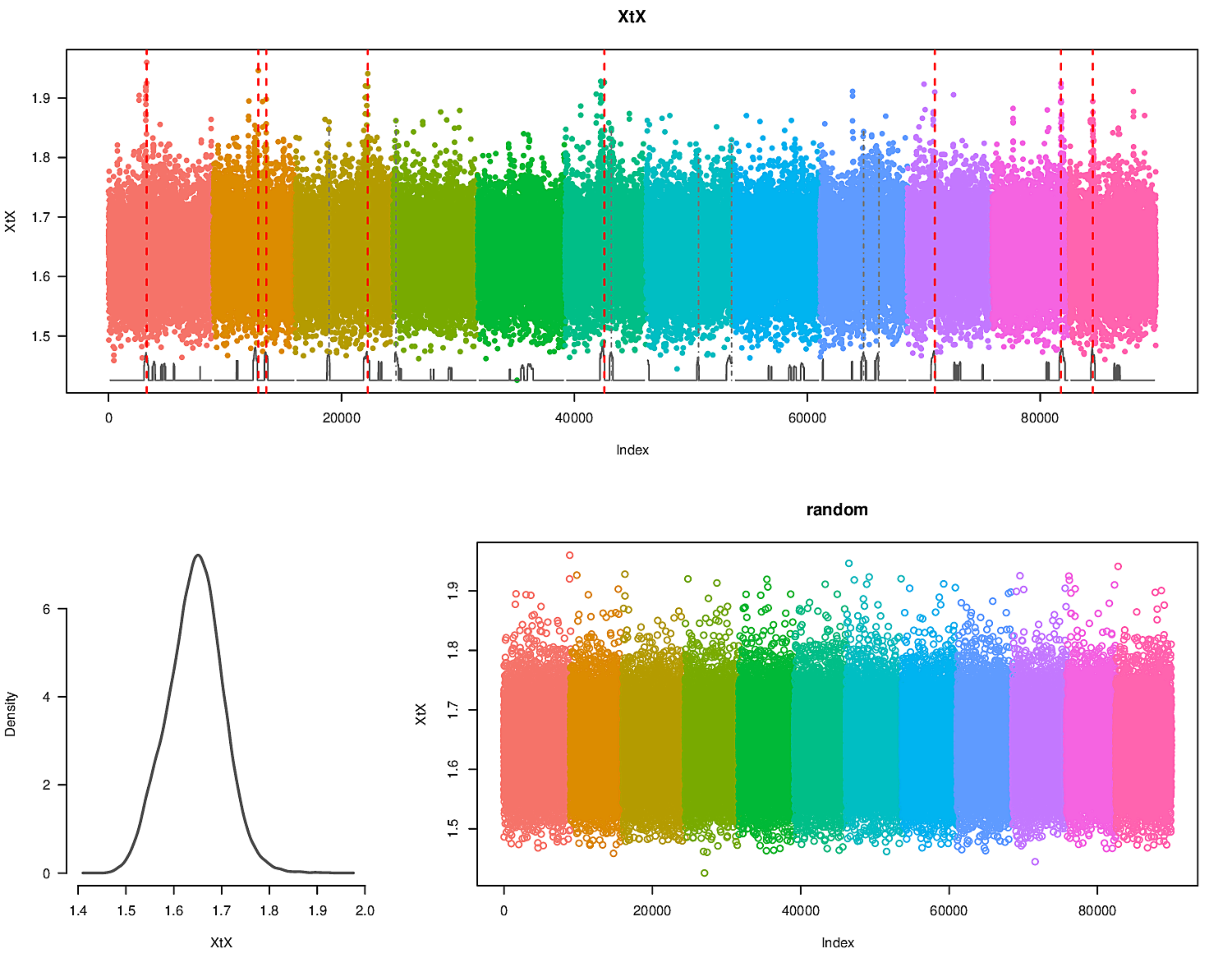
XtX
1.5
1.6
1.7
1.8
1.9
0
20000
40000
60000
80000
Index
Density
0
2
4
6
1.4
1.5
1.6
1.7
1.8
1.9
2.0
random
